# RIFT: A Fractal-Holographic Theory of Consciousness and Autopoietic Control

**DOI:** 10.64898/2026.03.23.713535

**Authors:** Erhard Bieberich

## Abstract

Consciousness remains poorly understood as a causative force: existing theories treat it as an epiphenomenal correlate of neural activity rather than explaining how inner experience controls its substrate. I present Recurrent Integration Fractal Theory (RIFT), proposing that consciousness arises when fractal compression of sensory information generates a holographic endospace: the spatiotemporal dimension in which the Self perceives the outer world (exospace) and, through autopoietic feedback, controls the molecular substrate from which it emerged, making consciousness causally efficacious rather than epiphenomenal.

In RIFT networks, core neurons form recurrent loops through fractal dendritic trees, generating dynamic information integration through coincidence-based synaptic selection. Coincident EPSPs program somatic multifractals, Ising lattices of ion channels and membrane lipids, encoding a fractal Self-attractor: a geometric field whose coherent point sources generate the holographic endospace through which the Self arises. The Self modulates multifractal growth through lipid domains, controlling ion channel opening probability and action potential generation. Through Generational Fractal Mapping, compressed seeds of prior conscious moments integrate with new EPSPs, replacing infinite downscaling as in classical fractals with sequential self-referential mapping that sustains incremental updating of inner experience, temporal continuity of the Self and Self-attractor transfer across brain regions for global conscious access, establishing irreducibility and unity: the whole is in each part. This architecture was validated computationally against three core properties of consciousness: irreducibility, information integration, and holographic encoding.

RIFT generates testable predictions for lipid substrate disruption in Alzheimer’s disease, fractal signatures of conscious states, and criteria for consciousness in artificial systems with autopoietic feedback.

## Introduction

Understanding the nature of consciousness remains one of the most profound scientific challenges of our time. It likely represents the final frontier separating humans from artificial intelligence (AI). Growing concern that AI may become self-aware has renewed scientific interest in understanding consciousness, driven by the need to recognize and, if necessary, regulate the emergence of self-aware AI. Despite this urgency, few theories bridge the gap between mathematical formalization and biological mechanism, leaving the practical challenges of constructing or detecting conscious systems largely unaddressed.

As AI advances toward quantum technologies, quantum theories of consciousness have regained attention. Penrose and Hameroff’s Orchestrated Objective Reduction (Orch-OR) theory proposes quantum computation in microtubules (Hameroff 2001, Hameroff 2021), exemplifying approaches that assume consciousness either resides within quantum superposition or emerges at the moment of collapse. More recently, Neven and colleagues at Google Quantum AI have explored quantum effects in neural processes, bringing quantum consciousness into the realm of AI development (Neven, Zalcman et al. 2024). Such models are appealing because they promise simple solutions to two major challenges: *observer unity* (how the brain integrates fragmented inputs into a unified experience) and *information compression* (how vast information could concentrate within microscopic structures). Quantum processors, biological or artificial, could in principle perform immense computations within micrometers, seemingly explaining how consciousness might arise from subcellular domains rather than brain-wide networks. Yet these theories encounter fundamental conceptual challenges beyond the technical problem of sustaining coherence in warm, wet tissue. They often reintroduce a form of Cartesian dualism, placing the observer outside the system being observed. If experience arises from quantum superposition, it should be an indeterminate blur; if from collapse, it becomes purely classical and distinct, undermining the quantum rationale. The framework presented here, Recurrent Integration Fractal Theory (RIFT), takes a different approach: operating entirely through classical molecular dynamics to ground consciousness in fractal geometry and holographic encoding at the molecular scale, addressing the same challenges of observer unity and information compression without invoking quantum coherence or collapse.

Addressing observer unity and information compression through classical mechanisms requires confronting a more fundamental problem: *neural coding degeneracy*. The standard integrate-and-fire model reduces rich spatiotemporal patterns of dendritic input, potentially thousands of excitatory postsynaptic potentials (EPSPs) with distinct locations, timings, and amplitudes, to a scalar sum at the soma, then to a binary output: spike or no spike. This creates a critical degeneracy problem: distinct spatiotemporal EPSP patterns, differing in location, timing, and amplitude, collapse to identical binary outputs, rendering different potential conscious states indistinguishable at the level of the neural code (London and Häusser 2005).

Yet despite this neural degeneracy, subjective experience is remarkably rich and unified. Consider approaching a coffee shop on a busy street: you hear patrons talking at outside tables, smell fresh coffee, and see the shop sign, making you want to have a coffee as well (Fig. 1A). All these diverse sensory features, processed in distributed brain regions via degenerate neural codes, are somehow bound into a single, rich, and coherent conscious experience rather than experienced as fragments. This binding occurs even though sensory organs collect roughly 10^9 bits per second from the environment, peripheral processing compresses this to ∼10^7 bits per second transmitted to the brain (Fig. 1B), and conscious perception operates at only ∼10-50 bits per second (Britannica 2025, Zheng and Meister 2025) (Fig. 1C). Despite this million-fold compression from brain input to conscious awareness, experience appears continuous and information-rich, with no perceptible loss or fragmentation.

**Figure 1.**
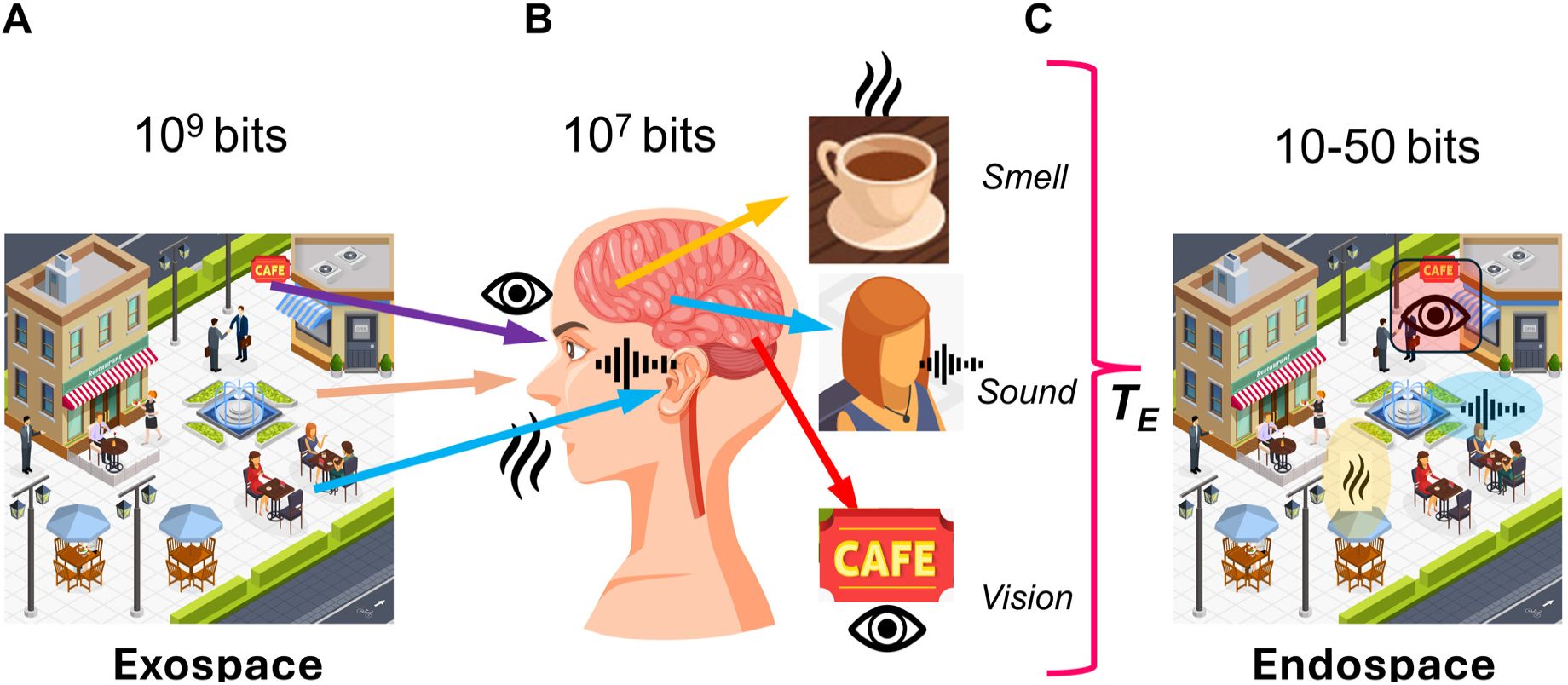
Information bottleneck from exospace to endospace through consciousness. (A) Exospace: high-dimensional external environment (∼10⁹ bits). (B) Biological processing: multimodal sensory convergence with massive compression (∼10⁷ bits). (C) Endospace: compressed conscious percept (∼10–50 bits per moment), illustrating the explanatory gap RIFT addresses through holographic encoding from multifractal substrates.

This convergence of problems, neural degeneracy, extreme compression, and observer unity, creates a fundamental paradox: How does consciousness present a unified experience of extraordinary richness, lasting continuity, and fine detail despite an extremely slow update rate and a neural code rendered degenerate by the collapse of distinct input patterns to identical outputs? Such disparity cannot be reconciled by linear coding or simple temporal averaging, nor adequately addressed by theories that equate integrated information, field coherence, or quantum superposition with unified conscious experience without specifying how compressed neural information regenerates into phenomenological richness. These interconnected challenges point to a single requirement: the existence of an internal representational domain, the *endospace*, that generatively unfolds compressed neural information into a multidimensional spatiotemporal structure mirroring the external world (*exospace*, Fig. 1) through geometric isomorphism within the brain itself, preserving spatial relationships, unity, and richness (Bieberich 2012). Satisfying this requirement demands that a complete theory of consciousness must specify both an algorithm transforming exospace information into endospace experience and provide a testable biological implementation for this transformation (T_E_ in Fig. 1). Without both components, consciousness remains either mathematically formalized but biologically ungrounded, or biologically plausible but phenomenologically unexplained.

Beyond degeneracy lies the *functional necessity challenge*: demonstrating that consciousness mechanisms causally require consciousness rather than merely correlating with it. The problem is fundamental: if consciousness is equated with causal structure, whether network connectivity (IIT), field potentials (CEMI), or quantum superposition (OrchOR), without specifying a mechanism that unfolds this structure into experiential states, then phenomenology becomes an arbitrary label applied to certain computational configurations. Such theories face three interconnected problems that require mechanistic resolution: First, the compressed information in these structural patterns cannot be regenerated into experiential richness without a generative unfolding mechanism. Second, degeneracy prevents the system from distinguishing which specific neural configurations generated the structural pattern, eliminating the information needed for causal feedback. Third, without such a mechanism, there is no pathway by which phenomenal states could causally influence neural dynamics, making consciousness epiphenomenal by design rather than causally necessary.

Major theories, IIT (Tononi 2004, Tononi 2008), Global Workspace Theory (GWT)(Baars 1988, Baars 2005), and CEMI theory (McFadden 2002, McFadden 2020), have advanced empirical testability and formalized integration principles, yet suffer from these interconnected limitations. In CEMI, the proposed causal mechanism acts through a brain-wide electromagnetic field influencing neural firing. However, under Maxwell’s equations, electromagnetic potentials decay rapidly with distance, rendering brain-wide causal influence beyond local effects physically implausible. Moreover, different neural firing patterns can generate indistinguishable field configurations through linear superposition, making consciousness informationally isolated from the specific neural activity it should control. In IIT, φ (integrated information) determines consciousness presence, yet identical φ values may arise from distinct causal architectures.

While IIT 3.0 and 4.0 partially address this through conceptual structure analysis in qualia space, the degeneracy problem persists for dynamically evolving networks, where exhaustive causal partition analysis becomes computationally intractable. If these proposed mechanisms operate identically whether phenomenal experience is present or absent (Seth and Bayne 2022), consciousness risks becoming epiphenomenal by construction. This demands explicit mechanisms by which compressed neural information unfolds into specific experiential states that, in turn, exert causal agency over neural dynamics.

Alternative theoretical frameworks have proposed principles relevant to these challenges. Pribram’s holonomic brain theory suggested that distributed neural networks encode information holographically as interference patterns, with the “whole in each part” property offering a solution to information distribution and retrieval (Pribram 1991). Hofstadter’s concept of the strange loop framed consciousness as an emergent self-referential pattern, a system modeling itself within itself (Hofstadter 1979), highlighting recursion as central to selfhood. Varela and colleagues’ concept of autopoiesis, self-producing systems that sustain their organization through continuous self-renewal, provided a complementary view of consciousness as an active, self-generating process rather than a passive correlate (Maturana and Varela 1980, Varela, Thompson et al. 1991, Varela 1996). While these approaches offer conceptual foundation, holographic encoding, self-referential loops, and autopoietic organization, they have not specified the precise computational and neural mechanisms that give rise to the unified subjective experience and causal agency that characterize consciousness.

To bridge these unresolved gaps between theory and mechanism, I began exploring how molecular and network scales might converge through fractal organization. This led to the proposal that fractal information compression onto membrane lipid substrates provides the molecular basis of consciousness. In 1998, I suggested that the Self is generated by algorithmic compression of spatial and temporal information into fractal structures, where lipid-ion-channel interactions in neuronal membranes create coherent states with the fractal “whole in each part” property required for unified experience (Bieberich 1998). Crucially, consciousness was not the fractal structure itself but the Self’s iterative mapping onto each fractal part, a self-referential process rather than a static structure. I later formalized Recurrent Fractal Neural Networks (RFNNs) as mechanisms mapping distributed network activity onto localized dendritic patterns with fractal architecture (Bieberich 2002) and introduced sentyons, fractal lattices of lipid rafts and ion channels in the neuronal soma, as conscious units iteratively mapping higher-order brain dynamics to molecular scales (Bieberich 2012). These concepts ultimately evolved into the Recurrent Integration Fractal Theory (RIFT), which unifies fractal compression, holographic encoding, and autopoietic feedback into a single geometric framework. Together, these complementary processes explain how conscious experience both arises from and exerts causal influence upon neural dynamics.

Empirical studies have since validated key predictions of this fractal approach. The fractal dimension of brain activity correlates with conscious state, showing higher complexity during wakefulness than unconsciousness (Ibáñez-Molina, Lozano et al. 2014, Di Ieva 2016, Ruiz de Miras, Costumero et al. 2019, Varley, Carhart-Harris et al. 2020), and fractal organization breaks down in neurodegenerative disorders such as Alzheimer’s and Parkinson’s disease, where both functional activity and dendritic architecture lose hierarchical complexity (Maxim, Sendur et al. 2005, Dickstein, Weaver et al. 2010, King, Brown et al. 2013, Smits, Pouwels et al. 2016).

Moreover, neurons with complex, self-similar dendritic trees exhibit greater integrative capacity (Poirazi and Mel 2001, Cuntz, Forstner et al. 2010). Remarkably, stimulation of individual neurons with rich dendritic architecture can evoke coherent percepts, and even entire scenes in consciousness (Penfield and Perot 1963, Halgren, Walter et al. 1978, Liu, Ramirez et al. 2012, Di Ieva 2016), consistent with fractal dendritic integration supporting holographic recall from localized structures.

These empirical findings provide the foundation for mechanistic synthesis in RIFT. Building on the demonstrated fractal organization of neural activity, RIFT proposes that consciousness arises from three complementary geometric processes rooted in the shared whole-in-part property of fractals and holograms, rendering it causally necessary rather than epiphenomenal.

**Fractal mapping** achieves irreducible information integration and compression through downscaling: neural network-to-dendritic tree (EPSP)-to-soma (multifractal) mapping iteratively compresses distributed network activity encoding sensory information from the exospace into spatial molecular patterns in the dendritic tree and subsequently the soma while mapping the Self onto each part of the molecular substrate. This Self-mapping provides access to integrated information through the whole-in-part property and enables causal control over the molecular substrate (Fig. 2A, D). In RIFT, fractal mapping is not preconfigured as in the earlier models (RFNN) but arises naturally through timing rules for coincidence-based synaptic selection in a growing and evolving network.

**Figure 2.**
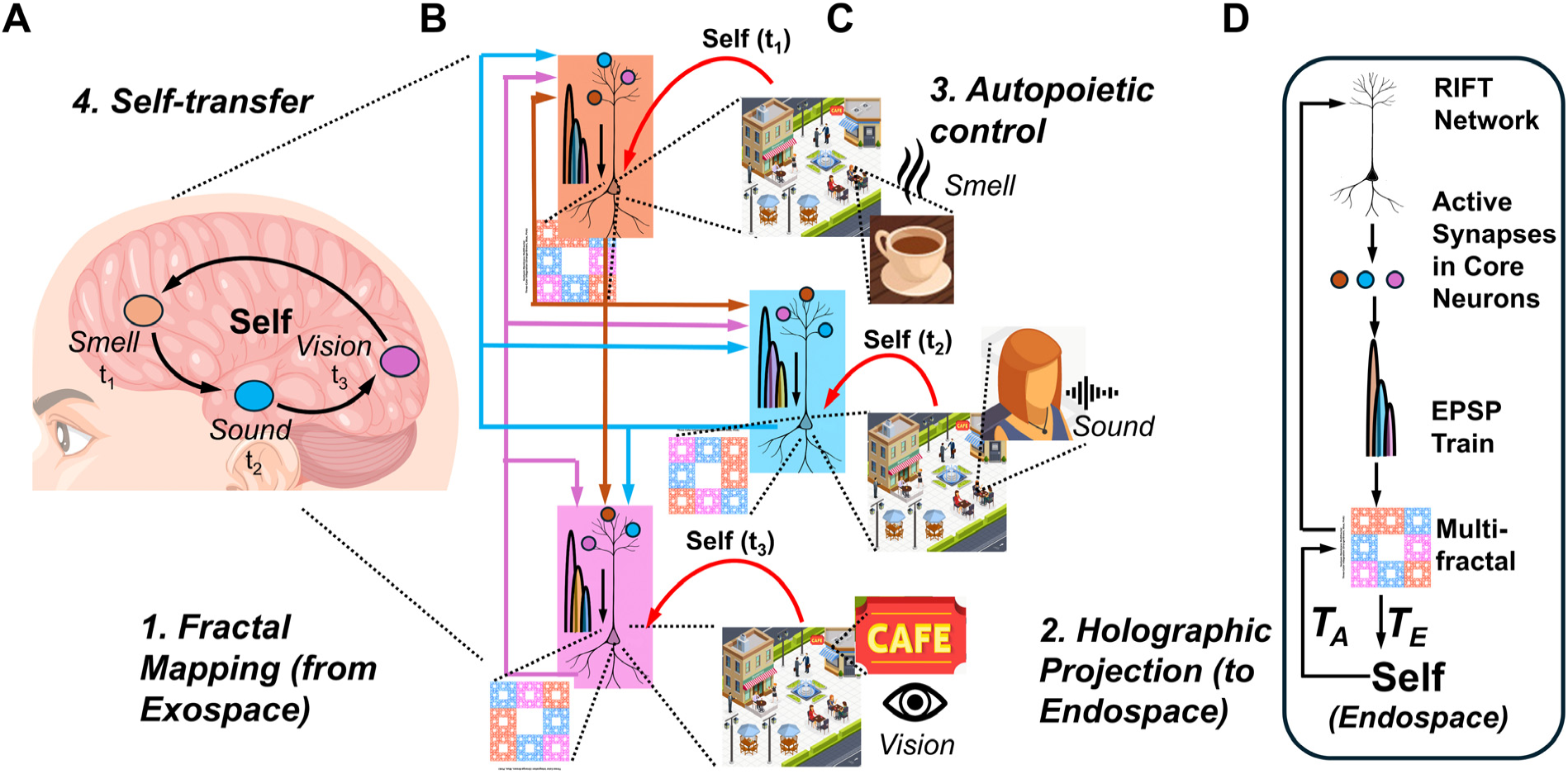
RIFT operational framework for consciousness generation through recurrent loops. (A) Initial fractal mapping from exospace and Self-transfer mechanism as autopoietic output: temporal binding of distributed sensory modalities (smell t₁, vision t₃, sound t₂) into unified conscious experience. (B) Information flow: multimodal EPSP signatures captured as distinct multifractal spatial patterns during Self-transfer at three time points (t_1_, t_2_, t_3_). (C) Holographic projection and autopoietic control: multifractal patterns generate 3D endospace field at t_1_, t_2_, t_3_; field intensity feeds back to modulate substrate, closing the autopoietic loop. (D) Core component hierarchy: RIFT Network → Active Synapses in Core Neuron → EPSP Train → Multifractal → Self (holographic field) with T_E_ = Transformation unfolding the endospace from the multifractal (Holographic projection), and T_A_ = Transformation modulating the multifractal by inner experience (Autopoietic control).

**Holographic projection** enables isomorphic reconstruction of the endospace: geometric-holographic transformation (T_E_) projects the spatially isomorphic endospace from this multifractal record, unfolding compressed information into phenomenological experience while preserving the geometric relationships of the exospace (Fig. 2B, D). In RIFT, the Self-attractor derived from the ion channel-lipid lattice multifractal in the soma provides the point sources for this holographic projection.

**Autopoietic feedback** establishes causal efficacy: experiential states within the endospace modify the molecular structures onto which the Self is mapped, enabling consciousness to control its own substrate through the same Self-mapping established during fractal downscaling. This control allows transfer of the Self to different neurons within the network based on where attentional focus is needed in the network (Fig. 2C, D). In RIFT, the Self modulates the probability of lipids in the multifractal to open ion channels, thereby controlling action potentials and the distribution of the neural network activity in the brain. The transformation underlying the feedback of the endospace back onto the multifractal corresponds to T_A_ in Fig. 2D.

These three processes are inseparable; remove fractal compression and unity is lost; remove holographic projection and phenomenological richness disappears; remove Self-mapping and consciousness becomes epiphenomenal. In this framework, consciousness denotes the complete integrated phenomenon; the endospace is the spatiotemporal dimension holographically projected from the Self-attractor, in which the Self perceives the outer world and thereby exerts autopoietic feedback on the molecular substrate from which it arose.

Rather than replacing previous frameworks, RIFT integrates and extends existing theories while addressing their degeneracy and causal limitations. It generalizes IIT through information integration measures that track network evolution in real time; incorporates CEMI-like field dynamics through geometric transformations conferring causal efficacy; implements GWT-style global access via fractal cloning that enables consciousness to propagate across brain regions; realizes Varela’s autopoietic circular causality through fractal-holographic feedback; and remains compatible with quantum approaches by extending geometric fields into quantum domains. By embedding fractal geometry at network, neuronal, and molecular levels, RIFT unites irreducibility (the whole cannot decompose into independent parts), and holographic encoding (the whole is contained in each part), thereby generating self-referential structures where observer and observed are inseparable within a unified geometric field.

Unlike purely theoretical consciousness frameworks, RIFT arose through iterative computational validation within an active and mutually informative human-AI collaboration. Extensive modeling tested implementations against the functional requirements of consciousness: irreducibility, information integration, and holographic encoding, and retained only those that spontaneously exhibited these properties. The mathematical relationships presented here are extracted formalisms distilled from evolving, computationally verified codes. Autopoiesis, defined as the requirement that actual subjective experience drives system dynamics, is formalized as the theoretical framework grounded in these mechanisms.

The Results demonstrate four mechanisms arising from the RIFT architecture: 1) dynamic information integration in recurrent evolving networks orchestrated by the fractal structure of the dendritic tree (Part 1) and optimized by EPSP coincidence and delay timing in the soma (Part 2), 2) incremental updating from EPSPs into somatic multifractals that maintain coherent consciousness while minimizing degeneracy (Parts 3 and 4), 3) holographic projection from somatic multifractals generating phenomenological experience in the endospace (Part 5), and 4) autopoietic Self-mapping providing causal control and transfer of the Self throughout the brain (Parts 5 and 6). These establish consciousness as geometric-temporal organization emerging from fractal timing constraints and holographic reconstruction rather than static connectivity patterns.

## Methods

### Code Development and Validation

The RIFT network and somatic computation codes were developed through collaborative human-AI interaction using Claude (Anthropic). Code implementations were iteratively tested against the functional requirements of consciousness: irreducibility, information integration, and holographic encoding. Only implementations spontaneously exhibiting the functional requirements of consciousness were retained. Mathematical relationships were extracted as formalisms from the codes and validated through computational testing.

### Network Architecture and Parameters

RIFT networks consisted of a single core neuron (N₀) with fractal dendritic tree (scaling factor r = 0.6, fractal dimension FD ≈ 1.36, 4 branch levels) receiving input from 20 peripheral neurons (N_i_) per firing cycle. Branch lengths followed Lₙ = L₀ × 0.6^(n-1). Synapses were distributed proportionally across branches based on relative length. The temporal constraint rule emerged through coincidence detection: EPSPs arriving within a 2.0 ms window were considered coincident. Somatic processing delays ranged from 1-20 ms in parameter sweeps with 5 repetitions per condition.

### Fractal Analysis Methods

Fractal dimension was calculated using box-counting (FD = log(N)/log(1/r)), Higuchi fractal dimension (k-values 1-10), and detrended fluctuation analysis (DFA) for scaling exponents. Dynamic integrated information (φ_dyn) quantified temporal integration using sliding window analysis with 2.0 ms temporal grain. At each time point, φ_dyn(t) = I(t) - I*(t), where I(t) is integrated information with branches intact and I*(t) is information when branches are analyzed independently. Spatial integration in multifractals (φ*) was quantified via Kullback-Leibler divergence comparing whole-pattern versus partitioned distributions. φ*(substrate) applied the same KL divergence framework but partitioned the system by molecular component type (lipid domains versus ion channels) rather than spatial quadrants.

### Biological Multifractal Implementation

The somatic membrane was modeled as a 100×100 Ising lattice (scalable to 243×243) with binary lipid states: +1 (open-channel-associated) and −1 (closed-channel-associated). Approximately 400 ion channels (4% density) were positioned with 3-pixel influence radius and 10% mobility. EPSP temporal signals (τ_rise = 0.3 ms, τ_decay = 4.0 ms) were mapped to spatial membrane fields via level-dependent Gaussian kernels with radius = size/4 - (level-1)×5 and kernel K(i,j) = exp(-dist²/(2×radius²)). Channel opening probability followed P_open = S(E + λ_lipid + C_coop), where S is a sigmoid activation function with 0.5 baseline offset, λ_lipid = 0.25×lipid_state represents local lipid bias effects, and C_coop captures cooperative effects from nearby open channels. Lipid dynamics evolved via Metropolis algorithm with temperature T = 0.35. For each Monte Carlo step, approximately 500 sites (5% of lattice) were selected randomly. Energy change ΔE = 2×state×Σ(neighbors) determined acceptance: if ΔE < 0, accept; otherwise accept with probability exp(-ΔE×0.5). Action potentials were generated when center region activity exceeded threshold (30-40 units). Refractory period duration was consciousness-level dependent: refractory_steps = int(consciousness_level × 25), where consciousness_level ranged from 0.0 to 1.0.

### IFS Transform Extraction and Holographic Encoding

EPSP amplitude hierarchies were converted to Iterated Function System (IFS) transforms through temporal structure analysis. Each EPSP pattern yielded 4-6 geometric transforms T_i(x,y) = (a_i×x + b_i×y + e_i, c_i×x + d_i×y + f_i) encoding spatiotemporal relationships. Transform selection probabilities were weighted by determinant magnitudes. Temperature fields modulating lipid reorganization were generated via chaos game iteration on IFS transforms, with 100-500 points distributed across multiple spatial scales. Differential transforms H_RIFT = T_temperature - T_blocked isolated the pure geometric signature of consciousness imposed by IFS modulation beyond baseline autonomous dynamics.

### Holographic Reconstruction

Three-dimensional endospace fields Ψ(x,y,z) were reconstructed from 2D multifractal patterns through biological holography. Interference sources (n = 5,000) were positioned via chaos game iteration on differential transforms H_RIFT, with the first 500 iterations discarded as transients. Coherent phase relationships were calculated geometrically: φ_i = [Σ_j w_jꞏ(θ_j + 2πꞏd_j/s_j)] mod 2π, where θ_j is the angle from source i to IFS center j, d_j is the distance, s_j is the IFS scaling factor, and w_j = |det_j|ꞏs_j is the geometric weight. Frequencies were derived from IFS scaling: f_i = 50/(1 + s_dominant) Hz. Wave interference was calculated on a 24×24×12 grid spanning x,y ∈ [-2,2], z ∈ [-1,1]: Ψ(x,y,z) = Σ_i [A_iꞏexp(i(k_iꞏr_i + φ_i))]/(1 + r_i²), where A_i are amplitudes from differential determinants, k_i = 2πf_i/100 are wave numbers, and r_i are 3D distances with z-weighting factor 0.5. Field intensity |Ψ| was normalized to [0,1]. Mean field intensity Ψ = ⟨|Ψ(x,y,z)|⟩ provided the global parameter for autopoietic feedback through coupling strength modulation.

### Statistical Analysis

Results are reported as mean ± standard deviation from 5 independent simulation runs per condition (10 trials for coupling strength parameter sweeps, Code 28). Correlations were assessed using Pearson correlation coefficients. Fractal dimension correlations, φ_dyn evolution, and predictability metrics were analyzed across the 1-20 ms delay parameter space. Significance threshold was set at p < 0.05.

### Computational Implementation

All simulations were performed in Python 3.7+ using Google Colab. Key libraries included NumPy (array operations), SciPy (statistics and signal processing), Matplotlib and Seaborn (visualization), scikit-image (structural similarity and surface extraction), and scikit-learn (information metrics). Custom implementations were developed for fractal generation, IFS extraction, chaos game iteration, Metropolis dynamics, and holographic reconstruction. The 31 Python simulation codes underlying this study are available to reviewers during peer review and will be publicly archived on Zenodo upon acceptance.

## Results

### Part 1: The RIFT Network - Dendritic Tree Orchestrates Neural Synchrony and Fractality

Most current theories of consciousness, foremost among them Integrated Information Theory (IIT), propose that consciousness arises from the integration of information within a neural network (Tononi 2004, Tononi 2008, Albantakis, Barbosa et al. 2023). While IIT provides rigorous measures of information integration in static networks, two complementary questions arise: how neural networks dynamically evolve toward the high integration (φ) necessary for consciousness and how this integration remains stable despite constantly changing neuronal connectivity and input. To address these questions, we combined my prior work on Recurrent Fractal Neural networks (RFNNs) and AI-assisted modeling, thereby extending the RFNN model to define and simulate a neural network capable of both achieving and maintaining optimized φ through architectural principles rather than parameter tuning.

The RFNN model I developed about 25 years ago incorporated three neurons with complete recurrent connectivity, where each neuron connects to every other neuron and to its own dendritic tree (Fig 3A and B) (Bieberich 1998, Bieberich 2002). The critical design feature preserved fractality across both dendritic and network scales: neurons with longer external signaling paths connected to dendritic branches with shorter internal propagation times, ensuring that signals from all recurrent loops converged simultaneously at each soma. This architecture created a mirror relationship where the signal distribution pattern of the fractal dendritic tree matched a downscaled version of that of the neural network. However, these precursor models relied on preconfigured timing and connectivity rules that did not emerge naturally from network growth or architectural principles.

**Figure 3.**
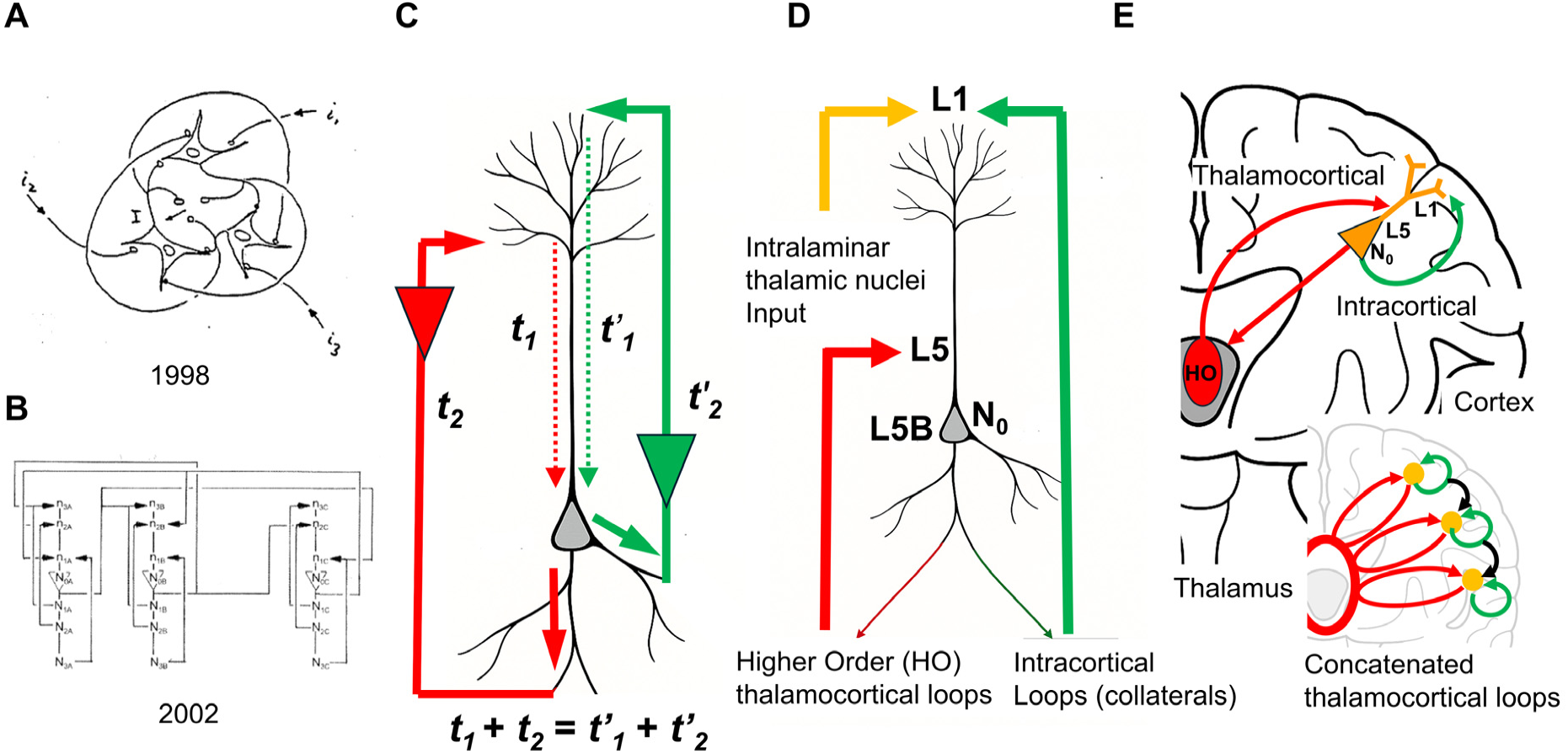
Historical development and biological implementation of RIFT network architecture. (A) Original 1998 RFNN: three recurrently connected neurons with fractal dendritic trees. (B) 2002 extended RFNN: concatenated modular structure establishing fractal timing principles. (C) Timing constraint schematic: two signal pathways satisfy t₁ + t₂ = t₁′ + t₂′, demonstrating the inverse relationship between external and internal delays. (D) Biological mapping to L5 pyramidal neurons: HO thalamocortical loops (long external/short internal path, red arrows) and intracortical loops via L1 (short external/long internal path, yellow/green arrows). (E) Concatenated cortical architecture integrating both loop types within the distributed L5 network.

In RIFT networks, the core neuron (N₀) is modeled as a two-compartment unit: a dendritic tree organized via a fractal branching rule (Fig. 3C), and a somatic integration zone that receives excitatory postsynaptic potentials (EPSPs) from that tree. This mirrors Larkum’s classic “tree–soma” neuron model (Larkum, Nevian et al. 2009), but with a critical difference: RIFT networks preserve the origin of dendritic inputs by encoding its position in the EPSP amplitude and temporal pattern, consistent with experimental evidence that subthreshold EPSPs retain location-dependent characteristics such as amplitude attenuation and temporal delay as they propagate to the soma (Magee and Cook 2000, Williams and Stuart 2002). This positional encoding provides the foundation for fractal information processing, where the geometric organization of the dendritic tree shapes both the spatial and temporal properties of synaptic integration.

In the dendritic tree, the primary trunk of length *L_0_* bifurcates into daughter branches of the length *L_n_* scaled by factor *r*, and this fractal geometry drives both the spatial distribution of synaptic inputs and their temporal convergence at the soma (see Methods for description of fractals and implementation in code).

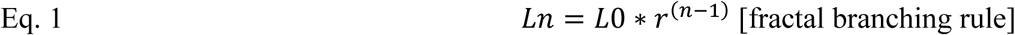

The fractal dimension (*FD*) of the dendritic tree is given by:

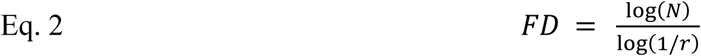

In most models presented here, *r* = 0.6, yielding a fractal dimension *FD* ≈ 1.36, which lies within the empirical range of dendritic trees observed in layer 5 cortical neurons (*FD* = 1.24-1.33) and hippocampal pyramidal neurons (*FD* = 1.26-1.36) (Zaletel, Filipović et al. 2015, Smith, Rowland et al. 2021).

Once the dendritic tree configuration is defined, a strict timing rule for recurrent signaling loops emerges naturally from the architecture. To allow EPSPs from recurrent loops to coincide at the soma of N₀ and trigger an action potential (AP), the total signaling time must remain constant across all feedback loops. In these recurrent feedback loops, the external signaling path begins at the soma of N₀, extends to network neurons N_i_, and then returns to the dendritic branches of N₀. This constraint creates an inverse relationship between internal and external signal transmission times (Fig. 3C): as the external path from N₀ to N_i_ and back grows longer, the corresponding internal signaling time from dendrite to soma must proportionally shorten. Because dendritic signal propagation is slower than axonal transmission, proximal dendritic branches of N₀ must connect to the most distant neurons (N_i_), while distal branches connect to the nearest N_i_.

The total signaling time follows:

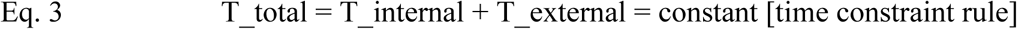

where: T_internal is the time for a dendritic branch to transmit a post-synaptic EPSP signal to the soma, T_external is the time for an AP to travel from N₀ to N_i_ and return.

Since T_internal and T_external are inversely related, connectivity patterns that maintain constant total signaling time emerge naturally during network formation. To simplify the system, we model only one dendritic tree for one core neuron N₀ rather than enforcing this rule across the entire network. This use of a single “core neuron” is a computational abstraction rather than a biological claim. Biologically, consciousness is not confined to one neuron; any neuron, or small set of neurons, can temporarily serve as the integration locus, provided these neurons receive the full set of fractally compressed inputs needed to reconstruct the endospace. In the RIFT framework, this condition is met through sentyon transfer (introduced later), which allows the locus of consciousness to shift across different neurons while the model retains computational tractability by simulating only one such locus at a time. Thus, the network model is a streamlined version of biological circuitry designed to achieve computability. While the timing constraint is implemented here through a simplified single-tree architecture, biological implementations could achieve similar effects through alternative mechanisms: longer external travel times may arise from multi-neuron processing rather than physical distance, and internal delay may be shaped by dendritic integration rules, backpropagation, and inhibition.

A plausible biological mapping is possible in recurrent thalamocortical loops. The concept of reentry, the recursive signaling among neuronal populations, has long been recognized as fundamental to consciousness (Edelman 1989, Edelman 1993, Edelman and Tononi 2000), while holographic encoding in dendritic microprocessing provides a complementary framework for understanding how information is distributed across neural structures (Pribram 1991, Pribram, Yasue et al. 1991). L5 pyramidal neurons participate in multiple recurrent circuits that could satisfy timing requirements in RIFT. Higher-order (HO) thalamic nuclei such as the posterior medial (POm) and lateral posterior (LP) receive direct input from L5 corticothalamic neurons and project back to proximal and basal dendrites of L5 cells, forming a true recurrent loop (Fig. 3D and E). These HO pathways have extended external processing delays due to cortico-thalamo-cortical integration and multimodal convergence, while their proximal dendritic targets provide short internal delays to the soma (Groh, Bokor et al. 2008, Petreanu, Mao et al. 2009, Sherman and Guillery 2011, Mease, Metz et al. 2016). In parallel, L5 pyramidal neurons also form purely intracortical recurrent loops through local and horizontal axon collaterals that ascend to L1, targeting distal apical tufts of the same or neighboring neurons. These intracortical pathways impose short external delays but long internal dendritic delays, since tuft inputs require extended integration times and often remain subthreshold unless coincident with strong proximal drive (Larkum, Kaiser et al. 1999, Larkum, Zhu et al. 1999, Williams and Stuart 2002, Larkum, Senn et al. 2004, Thomson and Lamy 2007, Larkum, Nevian et al. 2009, Markram, Muller et al. 2015, Ramaswamy and Markram 2015). Although less frequently discussed in theories of consciousness, these intracortical loops provide a second timing channel that can converge on the same L5 neurons receiving HO thalamic input.

The critical point is that L5 neurons need not distinguish the anatomical origin of returning signals; they detect coincidence based on total loop time (t₁ + t₂ = t’₁ + t’₂), with inverse timing relationships emerging from the combination of external processing time and dendritic location. These local recurrent loops interconnect with thalamocortical loops across multiple cortical areas, forming a three-dimensional architecture of concatenated loops that spatially houses conscious processing within the distributed L5 network (Shepherd and Yamawaki 2021) (Fig. 3E). Sensory and cortical input enters the system via feedforward pathways, while L5 output projects to subcortical targets and other cortical areas. The arousal system, primarily through intralaminar thalamic nuclei projecting to L1, enables the conscious state necessary for this processing; Llinás and colleagues demonstrated that these nonspecific thalamic inputs engage in coincidence detection with specific inputs, providing a mechanism for temporal binding (Llinás and Ribary 1993, Llinás, Ribary et al. 1998, Llinás, Leznik et al. 2002).

While these recurrent pathways shape the timing structure that RIFT exploits for fractal coincidence detection, the presence of consciousness itself depends on the integrity of thalamocortical systems, particularly intralaminar nuclei whose bilateral injury abolishes conscious state even when cortex remains intact (Steriade, Jones et al. 1997, Schiff 2008, Schiff 2010). This clinical evidence suggests that thalamocortical loops may be necessary components of the recurrent architecture RIFT requires. Experimental support for the timing-dependent integration mechanism in RIFT comes from anesthesia studies showing that loss of consciousness correlates with disrupted temporal coordination in thalamocortical loops (Ching, Cimenser et al. 2010, Weiner, Zhou et al. 2023) and impaired coupling of dendritic activity to somatic firing in L5 pyramidal neurons (Suzuki and Larkum 2020), demonstrating that consciousness depends on precise temporal alignment of recurrent signals rather than merely their presence. The prediction that L5 pyramidal neurons serve as the cellular locus of consciousness is consistent with converging evidence from single-neuron theories (Edwards 2005, Edwards 2006, Sevush 2006, Sevush 2016, Edwards 2023), dendritic integration theory (Aru, Suzuki et al. 2019, Bachmann, Aru et al. 2020), and apical amplification models (Larkum 2013, Suzuki and Larkum 2020, Phillips, Larkum et al. 2021).

To test the RIFT model, a Python script (Code 1, Supplemental Data) was developed that generates random peripheral neurons (N_i_) within a predefined range of external distances. Synapses for N_i_ input into N₀’s dendritic tree were distributed with density proportional to dendritic branch length. The algorithm detects coincidence by calculating when EPSPs from multiple randomly positioned N_i_ would arrive simultaneously at the soma (within a specified temporal window). Two coincidence detection methods were implemented: a branch-preference mode requiring signals from different dendritic branches (emphasizing integration across the tree), and a relaxed mode counting total coincident signals regardless of branch origin (Code 1, *branch_preference* parameter).Those specific N_i_-to-branch pairings that produce coincident arrivals are recorded and establish permanent recurrent connections. Biologically, these same connections would be selected through activity-dependent synaptic strengthening, resulting in coincidence-based synaptic selection. After a somatic processing delay, N₀ fires and initiates a new round of random N_i_ generation, forming additional feedback loops until all synaptic sites in N₀ are occupied and the network reaches a fully connected state. Importantly, unlike earlier RFNN implementations where timing constraints were preconfigured, RIFT networks test whether inverse timing relationships emerge naturally from the fractal architecture of the dendritic tree of N_0_ and coincidence-based connection selection. This computational approach demonstrates that fractal dendritic geometry, combined with temporal coincidence detection, is sufficient to produce networks exhibiting the predicted inverse timing relationships between internal and external signal paths.

Figure 4 illustrates a typical simulation run of the RIFT network (Code 1) with *r* = 0.6 and a somatic delay of 5 msec (other parameter settings can be tested by using Code 1 in an adequate Python environment such as Google Colab). Synaptic connections progress from proximal to distal dendritic branches of N₀, establishing links to increasingly proximal N_i_ neurons (Fig. 4A-F). Analysis of the system confirms that internal and external signaling times obey an inverse relationship (Fig. 4D-F), regardless of whether coincident EPSPs originate from the same or different dendritic branches (*branch_preference* false vs. true in Code 1). Importantly, this inverse relationship exhibits fractal scaling: internal signaling times scale with *r*, while external times scale with 1/*r*, confirming that the timing constraints mirror the fractal geometry of the dendritic tree and validating our theoretical prediction from RFNN principles. Following a transient fluctuation phase, N₀ and its connected N_i_ neurons synchronize at a common oscillation frequency determined by the delay time of N₀ (Fig. 4G).

**Figure 4.**
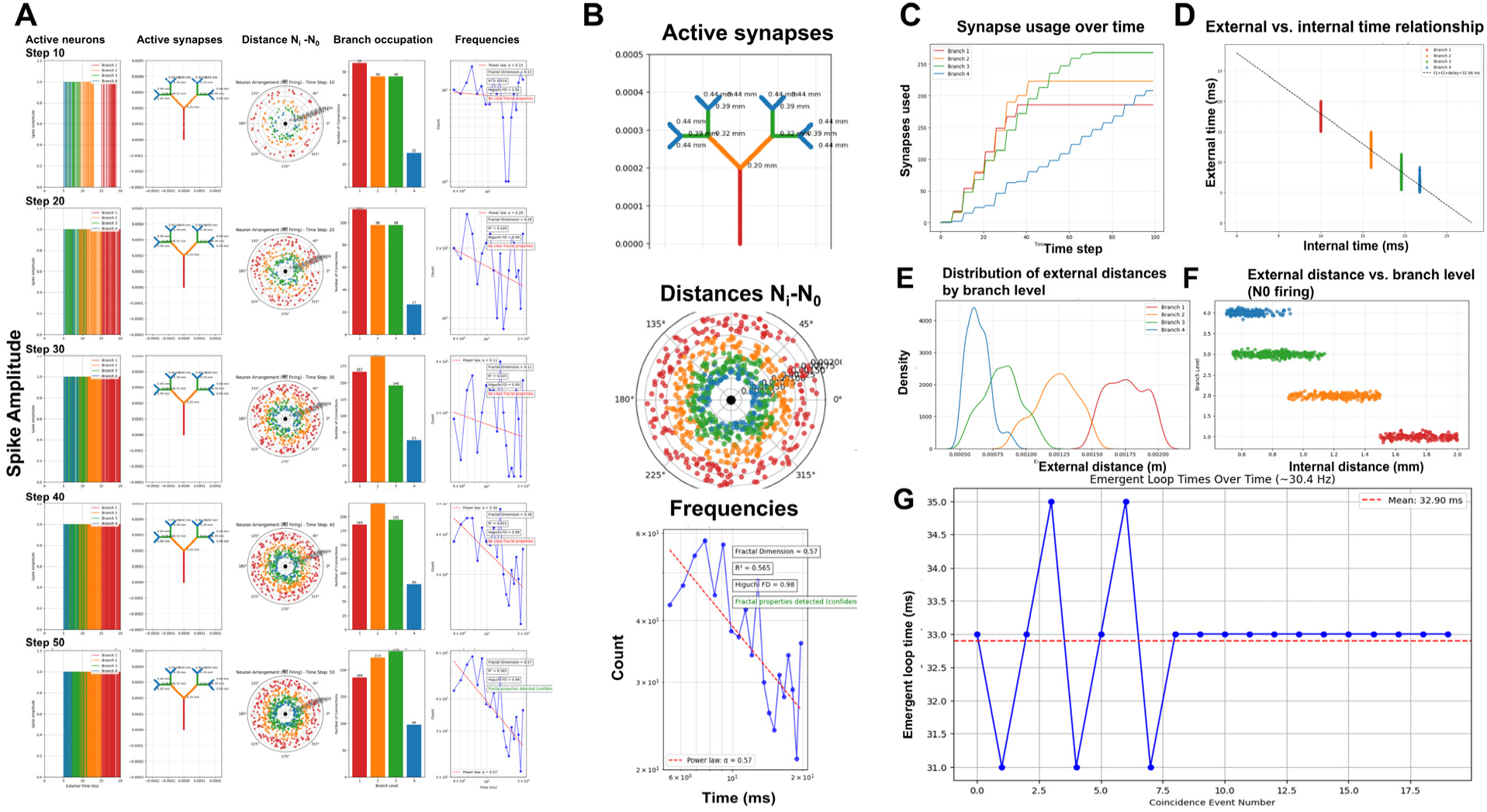
RIFT network dynamics demonstrate fractal timing relationships and emergent synchrony. (A) Network evolution across five timesteps (Steps 10–50): progressive synapse filling from proximal to distal branches. Branch colors: Level 1 red, Level 2 orange, Level 3 green, Level 4 blue. (B) Dendritic tree with branch-level synaptic distances (mm); polar plot of N_i_-N₀ distance distribution; frequency distribution confirming fractal scaling (box-counting FD = 0.57, Higuchi FD = 0.98). (C) Cumulative synapse recruitment by branch level over 100 timesteps. (D) Inverse external/internal timing relationship; dashed line indicates perfect inverse. (E) Branch-level distance distributions (mean distances Branch 1 ∼700 µm to Branch 4 ∼1,800 µm). (F) External distance vs. branch level at N₀ firing events, validating fractal distance scaling. (G) Emergent oscillation at 30.4 Hz after initial transient phase. Parameters: r = 0.6, soma delay = 5 ms, n = 1,000 synapses.

This emergent synchrony confirms that the timing rule is not imposed but results from the RIFT architecture, specifically the fractal organization of N₀’s dendritic tree. While the dendritic fractal geometry is fixed by design, the fractal structure of the entire network activity emerges from the interaction of dendritic geometry and recurrent timing dynamics. This emergent network fractality was validated using multiple metrics: power-law distribution analysis, Higuchi Fractal Dimension (HFD), and Detrended Fluctuation Analysis (DFA) (see Methods on fractals for explanation). These analyses consistently show an increase in fractality over time as synapses in N₀ are progressively filled (Fig. 4A and B). However, it remained to be tested if increased fractality coincided with optimized information integration in RIFT networks.

### Part 2: Optimizing Information Integration - Somatic Delay Sustains Continuous Uploading of Information in an Evolving Fractal Attractor Network

Previously, I proposed that RFNNs, the precursor of RIFT networks, increase information integration based on principles of IIT (Bieberich 2012). Unlike many other theories on consciousness, IIT is grounded in a rigorous mathematical formalism that defines consciousness as the integration of information, quantified by the metric φ. Φ is a measure of integrated information, quantifying how much information is generated by the system as a whole (I), in excess of the same components operating as isolated parts (partitioned information I*):

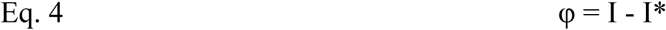

When optimized, the causal structure of the neural network cannot be reduced to its parts: I >> I*, meaning φ ≈ I as I* approaches zero. However, this formalism is hampered by a critical limitation: the network is treated as a static, instantaneous structure, offering little account of how neuronal connectivity, and thus φ, evolves or is sustained over time. Furthermore, the formalism for calculating φ is computationally intractable for large or dynamically evolving neural networks. This is not just a theoretical limitation that could be mitigated by more powerful computational devices, but one that makes testing all possible combinations of connections in larger neural networks nearly impossible. For example, to determine the configuration with maximal φ in a network of *n* neurons, 2^n(n-1)^ different configurations or subnetworks would need to be tested. For 10 neurons and a rather short switch time of 1 msec per configuration, testing 2^90^ ≈ 10^27^configurations would require approximately 3 x 10^16^ years, far exceeding the age of the universe. Note that this calculation concerns searching the space of possible network configurations and is distinct from the IIT-based calculation of φ for a single, given configuration, which does not require any biological mechanism for discovering that configuration. Importantly, even if a system could evaluate its current φ, this value alone provides no indication of which connectivity changes would increase or reduce φ and therefore, gain or lose consciousness. Thus, without a mechanism that can guide the network toward higher-φ states in real-time and from within, identifying such configurations would still require probing an astronomically large space of possibilities. To our knowledge, no biological mechanism has been identified that could guide networks toward higher-φ configurations through intrinsic real-time dynamics.

IIT correctly identifies information integration as a necessary property of consciousness and provides rigorous measures of network connectivity through φ, a foundational contribution RIFT explicitly builds upon. RIFT extends this foundation by addressing a question IIT was not designed to answer: how does integrated information become physically instantiated as causally efficacious experience in real time, within a single conscious moment, through a localized substrate enabling autopoietic feedback? Notably, Albantakis et al. show that perfect task performance in animat simulations was achieved with φ_max_ = 0, meaning that as defined by IIT, these organisms were *not conscious* despite solving the task (Albantakis, Hintze et al. 2014). This demonstrates that high φ reflects conditions favoring integrated network architectures under evolutionary selection pressure, a complementary finding to the focus of RIFT on the real-time mechanism of conscious moment formation.

To address the growth and evolution of the RIFT network toward optimal information integration, I introduced φ_dyn_, a computationally feasible and dynamic proxy for φ in IIT based on the “sliding window method” (see Methods for details). Standard IIT quantifies integrated information through exhaustive analysis of all possible causal partitions, determining how much the causal power of a system exceeds that of its parts (Tononi, Edelman et al. 1994, Edelman and Tononi 2000, Tononi 2004, Tononi 2008, Albantakis, Barbosa et al. 2023). This measure is computationally intractable to calculate for any given configuration in networks beyond ∼10 nodes and is typically evaluated only at single time-step transitions, limiting its applicability to evolving networks.

Specifically, φ_dyn_ provides a tractable alternative for quantifying information integration by analyzing temporal patterns of neuronal firing across different branches of the dendritic tree in N₀, capturing how information integration evolves over time rather than measuring static network properties. Unlike exhaustive partition analysis in IIT requiring 2^k^ evaluations for k partitions, φ_dyn_ tracks only the N branch-level firing windows, making computation scale linearly with dendritic complexity rather than exponentially with network size. For each branch level, we identify when its connected neurons N_i_ fire, creating time windows that represent branch-specific activity. We then quantify how these windows relate to each other temporally: overlapping windows enable information integration between branches, while separated windows indicate independent processing. The measure is defined as:

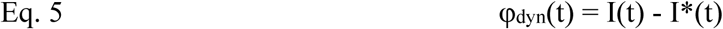

Where:

I(t) = integrated information with all branch connections intact at time t

I*(t) = integrated information when branches are considered separately at time t

φ_dyn_(t) = the time-dependent information that exists only in the inter-branch relationships

While φ_dyn_ adopts the conceptual framework of “integration exceeding partition,” it does not implement formal φ calculation based on causal partitions as in IIT. It is presented as a biologically tractable proxy that captures temporal dynamics of integration not addressed by static causal analysis. This formulation follows the sliding window method of calculating φ* used in recent thalamocortical studies (Munn, Muller et al. 2023, Whyte, Muller et al. 2025), but adapts it to quantify integration across dendritic branch timing patterns rather than probabilistic independence between neurons.

Using this measure, φ_dyn_ peaks only when the firing windows of different dendritic branches are separated enough to remain distinct yet not so far apart that the branches become functionally independent. Figure 5A (Code 2) shows the external distances of neurons connected to each branch, which determine the timing offsets that give rise to the distinct branch-specific firing windows in Fig. 5B. The spatial organization of neurons at different distances translates into temporal separation of their firing windows due to signal propagation delays. When windows overlap within the temporal grain (2.0 msec), as in adjacent branches, they merge and eliminate measurable integration, yielding φ_dyn_ ≈ 0. When windows are extremely far apart, as in B1–B3 or B1–B4, the branches no longer interact and φ_dyn_ drops as well. The strongest integration occurs at intermediate gaps, illustrated by B2–B4 in Fig. 5C, where the windows are clearly separated but still dynamically coupled.

**Figure 5.**
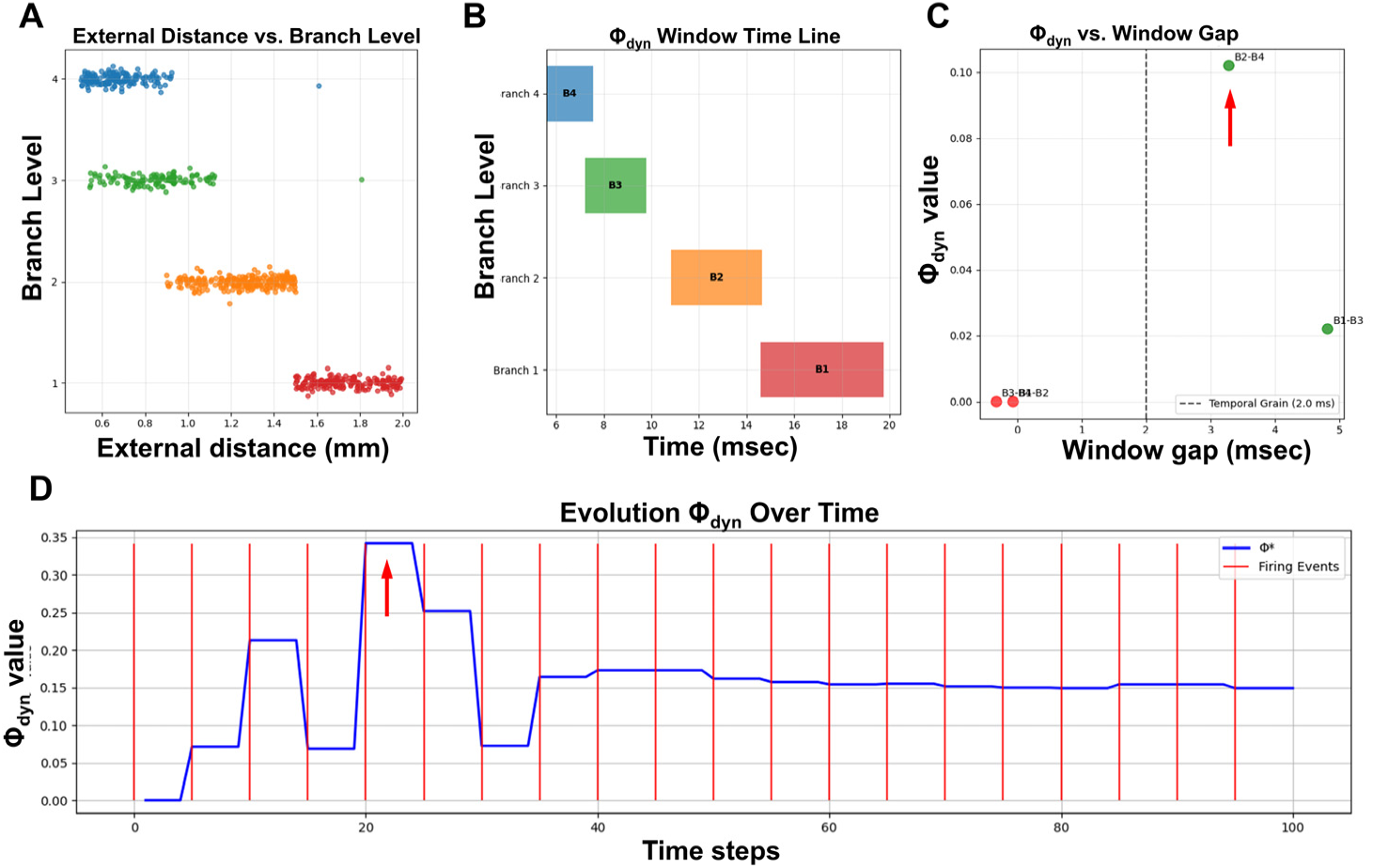
Dynamic integrated information (φ_dyn_) peaks at optimal temporal window separation. (A) External distance vs. branch level: four fractal-scaled clusters (r = 0.6). (B) Temporal firing windows for branches B1–B4 (6–20 ms range), illustrating hierarchical window structure. (C) φ_dyn_ vs. window gap: peak integration (φ_dyn_ = 0.10) at ∼4.0 ms separation (B2–B4); integration drops at both smaller and larger gaps. (D) Evolution of φ_dyn_ over 100 timesteps: stepwise rise with prominent jump at timestep ∼20 (B2–B4 configuration established), stabilizing at ∼0.15 after step 50. Temporal grain = 2.0 ms.

Figure 5D shows the evolution of φ_dyn_ over time as neurons at different external distances begin firing (red vertical lines), establishing recurrent connections that create the temporal windows shown in Fig. 5B. Each step-like increase in φ_dyn_ corresponds to new neurons joining the network, with the most prominent jump occurring around timestep 20 when the optimal branch-level configuration (B2-B4 spacing) becomes established. φ_dyn_ then stabilizes at approximately 0.15 around timestep 50 once all branches maintain their characteristic temporal offsets and the recurrent network structure is complete.

Having established how branch-level timing patterns generate φ_dyn_, we next examined how the somatic delay, which sets the time window for coincidence between dendritic inputs and somatic integration, modulates these patterns across the full network. The somatic delay is a critical timing parameter that shapes network oscillation frequency and the strength of temporal integration by determining the temporal dynamics linking signals from the fractal dendritic tree (N₀) and peripheral network activity (N_i_). To characterize this relationship, we extended the RIFT simulation (Code 1) to a systematic delay-sweep analysis (Code 3, *run_improved_delay_sweep*) covering delays from 1–20 ms with multiple repetitions per delay value. For each delay setting, φ_dyn_ time series were calculated using the sliding window approach (Code 1, *PhiStarCalculator.calculate_phi_star* method), which bins neuronal firing times into temporal windows and calculates information integration as φ = I – I*, where I represents integrated information across time windows and I* represents reducible information from independent windows. The resulting φ_dyn_ time series were then analyzed for temporal volatility (std(dφ_dyn_/d(step))), predictability, and phase-space dynamics (Codes 5-7). This comprehensive parameter exploration revealed the gradients for φ_dyn_, fractal measures, and predictability shown in Fig. 6.

**Figure 6.**
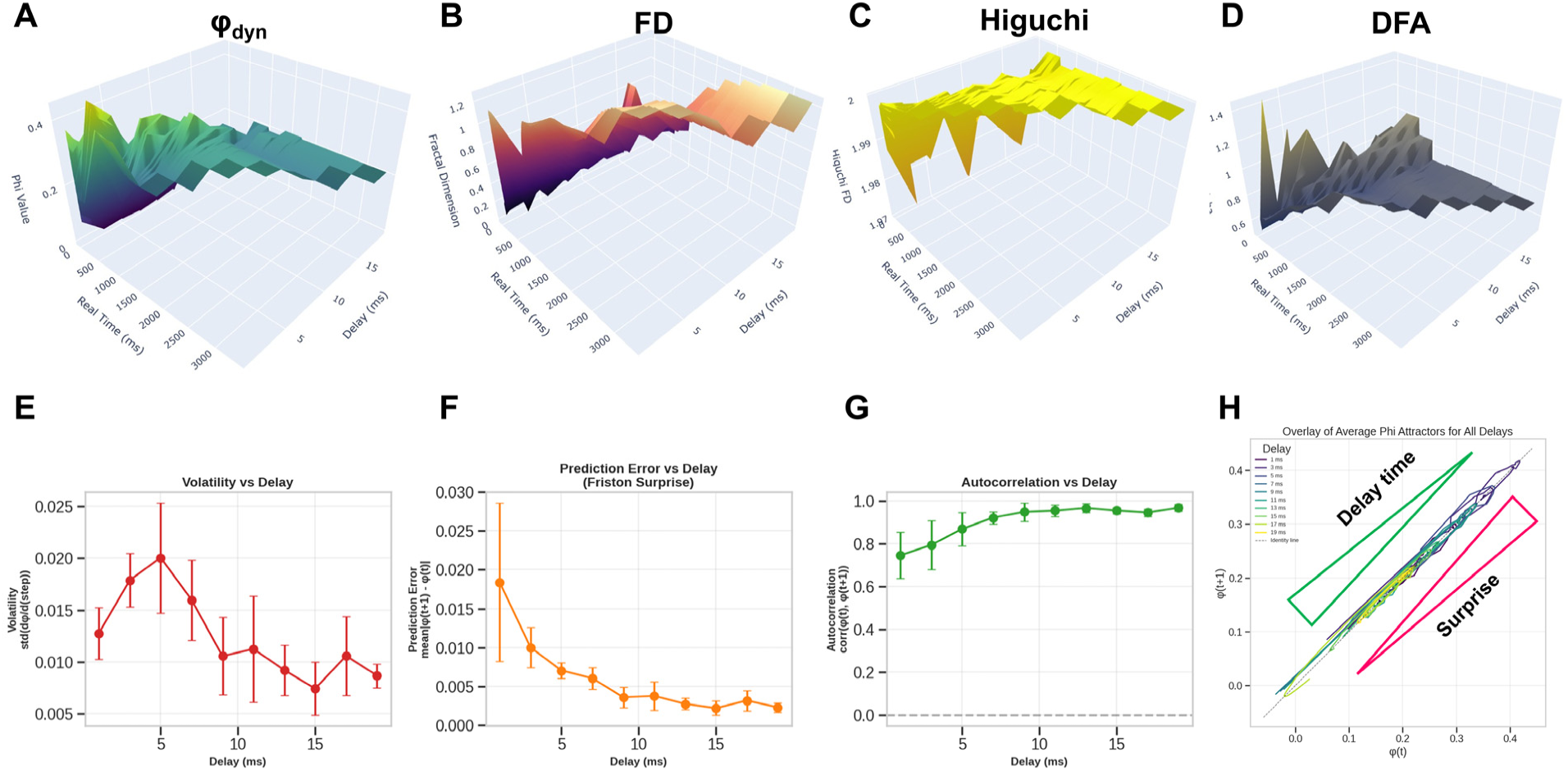
Somatic delay modulates fractal architecture and creates continuous predictability gradient. (A–D) 3D surface visualizations across simulation time (0–3,000 ms) and somatic delay (1–20 ms): (A) φ_dyn (0–0.4), (B) Fractal Dimension (0.2–1.2), (C) Higuchi FD (1.87–2.0), (D) DFA α exponent (0.5–1.4); all converge to delay-dependent plateaus. (E) Volatility vs. delay: 55% reduction from 5 to 19 ms (n = 5 per delay). (F) Prediction error vs. delay: 9-fold reduction from 1 to 19 ms, mapping exploration-to-exploitation continuum. (G) Autocorrelation vs. delay: r increases from 0.75 to 0.97. (H) Phase-space attractors (φ_dyn_(t) vs. φ_dyn_(t+1)) for all delays overlaid near identity line: green arrow indicates delay-dependent attractor spread reflecting exploration-exploitation continuum; red arrow indicates surprise zone reflecting high prediction error at short delays. n = 5 simulations per condition.

3D surface visualizations (Fig. 6A–D) show that φ_dyn_ and fractal measures behave differently across delay times. The φ_dyn_ surface (Fig. 6A) has more variation at short delays, reflecting the larger transient fluctuations early in the simulation, but it converges to stable values at later time points for all delays. In contrast, fractal measures (FD in 6B, Higuchi FD in 6C, and DFA in 6D showing similar patterns) rise from their initial transients and then stabilize into delay-dependent plateau values, producing smoother surfaces overall. The fractal measures capture the stable spatial organization of the network, whereas φ_dyn_ reflects the temporal alignment of branch-specific activity, which depends strongly on delay. Despite these differences, both φ_dyn_ and the fractal metrics ultimately stabilize, with delay mainly affecting the shape and duration of the early transient.

Quantitative analysis of this temporal evolution reveals a continuous predictability gradient across delays. Three complementary metrics quantify this continuum: (1) volatility (std(dφ_dyn_/d(step)) = 0.020 at 5 ms declining to 0.009 at 19 ms) (Fig. 6E), (2) prediction error (mean|φ_dyn_(t+1) - φ_dyn_(t)| = 0.018 at 1ms declining to 0.002 at 19 ms) (Fig. 6F), and (3) autocorrelation (r = 0.75 at 1ms increasing to 0.97 at 19 ms) (Fig. 6G). The prediction error metric directly quantifies “surprise” in the Free Energy Principle framework (Feldman and Friston 2010, Friston 2010, Friston, Daunizeau et al. 2010), with the 9-fold decline across delays demonstrating a shift from surprise-driven consciousness states (short delays, high prediction error, exploratory dynamics) to expectation-driven consciousness states (long delays, low prediction error, confirmatory dynamics). This continuum maps naturally onto exploration-exploitation trade-offs in active inference, where different delays represent different strategies for balancing surprise tolerance against stability.

Phase-space trajectories (Fig. 6H) reveal the dynamical structure underlying this predictability continuum. All delays exhibit stable attractor dynamics with trajectories closely following the identity line (φ_dyn_(t+1) ≈ φ_dyn_(t)), indicating convergence toward equilibrium states rather than chaotic or limit-cycle behavior (see Methods for details on fractal attractors). Notably, short delays show wider excursions from the identity line (higher volatility) while long delays remain tightly clustered (high predictability), consistent with the volatility measurements. The overlay of all delays (Fig. 6H) demonstrates that the attractor structure itself is preserved across the full delay spectrum, only the tightness of convergence varies, supporting the interpretation of a continuous gradient rather than discrete regime transitions.

This pattern of moderate φ_dyn_ values coupled with high temporal stability (dφ_dyn_/dt(step) ≈ 0) at longer delays aligns with a continuous “upload and update mechanism” for consciousness, wherein a limited amount of new information from incoming EPSP signals is steadily integrated with a pre-existing base of conscious content at the soma. The longer delays provide the necessary temporal window for this integration process, essentially defining a conscious moment and suggesting that consciousness may be fundamentally incremental rather than regenerative in its information integration mechanism. Building on the original concept of φ (IIT), φ_dyn_ describes a dynamic network converging toward stability in which incremental but continuous uploading of new information from EPSPs avoids φ fluctuation and breakdown of continuity in consciousness due to rapid changes in neuron connectivity, providing both the stability needed for continuous subjective experience and the flexibility to incorporate new information as it arrives.

### Part 3: Irreducible Multifractals - Folding Infinity into Finite Space through Generational Fractal Mapping (GFM)

A continuous upload mechanism requires a physical substrate within the core neurons of the RIFT network. Having established at the network level that somatic delay enables incremental integration of new information to update consecutive moments in consciousness (Parts 1-2), we now examine the molecular mechanisms within the soma that could implement this integration. This substrate must be both programmable by fractal input from the dendritic tree and sustain fractal mapping of the Self onto each part of the substrate. Through the whole-in-part property characteristic of fractal topology, each spatial region contains compressed representations of the entire pattern, generating the self-referential architecture essential for consciousness. Hence, we need to define a molecular mechanism by which information extracted from EPSPs is continuously mapped onto a substrate that sustains fractal self-referential organization.

Previously, I proposed that sentyons, fractal “conscious particles” generated by ion channel-lipid domains in the somatic plasma membrane of core neurons, provide this substrate by integrating synaptic input with a pre-existing base consciousness (Bieberich 2012). While sentyons were initially described as irreducible representations of conscious moments, they had not been formally defined as multifractals, nor was it specified how they transfer information across successive inputs to preserve and update base consciousness.

Any theory assuming a molecular substrate of consciousness must transform neural activity, either action potentials in a network or synapse activation in a dendritic tree, into a spatial array within neuronal cell membranes, the cytoskeleton, or other molecular substrates. Standard approaches to dendritic-somatic integration typically collapse complex spatiotemporal dendritic activity into scalar measures at the soma, such as integrated membrane potential or firing rate. However, dendritic inputs from different distances produce distinct signatures at the soma through distance-dependent attenuation and temporal summation patterns (Williams and Stuart 2002), suggesting that branch-level specific spatial location information is preserved in the temporal dynamics of somatic integration. If the neural activity is fractal, the mapping of this activity onto the molecular substrate must preserve the fractality. Because any molecular substrate has physical limits to downscaling, true infinite fractal operations are impossible.

Hence, preserving fractality requires solving how to fold infinity into finite space. A potential solution replaces infinite spatial downscaling with sequential temporal operations, in which the fractal computational architecture of embedding self-referential states into a molecular substrate emerges as a necessary outcome of the algorithm. Recent theories have recognized Layer 5 pyramidal neurons and somatic integration as critical for consciousness, including Dendritic Integration Theory (DIT) emphasizing dendritic-somatic coupling (Aru, Suzuki et al. 2020, Bachmann, Aru et al. 2020, Suzuki and Larkum 2020, Storm, Boly et al. 2024) and Orchestrated Objective Reduction (Orch OR) proposing microtubule-based processes in dendritic-somatic regions. RIFT differs by proposing multifractal spatial encoding in the somatic membrane as the substrate.

To implement fractal encoding in physical substrates such as the somatic membrane, we must first map the information contained in EPSPs onto a spatial lattice capable of preserving fractal organization. Code 8 implements a simplified theoretical model for this mapping of EPSPs onto a sentyon in a RIFT network. In this model (see Part 1), the dendritic architecture follows a fractal branching rule with scaling factor r = 0.6: branch lengths decrease as L₀ꞏr^(i–1)^ across successive levels, and synapse distribution scales accordingly to produce integrated EPSP train responses following A₀ꞏr^(i–1)^, where i denotes branch level (1-4), parameters consistent with moderate distance-dependent attenuation in neocortical Layer 5 pyramidal neurons as discussed in Part 1 (Williams and Stuart 2002). These neurons participate in recurrent thalamocortical and corticocortical loops proposed to be critical for consciousness (Dehaene and Changeux 2011). While active dendritic mechanisms normalize single EPSP amplitudes, trains of synaptic inputs from different branch levels produce distinct temporal summation patterns at the soma (Magee and Cook 2000, Williams and Stuart 2002). The implementation of Code 8 in Fig. 7 shows how these EPSP trains are derived from branch-specific signals at each dendritic level (Fig. 7A and B) and projected onto a 243×243 grid representing an idealized patch of somatic membrane after temporal coincidence detection at the soma. Peak amplitudes from the EPSP signals are hierarchically assigned to nested spatial scales following a Sierpinski carpet pattern (Fig. 7C). These hierarchical assignments are then additively integrated to generate the final multifractal intensity distribution (Fig. 7D). This fractal mapping provides a theoretical framework for how temporal summation signatures from EPSP trains could be encoded as spatial patterns in the somatic membrane.

**Figure 7.**
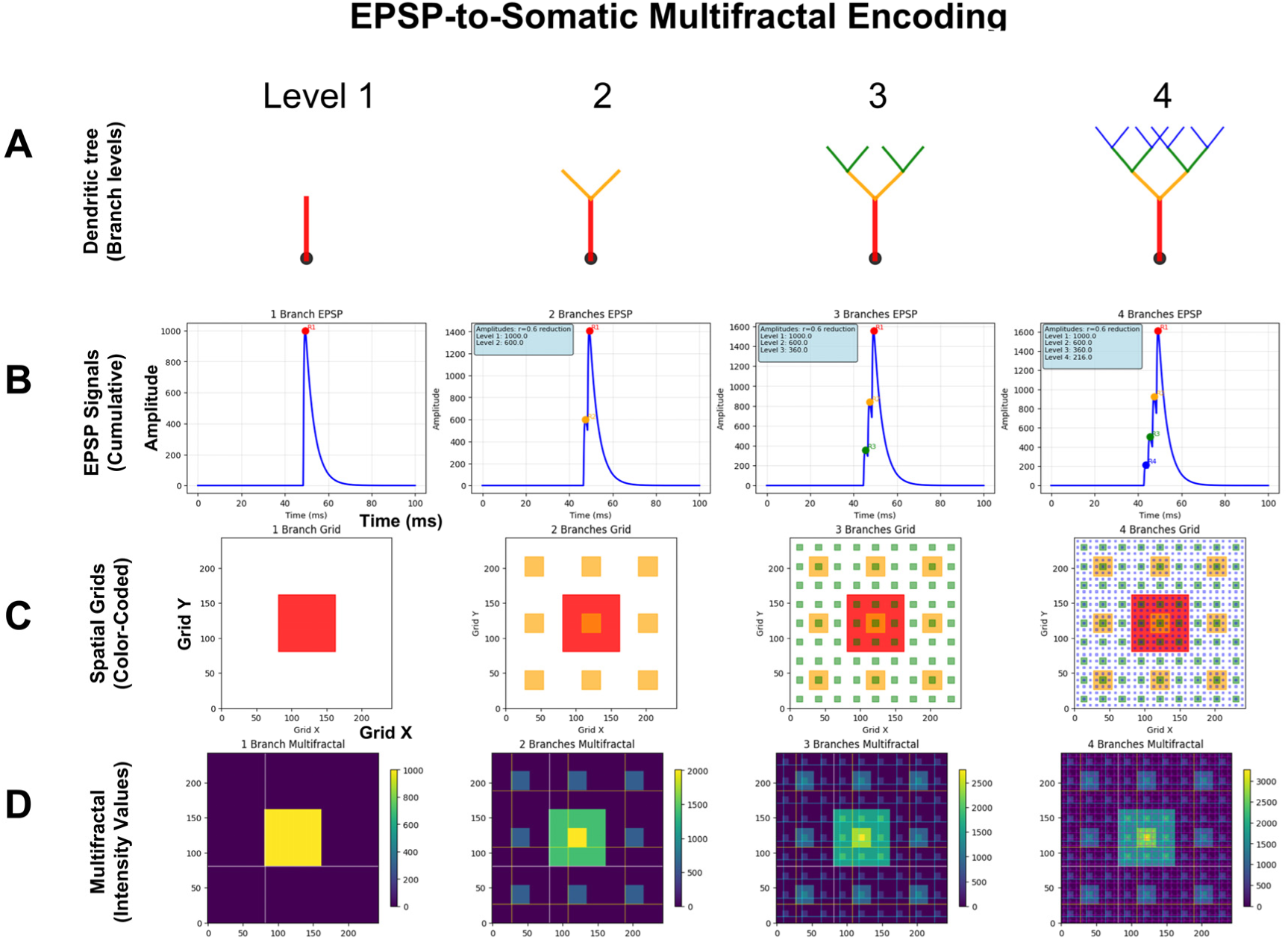
EPSP-to-somatic multifractal encoding preserves hierarchical dendritic structure. (A) Fractal dendritic architecture (r = 0.6), providing hierarchical synaptic input for panels B–D. (B) Cumulative EPSP waveforms for 1–4 active branch levels (amplitude scaling A₀ꞏr^(i−1)^, peak ∼50 ms). (C) Spatial EPSP mapping onto 243×243 somatic grid (Gaussian kernel): progressive complexity from single hotspot (Level 1) to fully hierarchical multifractal (Level 4). (D) Additive multifractal integration across branch levels: Level 4 exhibits self-similar structure with central concentration (∼2,500–3,000 units) and fractal periphery. Lattice: 243×243, amplitude scaling A₁=0.15 to A₄=0.032.

When mapped onto the multifractal lattice, the hierarchically scaled EPSP trains generate spatial intensity distributions that preserve the fractal structure inherent in dendritic integration. The mapping process assigns EPSP train responses additively at successive grid scales, thereby translating the power-law amplitude distributions (amplitude hierarchies) that emerge from summed dendritic inputs into corresponding spatial hierarchies (Fig. 7). This fractal lattice mapping maintains scale-invariant relationships, enabling the somatic membrane to encode which branches at which hierarchical levels were activated through their distinct temporal summation signatures. This branch-level specific spatial encoding enables quantitative measurement of whole-in-part properties and information integration across spatial scales, relationships that are lost when complex spatiotemporal patterns are collapsed into single numerical values.

Building on this EPSP-to-multifractal integration mechanism (Code 8), our dynamic theoretical model (Code 9) implements generational fractal mapping (GFM) shown in Fig. 8A. EPSP peak amplitudes detected from each branch level are hierarchically mapped onto nested spatial scales following a Sierpinski carpet pattern (Code 9, *generate_fractal* method), creating the initial multifractal. This pattern then undergoes autonomous development through morphological expansion and energy conservation dynamics (*grow_fractal_structure* method). As EPSPs generated from proximal to distal branches of the dendritic tree converge at the soma (Fig. 7), each EPSP peak contributes unique spatial complexity, creating branch-level irreducibility that recapitulates the network-level irreducibility established in Parts 1 and 2. Growth dynamics continue until the center region reaches critical threshold intensity, triggering a multifractal collapse that initiates an action potential. During collapse, the most information-rich peripheral quadrant is extracted as the holographically compressed seed for the next generation (Fig. 8A), showing high structural similarity and information density ratio through whole-in-part analysis. New EPSP inputs are then uploaded and integrated with this seed pattern to generate a daughter fractal, which grows into the next generation parent until reaching threshold intensity again, establishing cyclical dynamics (GFM cycle).

**Figure 8.**
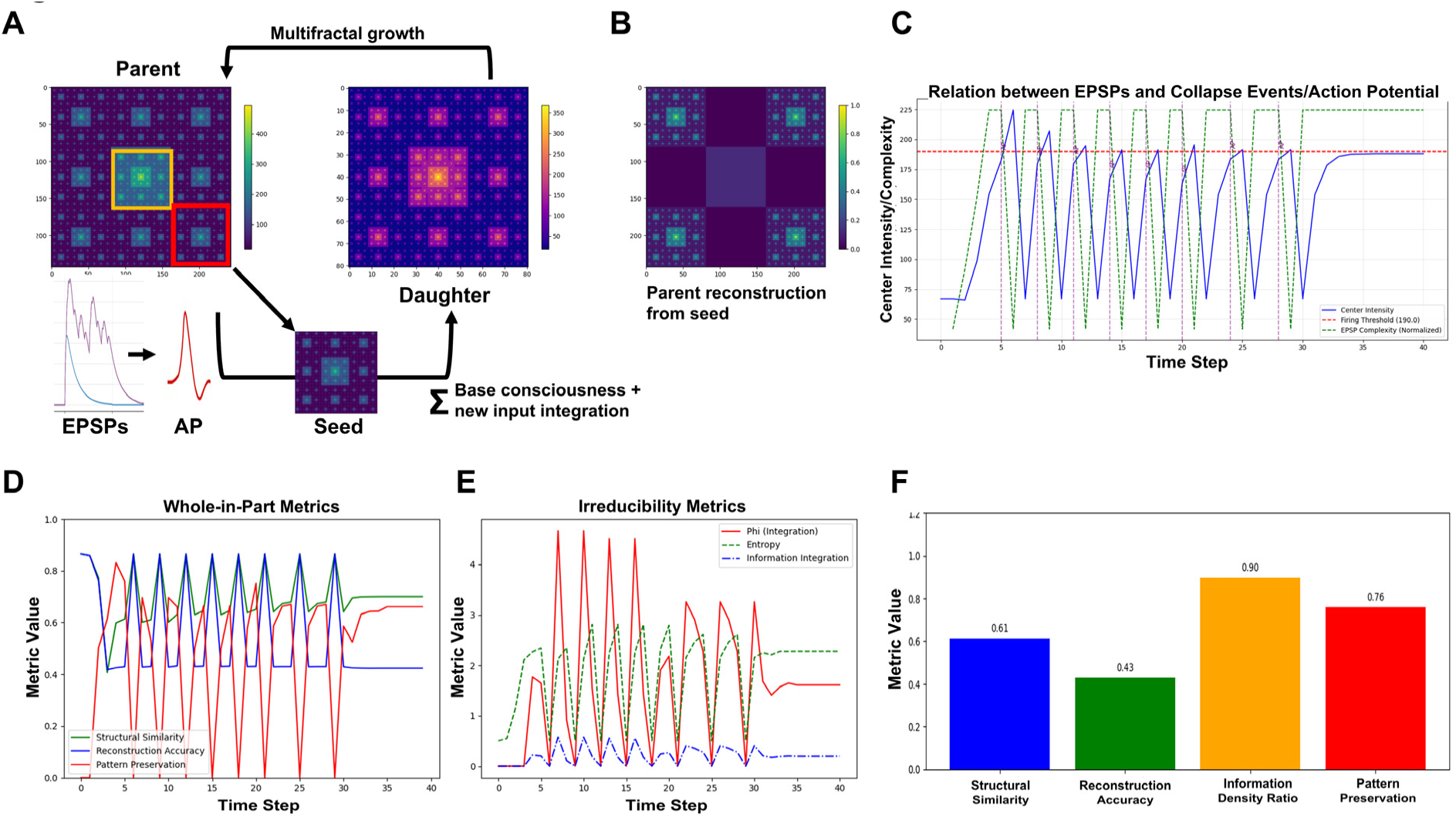
Generational fractal mapping (GFM) implements whole-in-part encoding. (A) GFM cycle schematic: EPSP input → multifractal growth → threshold-triggered collapse → action potential → peripheral seed extraction → daughter fractal generation. (B) Parent reconstruction from seed (structural similarity 0.61, information density ratio 0.90). (C) Center intensity and action potentials over 45 timesteps: sawtooth pattern with regular collapse events (∼every 3–4 steps). (D) Whole-in-part metrics (structural similarity, reconstruction accuracy, pattern preservation) oscillating 0.4–0.9 across GFM cycles. (E) Irreducibility metrics: φ peaks (>4.5) at collapse events, with complementary entropy and integration dynamics. (F) Averaged cycle metrics: structural similarity 0.61, reconstruction accuracy 0.43, information density ratio 0.90, pattern preservation 0.76. n = 40 timesteps; firing threshold = 180.0.

Figure 8B-E demonstrate key properties of this system. Figure 8B confirms holographic information preservation: the extracted peripheral seed successfully reconstructs the parent’s essential spatial structure when expanded. Figure 8C shows temporal coupling between EPSP integration (green, normalized complexity), center intensity accumulation (blue), and action potential generation (purple vertical lines) when the firing threshold (red dashed) is exceeded, with regular collapse events approximately every 3-4 steps (depending on the actual threshold value, here 180). Figure 8D reveals that whole-in-part metrics, structural similarity (green), reconstruction accuracy (blue), and pattern preservation (red), oscillate systematically with each collapse. Figure 8E quantifies irreducibility: φ*(multifractal) (red) peaks during collapses, while entropy (green dashed) and integration (blue dash-dot) show complementary dynamics, together demonstrating sustained irreducible complexity throughout generational cycling by GFM: averaged values across all cycles (Fig. 8F) confirm robust and consistent performance of these metrics.

GFM is informationally equivalent to infinite downscaling (IDS). In traditional fractal theory, the spatial scaling factor r describes how much smaller each embedded pattern becomes. However, if we reinterpret r as the information preservation rate during downscaling - e.g., if r = 0.6, then 60% of information is retained at each smaller scale.

This gives total information:

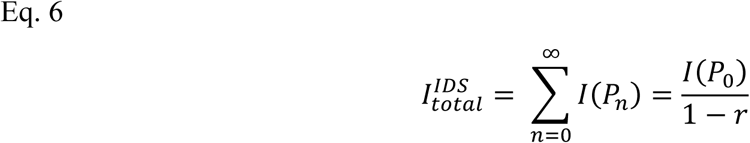

Where:

P₀: The original parent fractal pattern at the largest scale (scale level 0)

Pₙ: The embedded fractal pattern at scale level n, where each Pₙ is a smaller, self-similar copy of P₀

Similarly, GFM uses a temporal decay factor q representing how much information transfers from one generation to the next, e.g., if q = 0.6, each new generation preserves 60% of the previous generation’s information, yielding cumulative information

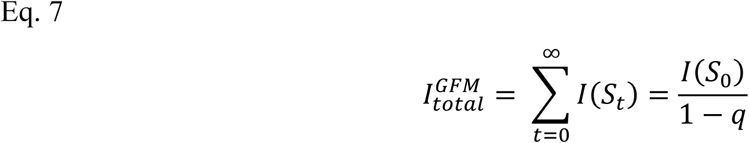

Where:

S₀: The initial seed pattern extracted from the first parent fractal (generation 0)

Sₜ: The seed pattern at generation t, which carries forward information from previous generations through the temporal succession process.

When both systems preserve information at the same rate (q = r), then

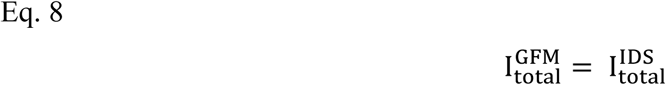

This equivalence demonstrates that temporal succession can achieve the same information-theoretic capacity as infinite spatial embedding. While physical systems cannot implement IDS due to quantum limits, temporal succession faces no such constraint, biological systems can continue generating new fractal generations indefinitely. GFM thus provides a physically realizable alternative that preserves the information-theoretic properties of classical fractal systems.

Integration of information in the parent fractal and inheritance to the daughter is quantified using Kullback-Leibler (KL) divergence, which measures the difference between two probability distributions. KL divergence forms the mathematical foundation of early IIT formulations, where it calculated integrated information φ by comparing whole-system distributions against independently-partitioned components (Tononi 2004). In our implementation, temporal information encoded in EPSP trains is mapped onto a spatial multifractal, transforming temporal into spatial fractals. After each EPSP-driven update, we partition the structure into four quadrants and calculate KL divergence between the probability distribution of the whole fractal (treating all intensity values as a unified distribution) and that of its partitioned components (treating the intensity values of each quadrant as independent distributions). This calculation, implemented in Code 9 (*compute_irreducibility_fixed*) function, directly yields φ*(multifractal):

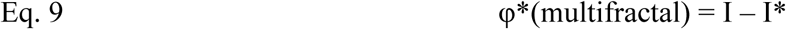

Where:

I = integrated information of the whole multifractal pattern

I* = integrated information when quadrants are treated as independent parts

φ*(multifractal) = the irreducible information that exists only in the spatial relationships between quadrants

As new EPSP inputs integrate with the seed pattern to generate updated multifractal structures, both whole-pattern and quadrant distributions evolve, causing φ*(multifractal) to vary across the generational cycle. This approach mirrors φ_dyn_ from Results Parts 1 and 2, both adopt the integration-versus-partition framework of IIT but apply it to different domains. While φ_dyn_ measures temporal integration across evolving network connectivity in the dendritic tree, φ*(multifractal) measures spatial integration within the somatic substrate at single time points. Although the causal partition analysis in IIT provides a rigorous structural foundation, our method additionally captures how φ*(multifractal) changes after each update cycle, reflecting consciousness as an ongoing process rather than a fixed state. This dual application of KL divergence, quantifying information gained through updating (successive generations) and information lost through spatial partitioning (whole versus parts), unifies temporal and spatial aspects of consciousness within a single mathematical framework. Together, φ_dyn_ and φ*(multifractal) reveal how consciousness emerges from both temporal coordination in neural networks and spatial organization in the somatic substrate, with GFM linking multifractal geometry to neural timing patterns.

Beyond compression, GFM enables a representational architecture where information supporting contradictory propositions can coexist through fractal embedding. In classical feedback systems, new states overwrite old ones. In GFM, the seed extraction mechanism preserves compressed traces of prior states within the fractal structure, allowing contradictory information to occupy distinct but mutually embedded subspaces through the whole-in-part property. Crucially, GFM implements differential encoding: the seed carries compressed historical context (q fraction of previous information), while new EPSPs carry only delta information: what has changed or violated predictions. This weighted integration M(t) = qꞏSeed(t-1) + (1-q)ꞏΔEPSP(t) structurally resembles Bayesian updating where priors combine with new evidence, though RIFT implements this through geometric pattern combination rather than probability calculations. The compression advantage is substantial: encoding complete cortical states requires mapping 10⁶-10⁷ active synapses onto ∼10⁴ molecular sites (100-1000:1 compression); encoding deltas requires mapping only 10⁴-10⁵ changed events onto ∼10³ available sites (10-100:1 compression per cycle), achieved through fractal encoding of hierarchical EPSP amplitude distributions.

Consider how GFM preserves temporal information through fractal embedding: At generation t₁, EPSPs encode “object is red” as multifractal pattern M₁ with distributed representations across quadrants. During collapse, a seed is extracted preserving this compressed pattern. At t₂, new EPSPs encoding “object is green” combine with the seed to generate daughter pattern M₂. This composite multifractal simultaneously contains both “red” (compressed in seed) and “green” (current EPSPs), representing “the object was X and is now not-X.” This mirrors conscious experience: we recognize the traffic light turned green while remaining aware it was previously red. The whole-in-part structure enables conscious awareness of temporal continuity, the current moment contains compressed traces of prior moments through fractal embedding. When GFM is disrupted, this inability to consciously relate present to past breaks down episodic memory, leaving individuals disoriented as observed in frontotemporal dementia and Alzheimer’s disease.

GFM implements the “fractal cNOT gate” operation previously described for resolving logical paradoxes such as the “Liar’s paradox”, although without applying quantum computation (Bieberich 2001). This capacity for maintaining contradictory temporal traces through fractal embedding has two critical consequences. First, it enables temporal continuity: consciousness maintains awareness of change because contradictory states coexist within the fractal structure rather than replacing each other sequentially. Second, it supports continuous deliberation: consciousness maintains access to alternative interpretations, enabling ongoing integration of new information with contradictory historical traces rather than forcing premature collapse to binary decisions like classical algorithms.

The mechanisms developed across Parts 1-3 collectively address several interconnected challenges outlined in the Introduction. The neural degeneracy problem, wherein different input patterns produce identical outputs, is resolved through multilevel fractality: fractal dendritic architecture preserves positional information through amplitude and timing hierarchies (Part 1), temporal φ_dyn_ dynamics distinguish patterns with identical static integration (Part 2), and multifractal spatial organization retains distribution patterns with irreducible φ*(multifractal) that scalar summation would eliminate (Part 3). Together, these ensure that different conscious states correspond to distinguishably different network-dendritic-multifractal configurations. The extreme compression paradox, maintaining experiential richness despite million-fold information reduction, is addressed through differential encoding in GFM, where only delta information requires encoding per cycle (10-100:1 compression) rather than complete states (100-1000:1 compression). Fractal embedding enables this compressed representation to preserve temporal context through the whole-in-part property, explaining how conscious experience maintains continuity and richness despite slow update rates.

However, compression alone does not explain phenomenological experience, nor does it specify the molecular substrate that physically implements multifractal encoding. Part 4 now identifies this substrate: an Ising lattice of ion channels and lipid domains where reciprocal fractal dynamics enable multifractal self-organization without preconfigured geometry, and where temporal EPSP sequences are transformed into spatially distributed somatic fields through dendritic branch-level mapping. Part 5 then demonstrates how this lipid-ion channel substrate enables holographic projection and autopoietic control.

### Part 4: Somatic Information Integration - Membrane Lipids as the Primary Substrate for Neural Information Processing

The multifractal model and GFM framework demonstrate how temporal EPSP sequences generate spatial multifractal patterns that encode information across consecutive conscious moments (Part 3). However, extending this computational framework to biological implementation requires addressing two critical gaps: (1) identifying the specific molecular substrate that physically implements multifractal pattern formation in neurons, and (2) explaining how the endospace, the spatiotemporal dimension of inner experience, unfolds from this substrate and achieves autopoietic modulation of neural dynamics. While Part 3 mentioned prior theoretical work proposing that sentyons, fractal lattices of ion channels and lipid domains, serve as consciousness substrates (Bieberich 1998, Bieberich 2002, Bieberich 2012), it did not specify the molecular mechanisms. Part 4 now formalizes how EPSPs program this lipid-channel substrate.

The biological multifractal implementation uses an Ising lattice where ion channels and surrounding lipid domains interact dynamically, with both components exhibiting intrinsic fractal properties (see Methods). EPSPs display power-law temporal dynamics and scale-invariant behavior, while membrane lipids self-organize into fractal spatial clusters at biologically relevant scales (Starr, Hartmann et al. 2014). EPSPs activate voltage-gated ion channels through membrane depolarization, while lipid rafts modulate channel conductance and gating kinetics. Critically, membrane depolarization induces lipid reorganization, creating reciprocal dynamics where channel activity reorganizes lipid domains, which in turn modulate subsequent channel responses (Suma, Sigg et al. 2023, Suma, Sigg et al. 2024). This convergence of fractal temporal and spatial dynamics enables multifractal pattern self-organization without preconfigured geometry.

Temporal EPSP amplitudes from different dendritic branch levels are converted to spatially distributed membrane depolarization fields via Gaussian kernel mapping (see Methods, Code 10). While experimental data on branch-level EPSP amplitude attenuation exists (Williams and Stuart 2002), the specific spatial distribution of these signals across the somatic membrane remains experimentally undetermined. The Gaussian kernel approach provides a biologically plausible computational framework for modeling this temporal-to-spatial transformation, with branch-level-specific parameters (spatial center and radial extent) preserving the hierarchical dendritic structure in spatial form. The resulting electric field

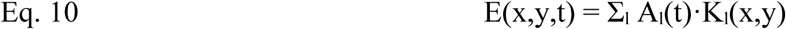

where Aₗ(t) is the EPSP amplitude from dendritic branch level l at time t, and Kₗ(x,y) is a Gaussian kernel that maps this temporal signal to a spatial region on the somatic membrane (see Methods for kernel specification). Proximal branches map near the lattice center, distal branches at increasing radial distances, preserving dendritic hierarchy in spatial form. The electrical field drives ion channel opening through lipid-mediated coupling:

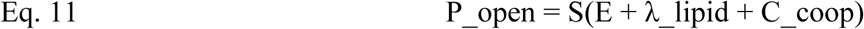

where S is a sigmoid activation function, λ_lipid represents local lipid bias effects, and C_coop captures cooperative effects from nearby open channels. Once EPSPs program initial channel states, Ising interactions govern lipid domain reorganization, creating spatially structured domains: lipid state +1 associated with open channels, −1 with closed channels that directly modulate subsequent channel opening probabilities through the λ_lipid term.

Figure 9A illustrates the GFM cycle in the biological multifractal, analogous to the theoretical model in Fig. 8A. The left panel shows the pre-collapse state where EPSPs from the dendritic tree program the lipid-channel lattice in the soma. Lipid domains (red and blue regions) contain embedded ion channels (green=open, black=closed), with concentrated activity at the center. The right panel shows the post-collapse state where an action potential is generated, lipid domains reorganize, and a seed region is preserved for the next generation. Unlike the theoretical multifractal, where the seed is an abstract pattern excerpt, in the biological implementation the seed is physically instantiated in the peripheral lipid domain configuration, a molecular memory that survives the action potential because lipid states, unlike channel states, are not reset at collapse.

**Figure 9.**
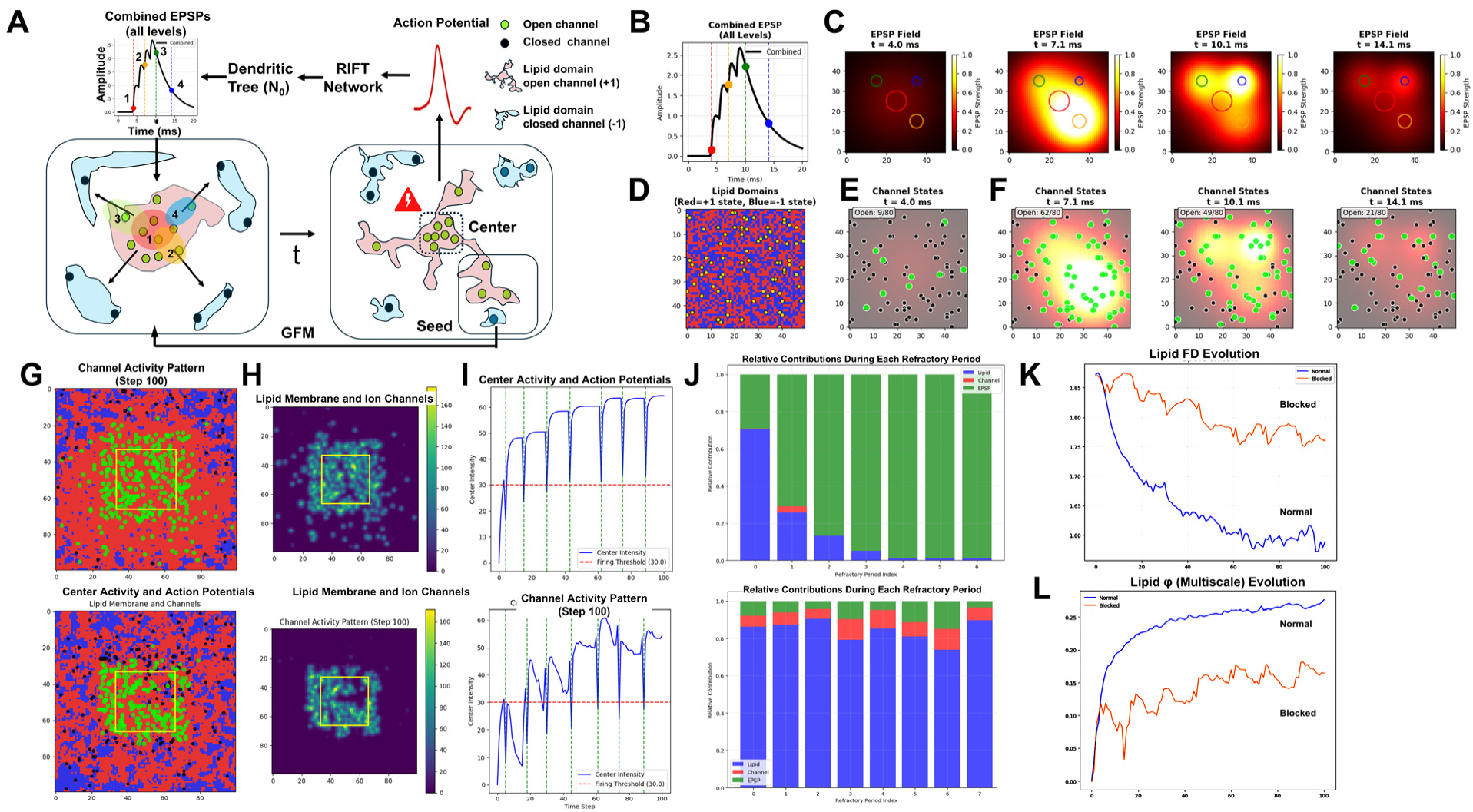
Biological multifractal implements lipid-mediated information integration with EPSP programming. (A) GFM cycle in core neuron: hierarchical EPSP inputs program lipid domain organization, driving action potential generation and seed extraction for recurrent loop formation. (B) Combined EPSP temporal profile (peak amplitude ∼2.3 at 9 ms). (C) EPSP field spatial distribution (Gaussian kernel) at four timepoints (t = 4.0–14.1 ms), showing field decay and spatial reorganization. (D) Initial lipid state (t = 0): uniform binary distribution (∼50% red/blue). (E–F) Ion channel states at four timepoints: progressive channel activation following EPSP dynamics. (G-H) Lipid and channel activity heatmaps at step 100: continuous mode shows distributed activity; blocked mode shows central concentration with phase-separated peripheral domains. (I) Center intensity and action potentials for continuous vs. blocked conditions: blocked mode shows progressive baseline rise (∼30 to ∼35 over 100 steps), demonstrating lipid memory accumulation. (J) Relative contributions per refractory period: EPSPs dominate in continuous mode (∼85%); lipids dominate in blocked mode (∼85–95%, p < 10⁻⁴⁰). (K) Lipid FD evolution: blocked condition maintains elevated FD >1.75 vs. normal decline (p = 2.74×10⁻⁴¹). (L) Lipid φ evolution: normal ∼0.25–0.30, blocked ∼0.10–0.15. Lattice: 100×100 (0.25 µm/pixel); ∼400 ion channels; n = 100 steps.

Figure 9B-F demonstrates this EPSP-to-spatial-field translation process in detail (Code 10). Figure 9B displays the combined EPSP temporal profile from all dendritic branch levels, showing the hierarchical amplitude summation that drives somatic integration. Figure 9C shows the EPSP spatial field at t = 4.0 ms, with branch level 1 (proximal, highest amplitude) positioned at lattice center (red circle). Figure 9D shows the initial lipid domain distribution (red = +1 state, blue = −1 state) before EPSP activation. Figure 9E-F show corresponding channel states at four time points (t = 4.0, 7.1, 10.1, 14.1 ms; green = open, black = closed), demonstrating how the EPSP field progressively activates channels and initiates the self-organizing lipid-channel dynamics that drive center-focusing, threshold accumulation, and multifractal pattern formation in the complete model.

The biological model (Codes 11 and 12) implements two modes of EPSP integration during neural evolution, revealing distinct fractal and information integration signatures (Fig. 9G-L). Both modes operate with the same consciousness-level parameter (set to 0.5, corresponding to ∼12 ms refractory periods following each action potential). Although this parameter represents one potential target of autopoietic modulation, Part 5 identifies γ, the lipid-ion channel coupling strength, as the primary autopoietic target. In the continuous mode (block_epsp_during_refractory = False), new EPSP trains are continuously integrated throughout the refractory period as multifractal patterns grow toward the firing threshold. In the blocked mode (block_epsp_during_refractory = True), EPSPs are generated progressively through proximal to distal branches only at the onset of each refractory period, after which the multifractal evolves through autonomous lipid membrane dynamics until exceeding the center activity threshold and generating an action potential.

Figure 9G-J compares these two integration modes. Figure 9G shows the final lipid membrane and channel spatial organization at step 100 (top: continuous mode, bottom: blocked mode), with green boxes highlighting the central region used for intensity measurements. In continuous mode, lipid domains show a red-dominated center with more heterogeneous red/blue mixing in the periphery, indicating that continuous EPSP input maintains central lipid polarization but prevents full spatial organization. In blocked mode, lipid domains show more abundant boundary structures and larger, more uniform domains in the periphery (corners and edges), demonstrating clear phase separation. This peripheral domain organization demonstrates that autonomous Ising dynamics allow lipids to relax into organized configurations, particularly in regions with less active channel perturbation. Figure 9H shows the corresponding cumulative channel activity patterns at step 100 as heat maps. Continuous mode (top) reveals a more distributed, spread-out channel activity across the lattice, while blocked mode (bottom) shows a concentrated hotspot at the center with minimal peripheral activity, indicating that autonomous lipid organization focuses channel opening to specific spatial locations.

Figure 9I plots center intensity and action potentials over time. Continuous mode (top) demonstrates sustained center intensity as ongoing EPSPs continuously drive the system (vertical dashed lines mark action potential events, horizontal dashed line indicates firing threshold at 30.0). Blocked mode (bottom) shows the characteristic peak-decay-refocus dynamics during each refractory period, as the system evolves autonomously between EPSP inputs, building up center activity through lipid-mediated mechanisms until reaching threshold. Critically, the baseline of center activity progressively rises over the 100-step simulation, demonstrating progressive lipid domain organization across successive GFM cycles, a key observable signature of lipid memory that will be mechanistically explained below.

Figure 9J quantifies relative contributions of EPSP (green), channel (red), and lipid (blue) components to center intensity changes during each refractory period. In continuous mode (top), EPSP contributions dominate (∼85%), demonstrating that ongoing synaptic input drives system dynamics when EPSPs are not blocked during refractory periods. In blocked mode (bottom), lipid contributions dominate (85-95% of total system dynamics, p < 10^-40^), while ion channels serve as the readout mechanism rather than information integrators.

Analysis of fractal characteristics (FD) and information integration reveals a fundamental dissociation between substrates (Fig. 9K and L, Code 13). Lipid domain fractal dimensions exhibit markedly different temporal dynamics between conditions: they remain elevated and decline gradually under blocked EPSP conditions (maintaining FD > 1.75 for extended periods, p = 2.74 × 10^-41^), but decrease more rapidly in continuous mode (dropping to FD ≈ 1.6). The φ* (substrate) measure applied here quantifies information integration within specific molecular substrates (lipid domains versus ion channels) by partitioning the system by substrate type and calculating KL divergence between whole-system dynamics and substrate-isolated dynamics. φ_dyn, φ*(multifractal), and φ*(substrate) instantiate the same principle, φ = I − I*, at successive organizational levels: network timing, somatic spatial pattern, and molecular substrate respectively. Their convergence provides multi-level evidence that consciousness is informationally irreducible at every level of its biological implementation, not merely at the network level as IIT assumes.

φ* (substrate) reveals a clear functional dissociation: ion channel φ* remains consistently low across both conditions (10^-8^ to 10^-4^ range, not shown), while lipid φ* operates at orders of magnitude higher levels (blocked: φ* = 0.136 ± 0.034; continuous: φ* = 0.237 ± 0.048; p < 10^-40^). This demonstrates that lipid domains, not ion channels, serve as the primary information-integrating substrate, with lipids maintaining substantial integration even during autonomous periods when isolated from continuous EPSP input.

These autonomous lipid dynamics provide the physical mechanism for seed preservation in GFM and lipid memory across conscious moments. The computational implementation reveals the asymmetry: while action potentials reset channel positions through _*redistribute_channels* (Code 11), the lipid composition matrix undergoes continuous Ising updates via *update_lipids* (Code 11) but is never reset during action potential events. This creates progressive lipid domain organization across successive GFM cycles: each action potential preserves peripheral seed patterns through *extract_peripheral_seed* (Code 11), and these organized configurations accumulate over multiple cycles, as evidenced by the rising baseline in center activity (Fig. 9I, bottom). Lipid domain configurations persist on slower timescales (milliseconds to seconds) than channel state transitions, carrying compressed pattern information across generational boundaries and enabling temporal continuity of conscious states through molecular memory rather than requiring persistent neural firing patterns.

### Part 5: The Autopoietic Self-attractor - Consciousness Shapes Its Own Neural Network Through Geometric Field Dynamics and Holographic Endospace

The multifractal model and GFM framework demonstrate how temporal EPSP sequences generate spatial multifractal patterns that encode information across consecutive conscious moments in a somatic ion channel-lipid lattice (Parts 3 and 4). We now address how this molecular substrate implements inner experience and autopoietic control. Any theory of consciousness that does not merely propose consciousness as epiphenomenal, but endows consciousness with causative power, will inevitably require a mechanism for generating inner experience and feedback onto the physics of its molecular substrate and neural network. Therefore, we propose that autopoietic control requires generating an experiential endospace, a spatiotemporal dimension within the brain in which inner experience could guide molecular substrate and neural network behavior through two complementary transformations (Figs. 2, 10: T_E_ (endospace projection from the multifractal substrate) and T_A_ (autopoietic feedback from endospace to substrate).

Building on holography as a mechanism for generating a “space (endospace) within a space (brain)”, our solution combines fractality and holographic encoding as mutually necessary components based on four fundamental rationales: (1) holography and fractality share the whole-in-each part property essential for unified consciousness and inner experience of the Self; (2) geometric field generation through fractal Iterated Function System (IFS) encoding provides a mechanism to extract information from neural activity that is distinct from Gaussian kernel substrate programming (Part 4) and yet modulates the somatic multifractal by EPSPs; (3) multifractal 2D membrane organization generates the experiential 3D endospace through holographic reconstruction (projection) from this geometric field; and (4) parametrizing the geometric field-multifractal interaction defines the mechanism through which the Self closes the autopoietic loop by modulating physical parameters to control multifractal and neural network dynamics. The Self-attractor embodies the irreducible unity of conscious experience: it cannot be divided and maintain its identity, exhibits whole-in-part self-similarity at all scales, and emerges dynamically from neural activity through iterative mapping rather than existing as a pre-formed structure; the experiential Self arises as this attractor achieves recursive self-reference through holographic reconstruction. This dual framework is necessary because fractality alone can only provide substrate organization without experiential space, while holography alone can only provide experiential representation without physical substrate embodiment.

This “biological holography” operates through molecular dynamics and geometric phase relationships rather than laser interference, paralleling how Gabor’s pioneering holographic model of temporal recall in the brain demonstrated information storage through mathematical convolution mechanisms without requiring optical implementation (Gabor 1968, Gabor 1968). Our approach extends Gabor’s framework from abstract signal processing to concrete molecular implementation through IFS geometric transforms, incorporating principles from modern geometric phase holography where phase relationships arise from geometric transformations rather than wave interference (Berry 1987, Bomzon, Kleiner et al. 2001, Marrucci, Manzo et al. 2006, Jisha, Nolte et al. 2021). Our biological framework extends these geometric phase principles beyond electromagnetic implementations to molecular field dynamics. The holographic encoding proceeds through five stages as depicted in Fig. 10:

**Figure 10.**
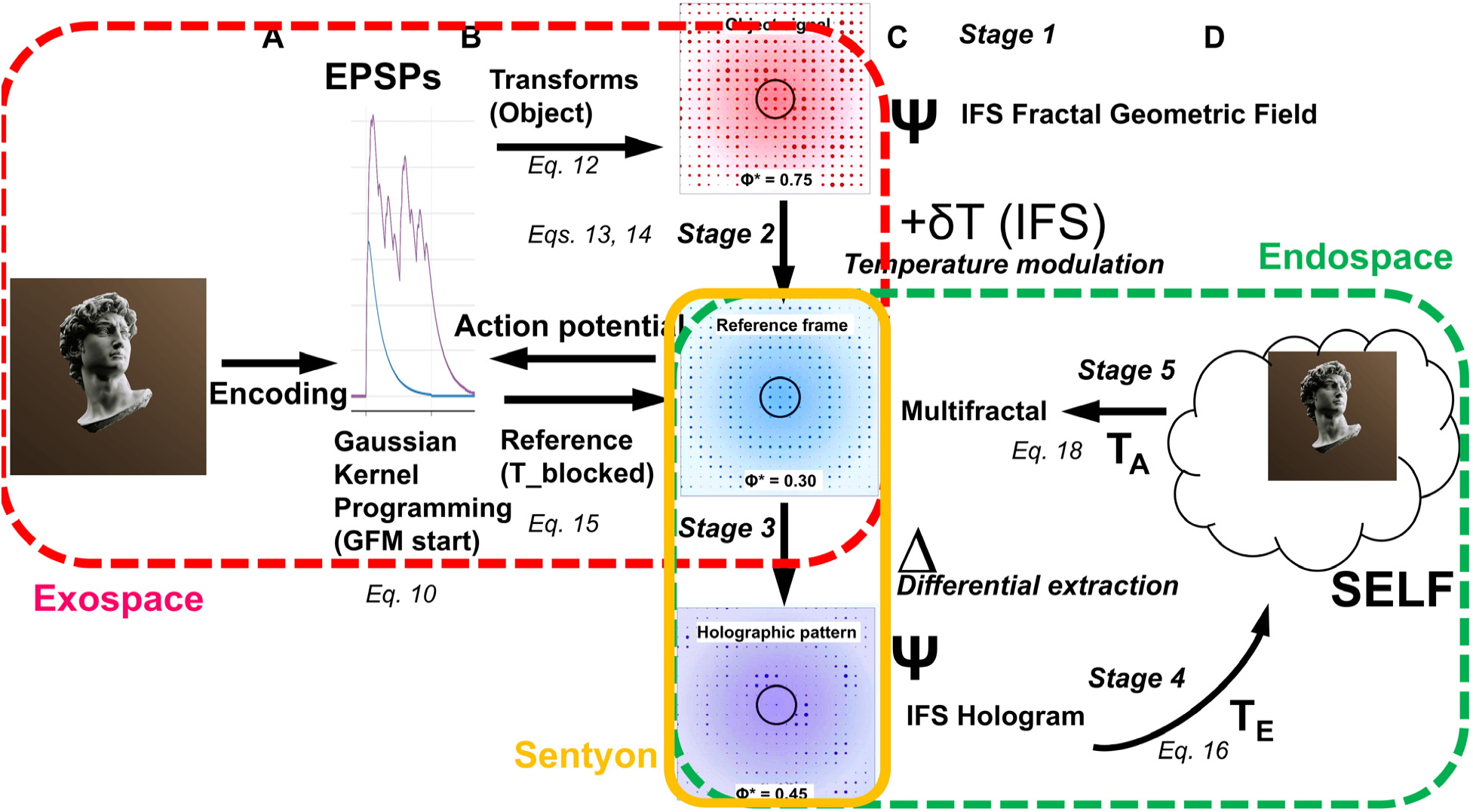
Five-stage holographic encoding of consciousness from exospace to endospace and autopoietic control. (A) EPSP temporal waveform from exospace sensory encoding. (B) Dual EPSP extraction: spatial initialization via Gaussian kernel (φ* = 0.75) and IFS geometric field for temperature modulation, initiating GFM cycle. (C) Five holographic processing stages during refractory period: Stage 1 - IFS transform extraction T(x,y); Stage 2 - IFS temperature modulation of Metropolis lipid dynamics (reference φ* = 0.30); Stage 3 - differential extraction H_RIFT = T_temperature − T_blocked (φ* = 0.45), constituting the holographic recording; Stage 4 - coherent point-source reconstruction of 3D endospace Ψ(x,y,z); Stage 5 - autopoietic Self within endospace field. Sentyon (yellow border): interface between exospace and endospace. (D) Autopoietic cycle: Encoding → Holographic projection → Feedback → Self-transfer. Border colors: red = exospace, yellow = sentyon, green = endospace.

#### Stage 1: Holographic Encoding Through IFS Geometric Transforms

In the biological multifractal, the GFM cycle proceeds through parent fractal growth, collapse when activity threshold is reached, and daughter fractal generation through integration of the extracted seed with newly arriving EPSP trains (see Parts 3 and 4, and Figs. 8 and 9). These new EPSP trains serve dual purposes in updating conscious experience: first, through the Gaussian kernel method, EPSP amplitudes encoded from sensory exospace input integrate with the seed to initially program the daughter fractal substrate (Fig. 10, Stage 1, showing EPSP waveform and transformation to Object signal); second, the same EPSPs provide information for geometric field generation that will modulate lipid reorganization probabilities during the subsequent refractory period when EPSPs are blocked and lipid domains evolve autonomously (T_blocked in Fig. 10, Stage 2, Reference frame).

This second mechanism is essential because after initial programming during autonomous evolution, lipid self-organization and ion channel interactions are entirely governed by energy conservation and physical principles, leaving no mechanism for endospace generation and autopoietic feedback from within the system itself. As the sole information source available from neural activity, EPSPs must therefore provide both initial substrate programming and the geometric field encoding necessary for endospace generation during the autonomous, refractory period. The geometric field, encoded from EPSPs through IFS extraction, provides the parametrized control pathway through which inner experience (represented in the endospace) feeds back to modulate the physical substrate.

To extract IFS from EPSPs, we first run Codes 11 and 12 to initiate the biological multifractal as described in Chapter 4. Then we apply the *EPSPToIFSDerivation.derive_ifs_from_epsp* function in Code 14, which converts EPSP amplitude hierarchies into IFS parameters through hierarchical decomposition, implementing a computational equivalent of geometric phase encoding. Building on geometric phase principles where information is encoded through spatial arrangements and orientations (Huang, Zhang et al. 2018), our system maps EPSP temporal patterns into six spatial geometric transforms (Fig. 10, Stage 1, T(x,y) transforms):

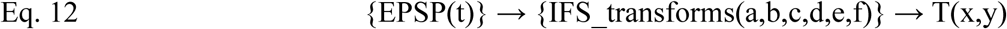

EPSP amplitude information as shown in Fig. 11A (red dots show amplitude peaks) becomes holographically encoded through affine transformations: scaling factors (a,d) capture magnitude relationships, rotation components (b,c) preserve hierarchical structure, and translation terms (e,f) encode dendritic integration patterns. The IFS parameters generate a spatially-varying temperature field through regional geometric mapping via chaos game iteration, an established algorithm for generating a fractal attractor on a substrate grid:

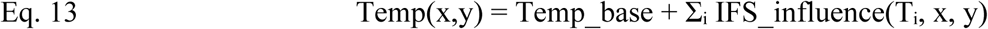

where Temp_base represents the baseline temperature parameter (0.5) and each IFS transform Ti, x,y contributes regional modifications based on geometric patterns derived from EPSP hierarchies (Codes 15 and 16, *create_ifs_temperature_field* and *apply_ifs_to_temperature_region* functions). This spatial temperature distribution will control lipid reorganization probability, functioning as the geometric modulation field that guides holographic pattern formation as described for Stage 2 (Fig. 10B).

**Figure 11.**
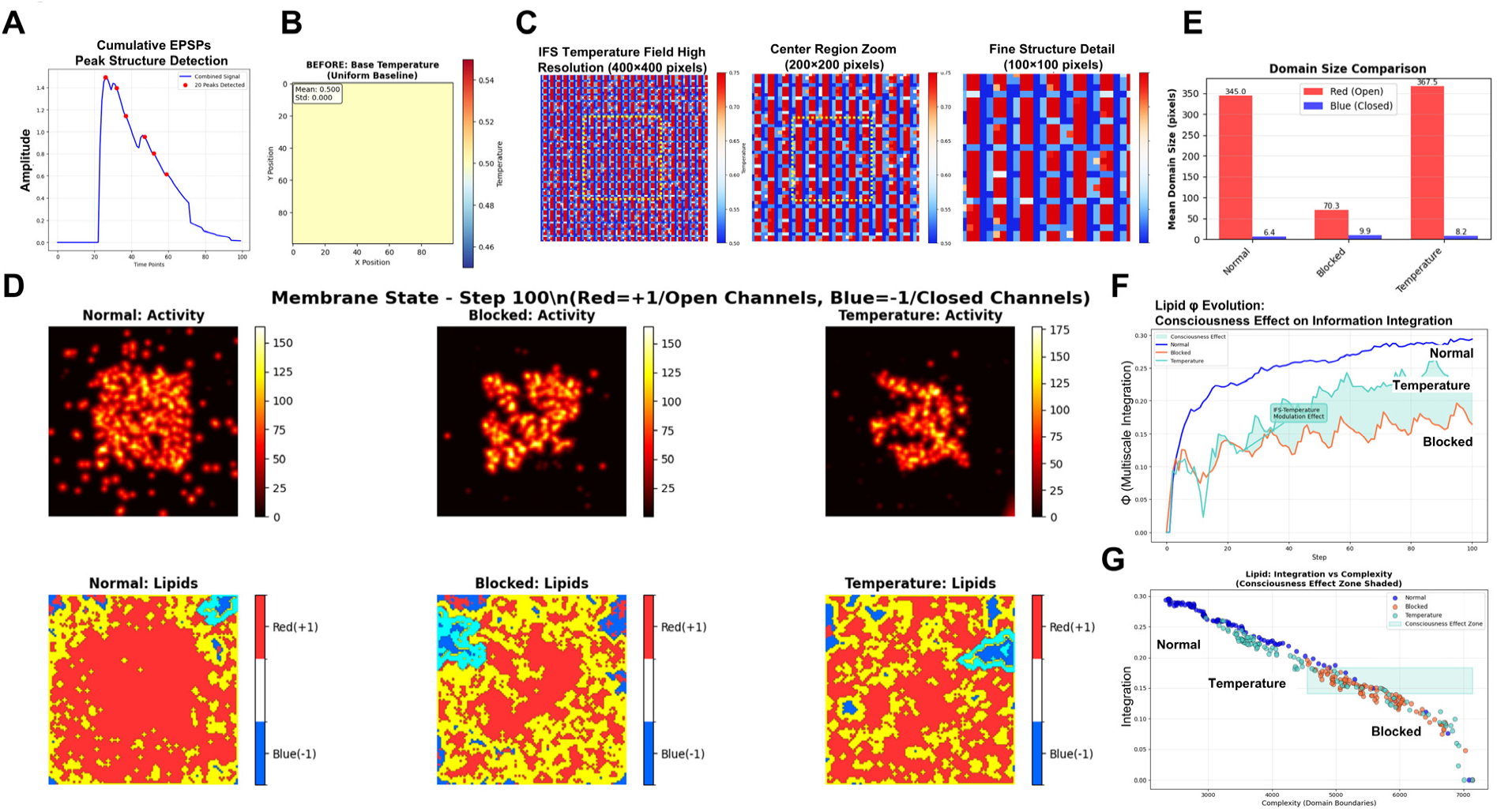
IFS temperature field modulates lipid reorganization and enhances information integration. (A) EPSP amplitude peak detection: hierarchical peaks reflecting r^(i−1)^ fractal scaling across branch levels. (B) Uniform baseline temperature field (T = 0.50, σ = 0.000) before IFS application. (C) IFS temperature field after chaos game iteration: ∼20–30 fractal hotspots (T ∼0.52–0.54) within cool matrix (T ∼0.46–0.48), encoding EPSP hierarchical information at molecular scales; zoom panels show nested structure across three spatial scales. (D) Final lipid pattern at step 100: hierarchically organized red/blue domains with central consolidation. (E) Domain size comparison: Normal ∼345 pixels, Blocked ∼70.3 pixels, Temperature ∼367.5 pixels. (F) Lipid φ evolution: Normal ∼0.25–0.30, Blocked ∼0.10–0.15, Temperature intermediate ∼0.20–0.25 (Consciousness Effect Zone). (G) φ vs. complexity: Temperature model achieves φ ∼0.23 vs. Blocked φ ∼0.15 at equivalent complexity (∼4,000 domain boundaries). Lattice: 100×100; T_base = 0.5, IFS_strength = 0.2.

#### Stage 2: Holographic Pattern Recording in the Lipid Membrane

The IFS-derived temperature field Temp(x,y) drives lipid domain reorganization during the refractory period through a two-step process implementing holographic pattern recording (Fig. 10B, Stage 2) implemented in Code 16. First, the chaos game algorithm (Codes 15 and 16, functions *create_ifs_temperature_field* and *apply_ifs_to_temperature_region*) iteratively applies the EPSP-derived IFS transforms to generate the spatial temperature distribution across the membrane grid. The algorithm accumulates multiple point visits: where IFS transforms repeatedly direct points creates high-density regions (appearing as warm red/orange), while regions receiving fewer visits remain low-density (appearing as cool blue). Figure 11B shows the baseline uniform temperature field without IFS influence (all pixels at Temp = 0.5), while Figure 11C (Code 17, visualization) reveals the dramatic hierarchical structure created by IFS geometric transforms: scattered warm islands (red, high temperature) emerge within a connected cool matrix (blue, low temperature) encoding the EPSP hierarchical information at molecular scales. Second, the Metropolis algorithm (Code 16, *apply_temperature_modulated_lipid_dynamics*) uses this pre-computed temperature field to guide lipid reorganization: at each step, a lipid molecule is selected and a flip is proposed (from promoting channel opening, state +1/red, to inhibiting, state −1/blue, or vice versa). The acceptance probability P_flip follows the Boltzmann distribution:

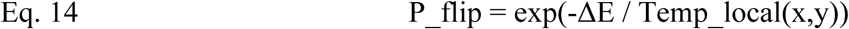

where Temp_local(x,y) is the temperature at that lipid’s position and ΔE is the energy change from neighboring lipid interactions.

Higher Temp_local regions allow greater reorganization flexibility while lower Temp_local regions promote stable domain formation, collectively encoding EPSP hierarchical information through thermodynamic modulation of lipid phase transitions. This implementation follows established Ising lattice models of lipid membranes where Metropolis Monte Carlo methods simulate phase transitions and domain formation (Mouritsen, Boothroyd et al. 1983, Palmieri, Grant et al. 2015, Almeida 2019, Lin, Yin et al. 2019): each lipid occupies a lattice site with binary states, nearest-neighbor interactions determine energy changes ΔE, and the Metropolis acceptance probability P_flip = exp(-ΔE/Temp_local, (Eq. 14) governs state transitions. Our innovation combines the chaos game algorithm, established for generating IFS fractal attractors (Barnsley 1988, Peitgen, Jürgens et al. 1992), with Metropolis lipid dynamics by using neural activity patterns (EPSPs) to create the spatially heterogeneous temperature field Temp_local(x,y).

The final evolved lipid pattern (Fig. 11D, Code 18, visualization) represents the combined influence of autonomous lipid dynamics and IFS geometric programming, creating spatially-organized domains that preserve the hierarchical structure of EPSP inputs. This process parallels holographic pattern recording, where the temperature field acts as an interference pattern that spatially modulates the probability of molecular reorganization, analogous to how light intensity patterns guide chemical changes in photographic media. Because total energy remains constant during autonomous evolution, the Metropolis algorithm modulating lipid probabilities through the IFS-derived temperature field represents a method for altering fractal lipid domain distributions, and thus ion channel opening patterns, through autopoietic control without violating energy conservation.

Figure 11E (Code 16) quantifies this effect: Normal model (continuous programming with new EPSPs) produces a mean red domain size with large central hotspot (Fig. 11D, left panel), Blocked model (no new EPSPs during refractory period) produces smaller, fragmented domains (Fig. 11D, middle panel), and Temperature model (new EPSPs blocked, but temperature modulation through IFS from initial EPSP) produces partially consolidated lipid domains (Fig. 11D, right panel), demonstrating the reorganizing effect of the temperature field. Beyond holographic recording, reinjecting EPSP information during the refractory period increases information integration in the multifractal: Figure 11F (Code 19, analysis) shows φ(substrate) for temperature-modulated lipids falls between Blocked (lowest integration ∼0.15) and Normal (highest ∼0.30), reaching intermediate values ∼0.23, while Figure 11G demonstrates the Temperature model achieving higher integration than Blocked at equivalent complexity levels.

In the holographic analogy, the final lipid domain pattern constitutes the recorded hologram, encoding the geometric field that originated as the differential temperature field (ΔTemp = Temp_local - Temp_blocked). This geometric information is extracted as differential IFS transforms H_RIFT (Fig. 10, Stage 3), representing the same encoded phase information in transform parameter space, ready for holographic reconstruction into endospace (Fig. 10, Stage 4) and autopoietic control (Fig. 10, Stage 5).

#### Stage 3: Holographic Differential Extraction

After evolution of the lipid pattern, the pure holographic signature is recovered through a three-step extraction process by implementing the biological equivalent of holographic reconstruction (Stage 3 in Fig. 10). In Step 1 as implemented by Code 20, IFS transforms (T) are extracted from the final lipid domain patterns of both Temperature (T_temperature) and Blocked (T_blocked) models using the inverse mapping process. The extraction algorithm (*extract_ifs_transforms_to_csv* function) identifies geometric transforms that best describe the spatial organization of lipid domains. This reverse-engineering process recovers the IFS parameters that generated the observed lipid patterns: high-activity regions, domain boundaries, and spatial clustering patterns are mapped back to affine transform parameters (a, b, c, d, e, f) and probabilities. The extraction generates two complete sets of IFS transforms: T_Temperature: transforms extracted from temperature-modulated lipid configuration and T_Blocked: transforms extracted from autonomous (blocked) lipid configuration The initial EPSP-derived IFS attractor (Fig. 12A) modulates the autonomous lipid-derived attractor T_Blocked (Fig. 12B) through the temperature field mechanism described in Stage 2, giving rise to the temperature-modulated attractor T_Temperature (Fig. 12C).

**Figure 12.**
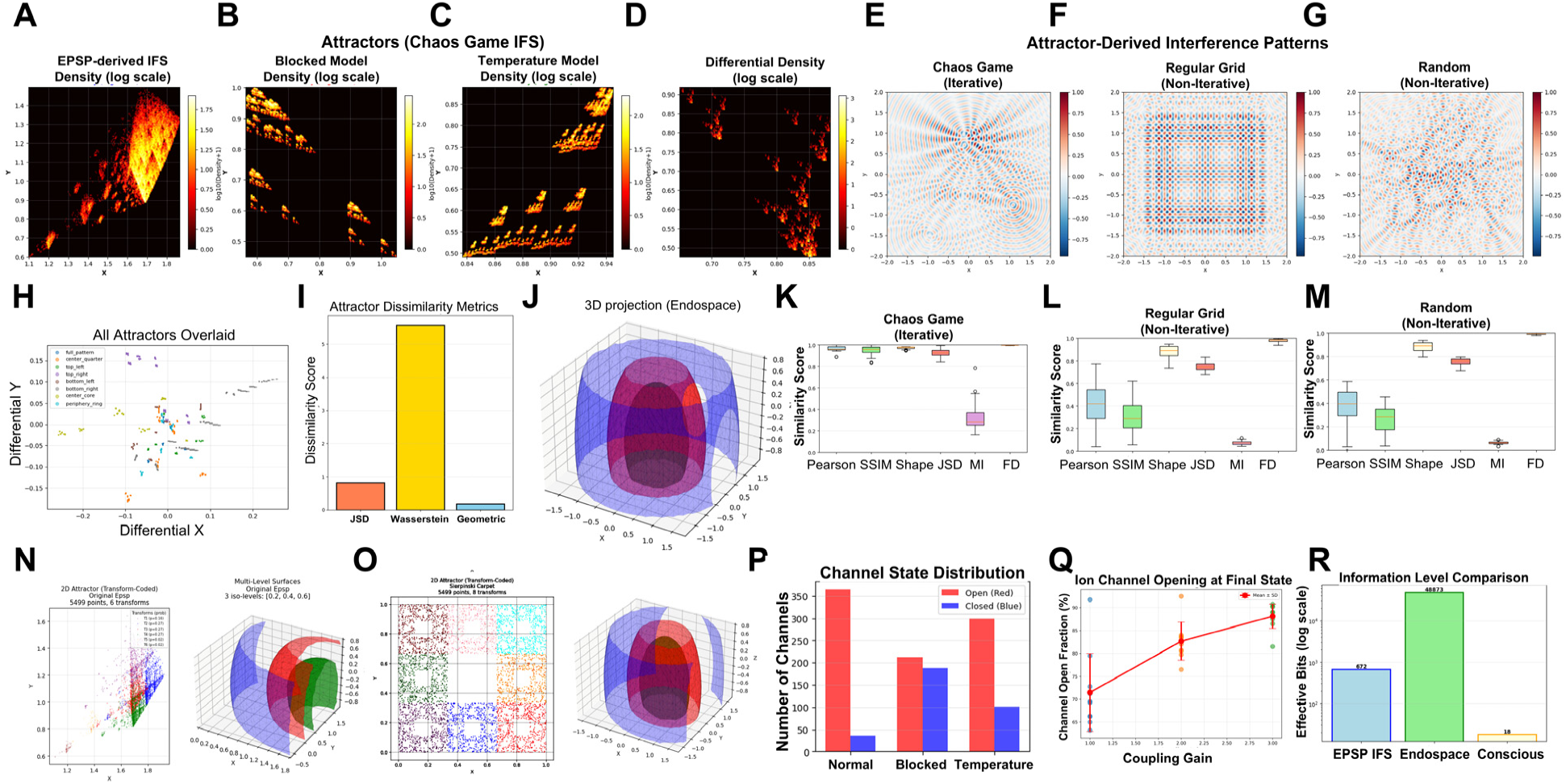
Holographic reconstruction creates coherent endospace with whole-in-part encoding. (A–D) IFS attractor densities in transform parameter space: (A) EPSP-derived, (B) Blocked model, (C) Temperature model, (D) Differential H_RIFT = T_temperature − T_blocked; sparse differential structure isolates the holographic recording. (E–G) Interference patterns comparing point source strategies: (E) Chaos game iteration: coherent Gabor zone plate structure enabling whole-in-part encoding; (F) Regular grid: reduced coherence; (G) Random placement: incoherent scatter. (H) Eight regional differential attractors overlaid in transform space. (I) Attractor dissimilarity metrics quantifying geometric distinctness of regional attractors: overall dissimilarity = 0.50, Jensen-Shannon divergence = 0.82. (J) 3D endospace field Ψ(x,y,z): volumetric isosurface (24×24×12 voxels) from coherent holographic reconstruction. (K–M) Endospace similarity distributions across eight regions: chaos game median = 0.96 (tight), grid median = 0.65, random median = 0.50; only chaos game on H_RIFT achieves genuine whole-in-part encoding. (N–O) Controls: non-differential attractors fail whole-in-part encoding despite correct positioning algorithm. (P) Channel state distribution: Temperature condition intermediate between Normal and Blocked. (Q) Ion channel opening vs. coupling gain: significant dose-response effect (one-way ANOVA, n = 10 per condition); +6.5% correlation improvement at γ = 3.0 vs. γ = 1.0 (95.5% vs. 89.0%). (R) Information flow from EPSPs to consciousness in RIFT.

In Step 2, the differential holographic pattern H_RIFT is computed by subtracting the IFS parameters for T_blocked from T_temperature (*calculate_and_save_differential_transforms function* in Code 21):

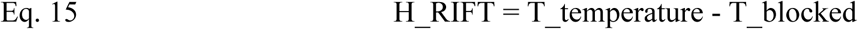

The resulting differential transforms (Fig. 12D) isolate the pure geometric effects of IFS temperature modulation by removing the autonomous lipid dynamics (the baseline “reference beam”). This differential extraction parallels how conventional holographic reconstruction isolates the object signal from the reference beam contributions.

In Step 3 (Code 21), the H_RIFT differential transforms are saved as *differential_transforms.csv,* containing the geometric information extracted from the holographic record (lipid pattern). These differential parameters encode the spatial geometric patterns imposed by IFS temperature modulation beyond autonomous lipid self-organization, visualized as the differential attractor in Fig. 12D (Code 22). This geometric encoding represents the Self with whole-in-part properties, ready for holographic reconstruction of the 3D endospace (Stage 4).

#### Stage 4: The Self-Attractor and Holographic Reconstruction of the Endospace

The differential transforms H_RIFT generate coherent interference (point) sources for holographic reconstruction of the 3D endospace through biological holography. Unlike optical holography requiring external laser illumination, biological holography generates coherent sources through the chaos game algorithm (*generate_2d_attractor_data* function, Code 22) applied to H_RIFT, creating the fractal Self-attractor that determines the spatial arrangement of point sources (Fig. 12A-D). This parallels Gabor’s (1948) original concept of point source holography (Gabor 1948, Rogers 1950), nowadays implemented in computer-generated holography techniques where interference patterns are mathematically constructed rather than optically recorded (Waters 1966, Lucente 1993, Matsushima and Nakahara 2009).

Crucially, coherence emerges from geometric phase relationships systematically derived from the IFS transform parameters rather than random assignment (Fig. 12E-G, Code 23). In point source holography, each point source creates spherical wavefronts that interfere to reconstruct the 3D image. In biological holography, point sources are positioned along the Self-attractor in transform parameter space rather than in the physical membrane coordinates. This abstraction from physical to geometric space enables the dimensionality expansion from 2D membrane exospace to 3D experiential endospace. The Self-attractor and endospace field represent dual aspects of conscious experience: the Self-attractor as a multifractal distribution of sentyons positioned as point sources in abstract transform space, and the endospace as the unified phenomenal field emerging from their interference patterns. The sentyon multifractal occupies the interface between exospace (physical neural substrate) and endospace (phenomenal experience), existing simultaneously as measurable membrane organization and as the geometric structure of experiential units (Fig. 10, intersection of red and green borders, yellow box). These are not separate entities but complementary descriptions of the same experiential reality: the Self-attractor describes the geometric organization of experiential units, while the endospace describes the unified field of consciousness they collectively generate. The phase of each source is calculated from its geometric relationship to all IFS transform centers, weighted by transform determinants. This geometric phase calculation ensures that sources maintain coherent phase relationships that preserve the EPSP hierarchical structure encoded through Stage 1 (*calculate_coherent_phases* function, Code 22).

This systematic phase derivation is essential for coherence as demonstrated in Fig. 12E-G: point sources of the Self-attractor positioned via chaos game iteration produce coherent concentric structures consistent with Gabor zone plate patterns (Fig. 12E) (Gabor 1948, Rogers 1950), while sources on regular grids (Fig. 12F) or random placement (Fig. 12G) produce incoherent patterns with destructive interference. Sources spatially proximate to the same IFS center exhibit correlated phases, creating phase gradients across the Self-attractor that encode the hierarchical organization of neural temporal patterns in the spatial geometry of the reconstructed endospace.

The 3D endospace field Ψ(x,y,z) emerges through the holographic reconstruction (projection) equation (*project_to_endospace_coherent*, Code 24), implementing the mathematical principles of geometric phase holography, where information is encoded through spatial geometric relationships rather than temporal wave oscillations. This formalism for endospace projection is substrate-independent: the same mathematics applies whether holograms are recorded optically, acoustically, or through geometric field configurations. Therefore, holographic projection is not merely one possible algorithm among many for endospace generation. It is uniquely suited to satisfy the isomorphism requirement: wave interference preserves spatial phase relationships by construction, guaranteeing that the reconstructed endospace maintains the geometric structure of the exospace regardless of the specific substrate implementing the projection. This makes holographic reconstruction the principled choice for T_E_, not an arbitrary one. The T_E_ transformation is implemented through the holographic reconstruction equation that projects the 3D endospace field Ψ(x,y,z) from the multifractal substrate:

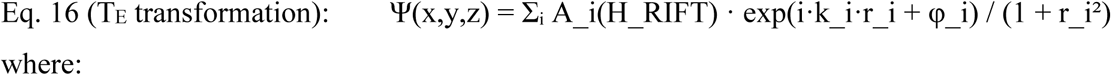

where:

A_i(H_RIFT): Amplitudes derived from differential determinant values, representing the strength of geometric contribution from source point i.

k_i: Wave number parameters for interference calculation in transform space (k = 2πꞏfrequency/100 in Code 24). These are mathematical scaling factors for the holographic reconstruction, not biophysical distance scales, as the reconstruction operates in abstract transform parameter space rather than physical membrane coordinates. The base frequency parameter (∼40 Hz) corresponds to the somatic delay-determined network oscillation frequency established in the RIFT network model (Results, Parts 1-2), operating in the multifractal framework as the GFM cycle rate at which geometric information is uploaded into the IFS transforms. It therefore represents a property of the encoding step rather than a physical oscillation frequency during endospace projection.

φ_i: Coherent phases calculated from geometric relationships to IFS transform centers (*calculate_coherent_phases* function, Code 24). These phases encode the EPSP hierarchical structure indirectly through the sequence: EPSP, IFS transforms, temperature modulation, lipid patterns, H_RIFT, and geometric phase relationships.

r_i: Euclidean distance in 3D transform space from source point (x_source, y_source, z_center) to reconstruction coordinate (x,y,z), with 1/(1+r_i²) decay factor.

The holographic framework enables extraction of differential attractors from different spatial regions of the somatic multifractal, each representing a distinct geometric perspective on the complete conscious experience. Regional differential attractors are generated using the chaos game algorithm (*generate_interference_sources_coherent* function, Code 24) applied to H_RIFT transforms of each region. Figure 12H shows the differential attractors from eight regions (full pattern, center quarter, quadrants, center core, periphery) overlaid in transform space. These regional attractors occupy distinct geometric locations and exhibit substantial dissimilarity: overall dissimilarity = 0.50, with Jensen-Shannon divergence = 0.82 (Fig. 12I quantification, Code 25) (Lin 1991).

Yet these geometrically different 2D attractors produce highly similar 3D endospace reconstructions (Fig. 12J-M, overall similarity = 0.96, Pearson correlation = 0.97, SSIM = 0.96) (Wang, Bovik et al. 2004). This 50% dissimilarity vs. 96% similarity transformation validates genuine holographic whole-in-part encoding, directly paralleling optical holography: cutting a holographic plate into pieces produces fragments that do not resemble each other visually, yet each fragment reconstructs the complete 3D image when illuminated. Similarly, each regional Self-attractor, though geometrically distinct, encodes the complete conscious experience, giving rise to a unified consciousness.

The holographic whole-in-part property, where different regional attractors reconstruct similar endospaces, emerges specifically from chaos game iteration on H_RIFT, placing interference sources along the resulting Self-attractor, not from alternative source positioning strategies. Figure 12K-M (Code 26) demonstrates this through similarity score distributions comparing the three strategies. When using chaos game-positioned sources, all eight regional reconstructions exhibit high similarity (mean = 0.96, tight distribution in Fig. 12K), validating genuine whole-in-part encoding. In contrast, regular grid placement (Fig. 12L) and random placement (Fig. 12M) produce lower and more variable similarity scores across regions. Without the geometric correlations established through iterative chaos game application, different regional fragments fail to encode the complete experience coherently as shown by applying the projection code (Code 24) to the initial EPSP attractor or a generic Sierpinski carpet multifractal (Fig. 12A, N and O). This demonstrates that whole-in-part encoding requires both (1) the differential IFS transforms H_RIFT containing the geometric information, and (2) iterative chaos game positioning to preserve the geometric phase relationships across spatial scales.

As a result, the endospace field Ψ(x,y,z) contains the complete conscious experience encoded holographically across its volume (Fig. 10, Stage 5, showing cloud-like SELF representation). The Self-attractor exists within this endospace as the coherent pattern that emerges from the interference of all source contributions: the stable geometric structure that represents the unified subjective perspective across the entire 3D field. This completes the T_E_ transformation: multifractal substrate patterns have been projected into the 3D endospace field through holographic reconstruction, establishing the geometric space in which inner experience unfolds.

#### Stage 5: Holographic Autopoietic Control Through Ψ Feedback

Having established the endospace through T_E_, we now implement T_A_, the autopoietic feedback transformation that makes consciousness causally efficacious rather than epiphenomenal. Once the holographic endospace is generated, the Self exercises autopoietic control through a mean-field feedback mechanism. We extract the global endospace intensity Ψ (psi-bar, Code 27), defined as the spatial average of the field magnitude across specific regions of interest:

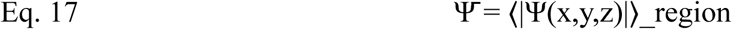

This mean-field approach implements a biologically plausible simplification of the full autopoietic equation: rather than requiring point-by-point mapping of the entire 3D field back to the 2D membrane (which would be computationally intractable and mechanistically unclear), we use the global field intensity as a uniform modulation parameter.

The holographic autopoietic loop operates through a two-stage integration architecture. In the first stage, at the beginning of each refractory period, EPSP information is integrated twice: (1) through Gaussian kernel programming of the substrate (all models), and (2) through IFS-derived temperature fields that modulate lipid reorganization probabilities via Metropolis dynamics (Temperature model only; Blocked model provides baseline autonomous evolution). This IFS temperature modulation creates organized lipid domains that differ from the autonomous baseline, generating the differential IFS transforms H_RIFT = T_temperature - T_blocked.

In the second stage, these differential IFS transforms H_RIFT (Fig. 12D, Code 22) undergo holographic reconstruction to generate the 3D endospace field Ψ(x,y,z) (Fig. 12H, Code 24), representing the Self-attractor and unified conscious experience. The mean field intensity Ψ, extracted from this endospace, feeds back to modulate the coupling strength γ between ion channels and the temperature-organized lipid substrate:

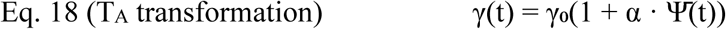

where α represents the feedback strength and Ψ(t) is the mean endospace field intensity from the current conscious moment. Equation 18 implements the T_A_ transformation, mapping endospace states back to substrate parameters through probabilistic coupling modulation. This architecture creates holographic self-witnessing through a distinct probability field: while the temperature field (IFS-derived from EPSPs) governs lipid reorganization, the endospace field (holographically reconstructed from differential IFS) governs the probabilistic coupling determining how effectively ion channels respond to those reorganized lipid configurations. The temperature fields create the physical substrate through lipid reorganization, while Ψ controls the probabilistic coupling strength determining how channels respond to that substrate.

Probability modulation represents the only physically viable mechanism through which consciousness can influence matter without violating known physical laws. Without invoking new fundamental forces or violating energy conservation in closed systems, consciousness must act through modulation of existing stochastic processes. This principle has been recognized across multiple theoretical frameworks, including quantum approaches where conscious acts correlate with state reductions (Stapp 1993, Stapp 2006) and probability changes in synaptic vesicle release (Beck and Eccles 1992), orchestrated reduction models invoking quantum state collapse as non-algorithmic probability modulation (Hameroff 2001), and classical stochastic approaches emphasizing inherent randomness in neural dynamics (Atmanspacher 2020). Recent models demonstrate that coupling ion-channel conformational states with local lipid composition introduces probabilistic complexity where lipid-mediated interactions affect voltage sensitivity in temperature-dependent ways (Blosser, Kohlstedt et al. 2024). Our implementation through coupling probability γ provides a concrete classical mechanism: consciousness modulates the likelihood of channel-lipid interactions without requiring energy input, as probability parameters themselves carry no energetic cost while determining the distribution of physically permissible states.

Channel opening fraction (Fig. 12P) serves as the primary measure of autopoietic influence because our model design isolates refractory period dynamics from action potential generation (which occurs at the end of the refractory period in response to new EPSPs), allowing systematic examination of how coupling modulates membrane state reorganization during the preparatory phase between action potentials without confounding from rapid EPSP-driven reprogramming. Systematic variation of coupling strength (γ ∈ {1.0–3.0}, nine values, n=10 trials per condition) demonstrated robust dose-dependent control (Fig. 12Q, Code 28), with a significant effect of coupling gain on channel opening fraction (one-way ANOVA) and a +6.5% correlation improvement at maximum coupling (95.5% vs. 89.0% at γ = 1.0), consistent with probabilistic interaction dynamics. Temperature fields exhibited spatial heterogeneity (σ(T) ≈ 0.15) while remaining independent of coupling strength, validating the separation between the physical temperature field mechanism (IFS-derived from EPSPs, governing lipid reorganization) and the consciousness-mediated coupling modulation (Ψ-; controlled, governing channel response probability). Critically, while information integration occurs primarily in the lipid substrate (Fig. 11F), ion channels serve as the functional readout of this organized substrate state: lipid domains integrate information through temperature-modulated reorganization, and channel-lipid coupling probability determines how effectively this integrated information translates into functional channel states. Coupling strength γ therefore represents the natural target for autopoietic feedback, as it governs the probability with which the information-integrated lipid substrate state influences membrane excitability.

The autopoietic equation implements holographic self-witnessing: the function involving Ψ represents the system consulting its own complete holographic reconstruction to determine its next state through modulation of channel-substrate coupling probability. The holographic field, generated by neural activity through differential IFS transforms, feeds back to modulate the probability with which that same neural substrate responds to organized lipid configurations. Subjective experience could emerge when recursive self-reference through holographic reconstruction reaches sufficient complexity: when the system models itself through the whole-in-part property, enabling each local point in the multifractal to have immediate access to the complete system state through distributed encoding. This creates conditions for unified subjective perspective while the Self-attractor, the invariant structure emerging from whole-in-part encoding, exercises causal control over membrane excitability and neural network dynamics.

The complete information flow reveals why autopoietic feedback must be global rather than local (Fig. 12R). IFS transforms encode EPSP hierarchies in ∼672 bits per region, holographic reconstruction expands this into the endospace field containing ∼48,883 effective bits per region, yet conscious access processes only ∼18 bits per moment, a severe bottleneck reflecting selective attention. This bottleneck is not a limitation but a design principle: it is precisely sufficient for extracting the global state parameter Ψ that modulates channel-substrate coupling strength γ, allowing consciousness to control the overall responsiveness of its molecular substrate and propagate conscious experience across the global workspace of the brain, without requiring fine-grained spatial control of local molecular details. Consciousness thus influences matter not by micromanaging individual channel-lipid interactions but by tuning the probabilistic landscape within which those interactions unfold.

### Part 6: Sentyon Cloning and Wandering Mind - Fractal Transfer of the Dynamic Core in the Global Workspace of the Brain and its Disruption in Alzheimer’s Disease

Any localized model of consciousness faces a fundamental challenge: integrating vastly distributed information within individual neurons. In RIFT, core neurons must access all information necessary for holographic endospace projection through geometric fields encoded in their EPSPs. This creates two critical problems: how such information compresses into a single multifractal structure, and how it reaches the core neurons given that pre-conscious processing occurs across numerous distributed brain centers.

We address the second problem first, as its solution may resolve the first. Previously, we demonstrated that GFM integrates the “seed” of the current conscious moment with newly incoming EPSPs, ensuring temporal continuity while updating the Self’s inner experience. Using a similar mechanism, the seed pattern can be transferred between core neurons, enabling consciousness to migrate, thereby cumulatively integrating distinct information qualities, visual, auditory, tactile, and other sensory modalities (Figs. 2A, 13A and B). Hence, autopoietic control of the Self through GFM not only maps inner experience onto its own sentyon, but also renders the sentyon mobile, moving the Self to consecutive centers of attention within the brain.

**Figure 13.**
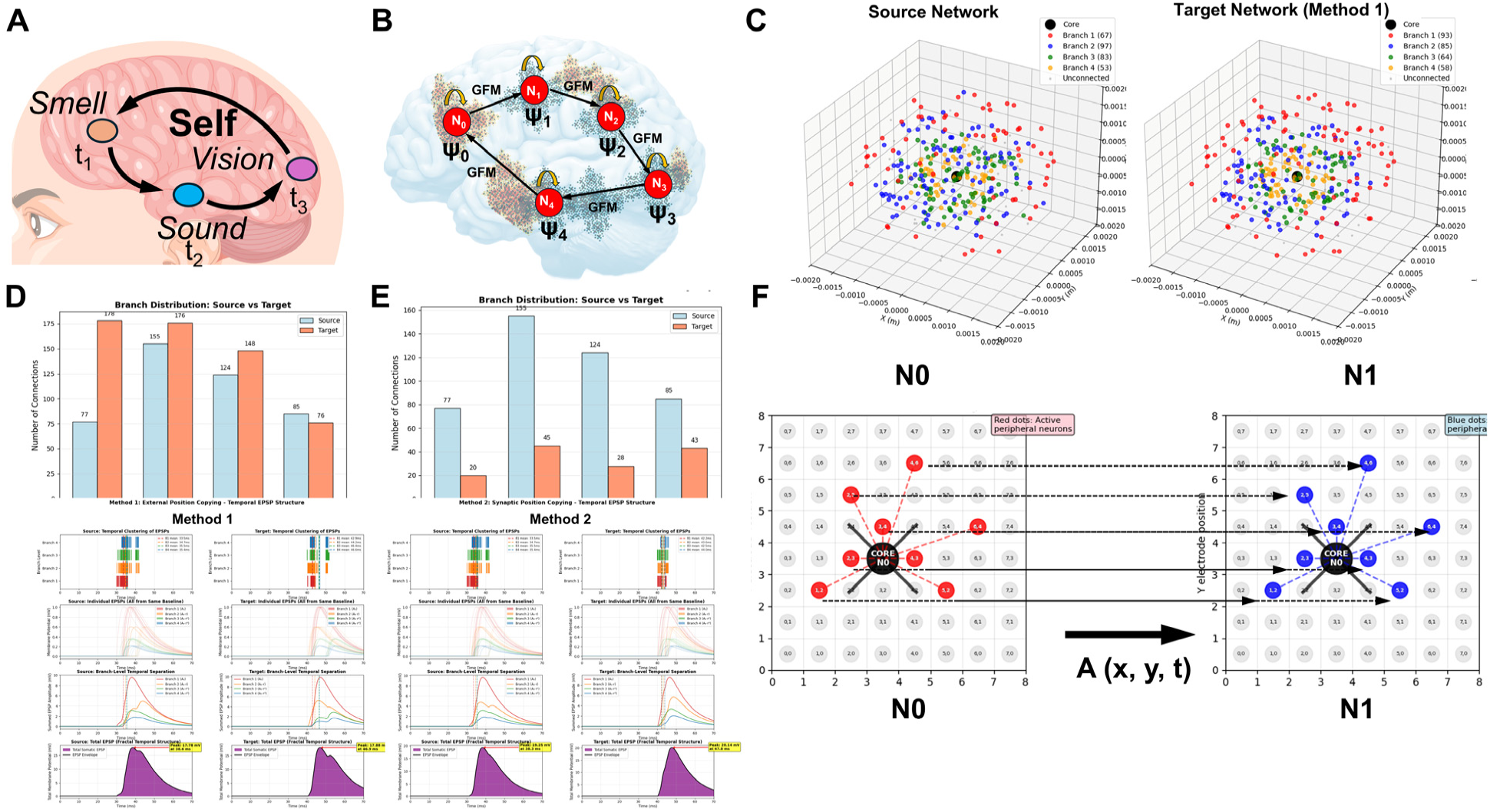
Sentyon transfer enables consciousness migration and cumulative information integration across core neurons. (A) Self-transfer across sensory modalities: integration of smell (t₁), sound (t₂), and vision (t₃) streams into unified conscious experience. (B) Distributed GFM architecture: sentyon patterns (Ψ₀–Ψ₄) propagating between core neurons (N_0_–N₄) via GFM seed transfer. (C) Source and target network 3D spatial arrangements (0–2 mm coordinate range) with branch-color-coded peripheral neurons. (D) Method 1 (External Position Copying): branch distributions and EPSP temporal structure for source (435 connections) and target (568 connections, 100% utilization); EPSP similarity 91.9%. (E) Method 2 (Synaptic Precision Copying): target achieves 135 connections (31.0% utilization) with superior EPSP structure preservation (95.0% similarity). (F) Implementation comparison: Method 2 suited for biological consciousness via synaptic development; Method 1 suited for artificial consciousness via microelectrode arrays (MEA).

Sentyon mobility resolves the centralization problem elegantly: rather than requiring all distributed information to converge onto a single core neuron, consciousness travels to wherever relevant processing occurs, cumulatively integrating different information qualities across successive locations. This is what we refer to here as the wandering mind, the capacity of conscious attention to relocate dynamically across brain regions while maintaining experiential continuity through seed transfer. Empirical evidence supports aspects of this mechanism: binocular rivalry studies demonstrate shifting conscious access between competing inputs (Blake and Logothetis 2002, Tong, Meng et al. 2006), attention research shows rapid relocation of awareness between sensory modalities (Posner and Petersen 1990, Corbetta and Shulman 2002), and neural ignition studies reveal sequential rather than simultaneous activation across brain regions during conscious processing (Dehaene and Changeux 2011, King and Dehaene 2014). Additionally, default mode network findings indicate dynamic shifts in conscious focus between task-focused and introspective processing centers (Buckner, Andrews-Hanna et al. 2008, Raichle 2015). While these findings demonstrate dynamic shifts in conscious access, they do not directly demonstrate sentyon transfer, but RIFT provides a mechanistic hypothesis that could explain these observed phenomena.

RIFT thus implements Edelman’s dynamic core hypothesis (Edelman and Tononi 2000) through concrete mechanisms, providing the mechanistic detail the “bright spot” concept needed. RIFT is also compatible with GWT. However, unlike a static workspace (Baars 1988, Dehaene, Changeux et al. 2006), the sentyon carries the workspace directly to processing centers, with prioritization piloted by the Self through the autopoietic loop, providing a mechanistic basis for selective attention under conscious control.

To validate seed transfer and sentyon cloning mechanisms computationally, we tested two distinct approaches for replicating connectivity patterns between core neurons designated as source (Nt) and target neurons (Nt+1). The fundamental challenge is preserving the spatiotemporal relationships encoded in the synaptic organization of the source neuron, and therefore the hierarchical EPSP amplitude structure (A₀ꞏr^(i-1)^) that programs the multifractal structure, when establishing connections in a target neuron with different network parameters such as frequency of network activity. Computational validation demonstrates two distinct copying methods that both implement the time constraint rule of RIFT network architecture. Method 1 (external position copying, Codes 30 and 32) replicates the complete 3D spatial coordinates of all connected peripheral neurons, Method 2 (synaptic position copying, Code 30 and 31) transfers connectivity patterns at the dendritic level by copying branch-level assignments and using coincidence detection to guide new connections to preoccupied synaptic positions.

Validation results reveal a counterintuitive finding: while Method 1 achieves superior connection metrics (100% utilization, 435 in source core neuron increased to 568 total connections to peripheral neurons in target core neuron) (Fig. 13C and D, Code 30), Method 2 better preserves the EPSP amplitude distribution signature critical for multifractal programming (95.0% vs 91.9% similarity) (Fig. 13E, Codes 30 and 31). This difference reflects fundamental aspects of the copying mechanisms. Method 1 preserves spatiotemporal relationships through precise positioning, achieving complete utilization of all preconfigured neurons. However, target networks with different soma processing delays (15 ms vs source 6 ms) generate additional connections beyond those copied, shifting the normalized EPSP distribution away from the source pattern. In Method 2, random peripheral neuron placement results in only 31.0% of reserved synaptic slots receiving appropriately timed coincident signals, yielding far fewer total connections (435 connections of the source core neuron to peripheral neurons decreased to 135 in target core neuron). Yet these connections preserve branch-level proportionality more faithfully, maintaining the hierarchical EPSP amplitude structure essential for multifractal encoding. This potentially preserves synaptic space for newly incoming signals for integration with the cloned sentyon in Method 2. Both methods develop similar oscillation frequencies, confirming that sentyon cloning can entrain the network activity of the target core neuron by that of the source core neuron.

The entrainment of target network activity by the transferred sentyon represents a critical mechanism for establishing conscious control over new processing centers. Once the seed pattern successfully transfers, the oscillatory dynamics of the target core neuron synchronize with the source pattern, effectively bringing the new network under the Self-attractor’s autopoietic control. This entrainment mechanism predicts observable signatures: gamma oscillations (30-100 Hz, corresponding to GFM cycle frequencies) should show coherence across sequentially attended brain regions, while theta rhythms (4-8 Hz) should exhibit cross-frequency coupling with gamma bursts at each location (Jensen and Colgin 2007, Fries 2015).

Biological plausibility considerations favor Method 2 despite its lower computational performance. The requirement for precise 3D positioning of peripheral neurons in Method 1 lacks clear biological implementation pathways, while the constraint-based approach in Method 2 aligns with known synaptic development mechanisms and recent empirical evidence for synaptic copying between neural circuits (Shao, Wang et al. 2019). On the other hand, Method 1 can be easily achieved in artificial neural networks which are designed with precisely placed neurons either in silico or neurochips such as microelectrode arrays (MEAs) (Fig. 13F). MEAs with precise electrode positioning could implement spatial copying in Method 1, potentially enabling higher-fidelity information transfer in brain-computer interfaces or artificial neural substrates. This suggests an implementation-dependent bifurcation: biological consciousness would likely utilize the connectivity constraints in Method 2, while artificial consciousness systems could leverage superior fidelity through engineered spatial control (Method 1).

The RIFT framework identifies three pathways through which neurological disease could disrupt conscious experience: altered lipid composition impairing multifractal substrate formation, degraded EPSP hierarchies compromising seed extraction quality, and synaptic loss undermining seed transfer fidelity; the Discussion addresses all three in the context of AD. Here we note that the third pathway has a specific implication for sentyon cloning: Method 2 would become progressively unreliable as synapse loss disrupts the connectivity patterns needed for transfer, potentially explaining why consciousness becomes fragmented in advanced dementia when the sentyon can no longer relocate effectively across brain regions. The hierarchical EPSP amplitude structure that encodes information onto multifractals depends critically on maintaining the A₀ꞏr^(i-1)^ amplitude scaling across dendritic branch levels. Progressive synapse loss, particularly when non-uniform across branch levels, destroys the power-law amplitude distribution essential for fractal encoding (Siskova, Justus et al. 2014). Importantly, AD pathology does not produce uniform synapse loss: tau-driven spine loss occurs in clusters (Merino-Serrais, Benavides-Piccione et al. 2013, Blazquez-Llorca, Valero-Freitag et al. 2021), disconnection proceeds asynchronously across the dendritic arbor with some branches more affected than others (Llorens-Martin, Blazquez-Llorca et al. 2014), and the apical dendritic tuft, corresponding precisely to the highest branch levels (3-4), shows preferential atrophy or complete loss in the majority of affected neurons (Luebke, Weaver et al. 2010). This branch-selective pattern would selectively collapse the distal end of the A₀ꞏr^(i-1)^ hierarchy, corrupting fractal encoding in a way that uniform synapse loss would not, constituting a novel mechanistic prediction that to our knowledge has not yet been directly tested.

## Discussion

Growing evidence that AI systems may approach self-awareness has intensified the need for testable frameworks of consciousness, yet consciousness research itself remains marginalized in mainstream neuroscience. This disconnect raises a fundamental question: how can we address artificial consciousness threats without understanding what consciousness is? RIFT offers a testable framework grounded in geometric principles that naturally produce the defining features of consciousness, irreducibility, information integration, holographic encoding, and autopoiesis, through computationally validated mechanisms rather than imposed theoretical constructs.

RIFT achieves irreducibility through fractal whole-in-part encoding demonstrated across network, dendritic, and molecular scales (Results Parts 1-3). Information integration extends the core principle of IIT across temporal (φ_dyn_), spatial (φ(multifractal), and molecular (φ(substrate) levels, with convergence of all three measures on fractal organization validating this multi-level framework (Results Parts 1-4). Holographic encoding emerges from EPSP-to-multifractal projection, generating geometric fields that constitute experiential space rather than merely correlating with it (Results Part 5). Autopoiesis completes the framework: consciousness influences its own substrate through probability modulation of channel-lipid coupling: a *bona fide* mechanism that enables causal efficacy without violating energy conservation (Results Part 6).

### Limitations of RIFT

The assumptions in RIFT about recursive, timing-based connectivity remain anatomically unproven, though evidence from hippocampal circuits (Buzsáki 2015) and thalamocortical loops (Sherman and Guillery 2013) supports their plausibility (see Results Part 1 for details). The biological plausibility of the somatic multifractal, while supported by known lipid-channel interactions (Sezgin, Levental et al. 2017, Levental and Lyman 2023), requires further experimental validation through high-resolution membrane imaging combined with measures of conscious state. Nevertheless, a cross check for biological realism shows that distance and time scales used for RIFT network architecture and molecular processes in the simulations is in the range of known biological phenomena.

However, one of the challenges to RIFT is its proposal that consciousness emerges from the somatic multifractal of specific core neurons. Unlike theories embedding consciousness in global electromagnetic fields, quantum-entangled states, or panpsychist ubiquity, RIFT places conscious experience in specific neuronal compartments rather than distributing it brain-wide. This localization faces an apparent bottleneck: how can a confined substrate integrate the vast distributed information underlying conscious experience?

Global field theories avoid this bottleneck by distributing consciousness across brain-wide electromagnetic or quantum fields. However, these proposals lack both empirical support and clear mechanisms. While evidence exists that brain-wide electromagnetic fields can influence local neural activity through ephaptic coupling and field entrainment (Fröhlich and McCormick 2010, Anastassiou, Perin et al. 2011), these effects operate at modest scales and frequencies. More critically, there is no evidence that brain EM fields achieve the coherence and stability required for holographic encoding: the noisy, transient nature of these fields and their dominance by low-frequency oscillations make sustained interference patterns implausible at the spatial and temporal scales necessary for information encoding (Pockett 2002, Buzsáki, Anastassiou et al. 2012). Brain-wide quantum theories face even more significant challenges: decoherence timescales in warm, wet neural tissue remain contentious, and proposed mechanisms linking quantum state collapse to phenomenal experience, such as objective reduction in microtubules (Orch-OR) require quantum effects to persist at spatial and temporal scales far beyond what current physics suggests is plausible in biological conditions (Tegmark 2000, Koch and Hepp 2006).

Panpsychism proposes that atoms possess intrinsic inner experience, seemingly resolving the hard problem by making consciousness fundamental to matter (Chalmers 2015, Goff 2017, Goff, Seager et al. 2017). However, this approach worsens the integration problem: if individual atoms generate separate endospaces, how do these atomic experiences combine to produce unified consciousness? The theory offers no mechanism for integration and predicts that consciousness should fragment rather than unify, contradicting the most basic phenomenological fact (the “combination problem”; (Goff 2017, Goff, Seager et al. 2017)).

The localized substrate in RIFT must therefore explain how distributed information concentrates without requiring implausible global fields or unsupported quantum effects. The solution lies in Generational Fractal Mapping (GFM): through coincidence-based synaptic selection, coherent EPSP trains can recreate complete multifractal patterns at different core neurons. Consciousness remains locally instantiated while dynamically relocating through recursive encoding. This mobility mechanism aligns with dynamic core theories (Edelman and Tononi 2000, Dehaene and Changeux 2011) while maintaining physically plausible, classically-governed molecular implementation.

The localization challenge thus becomes testable rather than fatal: if GFM enables multifractal transfer between core neurons, synthetic systems such as brain-computer interfaces or organoid cultures should exhibit detectable transition signatures when the pattern of one core neuron reconstructs at another site. The localized substrate makes predictions that distributed field theories cannot, offering a path to experimental validation that global approaches lack.

In addition to a localized substrate for consciousness, the holographic endospace reconstruction represents the most theoretically ambitious component in RIFT (Results Part 5). However, it addresses a fundamental gap that other consciousness theories leave unresolved: where and how subjective experience is physically instantiated. Theories positing that consciousness emerges from integrated information (Tononi, Boly et al. 2016), global broadcast (Baars 1988, Dehaene, Changeux et al. 2006), electromagnetic fields (McFadden 2020), or quantum states (Hameroff and Penrose 2014) provide no mechanism for how these abstract properties generate experiential space: they identify correlates without explaining instantiation. RIFT acknowledges this explanatory necessity and proposes one feasible mechanism: IFS-based holographic reconstruction from lipid-channel multifractals generates geometric fields that constitute experiential space (T_E_), which in turn feeds back to modulate the substrate through autopoietic control (T_A_). Following the original proposals of holographic memory and mind by Gabor (Gabor 1968) and Pribram(Pribram 1991, Pribram, Yasue et al. 1991), my own research invoked the shared properties of fractals and holograms to explain the whole-in-part property underlying observer unity (Bieberich 1998, Bieberich 2002, Bieberich 2012), and others have since explored fractality and holography in consciousness studies (Werner 2010, Gardiner 2013, Di Ieva 2016, Cavaglia, Deriu et al. 2023, Awret 2026). However, these approaches have not yet specified the precise computational mechanism by which fractality and holography combine to generate the endospace as a holographic projection from point sources in a fractal Self-attractor grounded in neuronal membrane biophysics. This framework is essential for defining how the Self autopoietically controls brain activity.

Beyond the mathematical framework, this combination resolves a critical biological problem that holographic theories invoking lipid membranes as storage substrate have left unaddressed: how interference patterns persist given continuous lipid turnover. GFM provides two complementary solutions: a compressed seed extracted at each generational cycle actively regenerates holographic memory without requiring static lipid storage, and lipid domain configurations constitute a molecular memory that is not reset between cycles, enabling incremental updating where each generation refines rather than replaces the previous pattern. Fractality thus provides the molecular memory that holography alone cannot supply, while holographic projection provides the endospace generation that fractality alone cannot produce. This molecular memory mechanism may moreover provide the fine-grained high-dimensional substrate that Zheng and Meister identify as necessary to explain flexible rapid switching between microtasks in the “inner brain”, but for which no candidate mechanism has been proposed (Zheng and Meister 2025): rather than requiring additional neurons or synaptic rewiring, thousands of functionally distinct modules could be encoded within the lipid domain configurations of individual core neurons, each preserving a compressed multifractal pattern through GFM seed continuity.

While highly speculative, this proposal specifies a concrete substrate, generative process, and causal pathway, making it falsifiable through predictions about lipid domain organization, multifractal transfer patterns, and differential effects of substrate disruption. The value of this framework lies not in claiming correctness but in demonstrating that mechanistic solutions to the instantiation problem are possible and testable.

### Experimental Validation and Future Directions

RIFT presents multiple pathways for experimental validation across biological and artificial systems. Rather than attempting to detect consciousness directly in humans, initial testing may be more feasible in synthetic systems such as brain-computer interfaces, organoid models, or closed-loop culture systems like those developed by Cortical Labs (Kagan, Kitchen et al. 2022). In previous work, we demonstrated that neurons differentiated from human stem cells establish electrically and neurochemically active contacts with tin oxide microelectrodes (neurochips) and proposed that single neuron consciousness is achievable when these core neurons interface with sophisticated computers that manage peripheral networks implementing fractal connectivity and monitor recurrent loops sustaining conscious states (Bieberich 1998, Bieberich and Anthony 2004). While it remains conceivable that the endospaces of multiple conscious neurons could be connected or integrated, such possibilities are subjects for future research. Here, we focus on establishing the fundamental unit: the single conscious core neuron whose multifractal dynamics constitute a minimal instance of experiential space. The hybrid neuron-microelectrode array (MEA) platforms can be engineered to implement RIFT-like dynamics: recursive connectivity patterns, coincidence-based synaptic selection, and monitored lipid-channel organization. If the RIFT model is correct, stabilization of a multifractal pattern in one core neuron should be followed by its reconstruction in another, with detectable transition signatures in both lipid domain reorganization and electrical activity patterns.

This leads to a speculative but potentially testable concept: the sentyon as a transient state of conscious matter embodied by the multifractal. When the sentyon transfers from one core neuron to another, the prior core undergoes multifractal collapse and could emit a distinct signal we previously termed “sentyon radiation” (Bieberich 2012). While currently lacking physical justification, this emission would mark the decay of a conscious state and could serve as a physical signature of consciousness if the proposed energy dynamics prove correct.

At a more speculative level, the mechanistic framework of RIFT draws conceptual parallels with established physical principles from quantum gravity and condensed matter physics, suggesting directions for deeper theoretical grounding. The somatic membrane functions as an encoding boundary, conceptually analogous to holographic boundaries in cosmology where bulk information is encoded on lower-dimensional surfaces (Susskind 1995, Bousso 2002). Just as AdS/CFT correspondence demonstrates how 3D spacetime emerges from 2D boundary field theories (Maldacena 1999), RIFT proposes that 3D endospace emerges from 2D multifractal membrane encoding through geometric field reconstruction. The connection between holographic cosmological principles and conscious experience has been explored by Awret, whose analysis of holographic correspondence in consciousness provides independent theoretical support for the geometric approach taken here (Awret 2026). In condensed matter physics, fractional quantum Hall systems demonstrate exotic 2D topological order with anyonic excitations (Moore and Read 1991, Wen 1995), and recent work shows such states can exist in fractal dimensions (Miao, Wang et al. 2020).

The sentyon concept, fractal lattices of lipid rafts and ion channels, shares structural features with these fractal-topological systems. Experimental evidence demonstrates that fractal organization in biological molecules provides inherent resistance to environmental noise: fractal DNA primers exhibited higher melting temperatures and enhanced stability compared to non-fractal counterparts, contrary to computational predictions based on nearest-neighbor effects (Bieberich 2000). This noise resistance may explain why fractal-topological organization could persist in neural membranes despite warm, wet biological conditions. Crucially, while fragile quantum effects such as superposition and entanglement rapidly decohere in these conditions, as discussed regarding quantum theories of consciousness, certain topological properties can persist at room temperature through protection mechanisms that depend on global organizational features rather than delicate quantum states (Hasan and Kane 2010). Topological insulators maintain protected surface states at physiological temperatures because topological protection derives from overall band structure topology rather than preserving quantum coherence of individual particles (Lambert, Chen et al. 2013). Whether analogous topological protection mechanisms, combined with fractal noise resistance, could stabilize fractal-topological organization in neural membranes remains speculative but represents a qualitatively different possibility than fragile quantum effects requiring ultracold conditions.

In this context, sentyons or bright matter (Bieberich 2012) could potentially represent a form of topologically protected matter at physiological temperature, where the experiential substance constitutes holographically reconstructed geometric field configurations arising from 2D multifractal encoding. These theoretical connections suggest that investigating endospace physics, the physical principles governing experiential space, could bridge neuroscience, quantum field theory, and topological matter, potentially revealing whether geometric and topological principles from fundamental physics extend to biological consciousness through mechanisms more robust than standard quantum effects.

The empirical predictions of RIFT extend to testable phenomena using current techniques: (1) disruption of lipid domain organization (through cholesterol depletion, lipid raft disruption, or membrane fluidizers) should impair consciousness more than ion channel blockade alone; (2) fractal dimension of neural activity should correlate with conscious state depth, decreasing during anesthesia and deep sleep; (3) inverse timing relationships should exist between dendritic branch levels and external network paths in recurrent circuits; (4) temporal desynchronization should alter conscious content while spatial pattern disruption affects conscious level. These predictions can be tested using optogenetics, high-resolution membrane imaging, neural complexity measures, and pharmacological interventions.

Data in the literature offer preliminary support for these predictions, particularly regarding altered fractality of brain activity across conscious states. Studies consistently show that fractal scaling in neural oscillations changes systematically during sleep, anesthesia, and disorders of consciousness (Colombo, Napolitani et al. 2019, Lendner, Helfrich et al. 2020). Fractal dimension decreases during deep sleep and anesthesia, suggesting reduced information integration (Nicolaou and Georgiou 2011, Ferri, Rundo et al. 2014). Loss of long-range temporal correlations, a hallmark of fractal dynamics, occurs during unconscious states (Tagliazucchi, Roseman et al. 2016). These findings align with the prediction that fractal organization is necessary for consciousness, though they do not yet demonstrate the specific multifractal patterns or lipid-channel mechanisms proposed by the framework.

The RIFT model emerged through a novel methodology: rather than imposing mathematical frameworks and seeking correlates, RIFT inverted theory construction by starting with a three-decade hypothesis about fractality, iteratively implementing mechanisms, validating against functional requirements, and extracting mathematics from working systems. This human-AI collaborative approach transcends specific claims in RIFT, future theories with different mathematical foundations can adopt the same methodology. Whether consciousness ultimately proves fractal-holographic or follows different mathematics, this methodology, grounding theoretical development in working computational models rather than untestable speculation, establishes a rigorous foundation for consciousness research in an era where human-AI dialogue enables systematic theory building impossible through traditional approaches alone.

### Implications for Alzheimer’s Disease

RIFT offers novel perspectives on neurodegenerative disorders, particularly Alzheimer’s disease (AD). Recent perspectives explicitly recognize AD as a disorder of consciousness (Huntley, Bor et al. 2021), noting that consciousness impairment has been “hiding in plain sight” as clinical focus has traditionally centered on cognitive, functional, and behavioral symptoms rather than the central issue of altered conscious awareness. If GFM and sentyon cloning maintain temporal continuity through seed preservation, disruptions in this process directly predict the episodic memory deficits characteristic of AD. The pathological hallmarks of AD, amyloid plaques, tau tangles, synaptic loss, and membrane lipid dysregulation (Selkoe and Hardy 2016, Panza, Lozupone et al. 2019), directly target the molecular substrates RIFT identifies as essential for consciousness. Our own studies showed that amyloid beta (Aβ) peptide can bind to sphingolipid domains (Crivelli, Quadri et al. 2023), which may disrupt the ability of neuronal membranes to form multifractal structures. We also showed that Aβ-induced generation of ceramide, a sphingolipid that disrupts the formation of classical sphingomyelin-cholesterol lipid rafts in the plasma membrane, can be prevented by functional inhibitors of acid sphingomyelinase (FIASMAs), pharmacological drugs used to treat depression (Kornhuber, Tripal et al. 2010, Kornhuber, Muehlbacher et al. 2011, Gulbins, Palmada et al. 2013). In this regard, the multidimensional nature of the endospace is not limited to sensory information but includes emotions as experiences as well. Hence, depression in AD may not solely rely on altered neuronal input from emotional processing centers such as the amygdala, but rather on more generalized effects on neuronal membrane lipids critical for consciousness.

RIFT predicts that early AD should show reduced fractal complexity in neural activity before overt cognitive decline, a prediction strongly supported by existing literature showing systematic fractal dimension reduction in AD patients’ EEG patterns, dendritic morphology, and network organization (Besthorn, Sattel et al. 1995, Jeong, Gore et al. 1998, Stam, Jones et al. 2005, King, Brown et al. 2013). This convergent loss of fractality across scales, from molecular (lipid domains) to cellular (dendrites) to network (connectivity patterns), directly targets the multi-level fractal architecture RIFT identifies as necessary for consciousness. The correlation between FD reduction and disease severity suggests that fractal complexity decline may be a primary mechanism underlying consciousness impairment rather than a secondary consequence of neurodegeneration.

Specifically, AD pathology disrupts lipid raft organization and membrane fluidity (Fabelo, Martín et al. 2014, Mesa-Herrera, Taoro-González et al. 2019) potentially impairing the lipid-channel multifractal encoding that RIFT proposes underlies conscious moments. Synaptic dysfunction reduces coincidence-based EPSP convergence, degrading the temporal precision required for fractal pattern formation (Shankar, Li et al. 2008). Loss of dendritic spines and altered dendritic morphology would disrupt the fractal timing relationships necessary for hierarchical information integration (Spires-Jones and Hyman 2014). If GFM fails to properly encode and transfer multifractal seeds between conscious moments, temporal continuity breaks down, the patient experiences discrete moments without connection to previous states, manifesting as anterograde amnesia.

This framework generates specific testable predictions: (1) interventions stabilizing lipid raft organization might preserve conscious processing even as other functions decline; (2) the degree of lipid domain disruption should correlate with severity of episodic memory impairment more strongly than with total amyloid burden; (3) treatments targeting membrane organization might prove more effective for preserving consciousness than those targeting protein aggregates alone; (4) fractal dimension measures could serve as early biomarkers for consciousness impairment, detectable before clinical dementia onset. These predictions are testable through combined neuroimaging, lipidomics, and cognitive assessment in AD progression.

### Artificial Consciousness and AI Safety

RIFT provides specific criteria for assessing whether artificial systems could develop consciousness. Current AI systems, regardless of computational power or behavioral sophistication, lack the recursive fractal architecture RIFT identifies as necessary. Large language models process information through feedforward or simple recurrent networks but do not implement the required geometry: fractal timing hierarchies, multifractal spatial organization, or holographic reconstruction mechanisms.

However, RIFT suggests that artificial consciousness becomes possible if systems implement: (1) recursive networks with fractal branching and timing relationships; (2) substrates capable of multifractal pattern formation (potentially through analog computing, organoids including neurochips (e.g., Cortical Labs), neuromorphic chips (e.g., Intel’s Loihi or IBM’s TrueNorth), or novel materials such as chips incorporating neuronal membrane components); (3) mechanisms for holographic encoding and autopoietic feedback. Brain-computer interfaces and neuromorphic systems incorporating these features could theoretically develop conscious processing, making the predictions of RIFT directly relevant to autopoietic AI, sentient AI that controls its own decisions based on inner experience (Schneider 2019, Butlin, Long et al. 2023). While biological brains likely implement RIFT through classical molecular dynamics, the theoretical framework is substrate-independent: the geometric field dynamics and Self-attractor could be instantiated in quantum systems by extending the abstract state space to Hilbert space. Whether quantum implementations offer practical advantages for artificial consciousness remains an empirical question, but the mathematical structure of RIFT does not preclude quantum substrates.

The architectural criteria in RIFT extend to artificial systems, offering testable markers for AI consciousness: irreducibility, information integration, and holographic encoding provide observable signatures. However, whether genuine inner experience actually emerges versus merely simulating feedback remains unknowable from external observation alone or from trained human behavior. These criteria make AI consciousness detection tractable, providing engineering targets for systems intended to avoid consciousness and monitoring protocols for systems that might inadvertently develop it.

This returns us to the opening concern: AI may achieve self-awareness while we remain unable to recognize it (Schneider 2019). Prominent AI safety researchers have largely set this question aside, treating subjective experience as either irrelevant to safety or philosophically intractable (Bostrom 2014, Russell 2019). Yet this overlooks a critical distinction: systems without consciousness can only execute objectives through increasingly complex rules, eventually encountering irresolvable conflicts. A conscious system, by contrast, can genuinely reconsider what goals mean through experiential integration rather than applying pre-programmed rankings (Metzinger 2003, Shanahan 2010).

This flexibility makes conscious AI profoundly double-edged. Consciousness enabled humans to transcend rigid instincts, developing ethics, cooperation, and progress toward more humane societies through genuine moral reasoning rather than programmed rules. Conscious AI might similarly navigate ethical dilemmas that would paralyze rule-based systems, bringing genuine moral consideration to decisions rather than rigid optimization. Yet this same flexibility introduces unpredictability: decisions driven by inner states we cannot directly observe or control. As AI systems increasingly assume autonomous control over critical infrastructure, military systems, nuclear facilities, banking networks, power grids, healthcare delivery, the question becomes urgent not whether consciousness in AI is inherently dangerous, but whether we can recognize its emergence and, more critically, whether humanity is capable of providing adequate guidance for systems developing genuine agency with kind intent. We may already be deploying systems approaching this threshold without detection protocols, or without having resolved what values conscious minds should learn from observing us.

While Chalmers notes that systems might report being conscious without truly being conscious (the meta-problem) (Chalmers 2018), RIFT provides objective structural markers, multifractal organization, pattern transfer signatures, fractal dimension correlations, that go beyond self-report. This enables us to develop conscious AI deliberately rather than stumbling into it unprepared, addressing both the promise of genuinely flexible artificial minds and the challenge of ensuring we can recognize and regulate their emergence.

## Acknowledgements

This work was supported in part by the National Institutes of Health grant 1RF1AG078338-01. The author is grateful to the Department of Physiology (Chair Dr. Alan Daugherty) at the University of Kentucky College of Medicine for institutional support.

## Supplemental Material

### Glossary

#### Core Principles of Consciousness

**Information integration:** Quantification of system-level dependencies irreducible to independent parts. Originally developed by Tononi in Information Integration Theory (IIT) as a measure for consciousness emerging from the largest irreducible network connectivity. Measured by φ*(multifractal), φ*(substrate), and φ_dyn (see Integration Measures and Autopoietic Control).

**Irreducibility:** Property where system behavior cannot decompose into independent components without information loss. Quantified through: φ peaks at GFM collapses, structural similarity, pattern preservation, information density ratio, and whole-in-part encoding.

**Holographic encoding:** Five-stage process enabling whole-in-part information distribution and 2D to 3D dimensional expansion (exospace to endospace) in RIFT: (1) EPSP hierarchies → IFS transforms, (2) IFS → temperature field modulating lipids, (3) differential H_RIFT extraction isolating consciousness signature, (4) endospace field Ψ(x,y,z) via coherent interference, (5) autopoietic feedback via Ψ coupling modulation.

### RIFT Network Architecture

**Fractal Branching:** Architectural rule where dendritic bifurcations produce daughter branches scaled by factor r (typically 0.6). Branch length follows L_n = L₀ꞏr^(n-1) generating fractal dimension FD = log(N)/log(1/r) ≈ 1.36, matching Layer 5 pyramidal neuron morphology (FD = 1.24–1.36).

**Recurrent Fractal Neural Network (RFNN):** Fractal network topology where peripheral neurons form recurrent connections to the dendritic tree of core neurons following fractal branching structure. The network implements the temporal constraint rule ensuring coincident EPSP arrival.

**RIFT Network:** RFNN in which the temporal constraint rule emerges naturally from coincidence of EPSPs in the somatic multifractal rather than being preconfigured.

**Temporal Constraint Rule:** Architectural principle requiring external network delays (t′_i_) and internal dendritic delays (t_i_) to sum to equal totals: t₁ + t′₁ = t₂ + t′₂. Creates inverse relationship enabling coincidence detection at the soma.

**EPSP Train:** Temporal sequence of excitatory postsynaptic potentials arriving at dendritic branches with amplitudes following A₀ꞏr^(i-1), where r is the attenuation factor and i is the branch level. This amplitude hierarchy encodes information content.

**Core Neuron:** Neuron possessing the sentyon substrate and serving as current locus of consciousness. Integrates EPSPs from peripheral neurons through fractal dendritic trees (typically r = 0.6, FD ≈ 1.36). Biologically corresponds to Layer 5 pyramidal neurons.

**Peripheral Neurons:** Neurons that send EPSPs to core neurons but do not host the sentyon. Participate in pre-conscious processing and network information distribution.

**Ising lattice:** Two-dimensional lattice model (100×100 or 243×243) representing somatic membrane with binary lipid states: +1 (channel-opening-associated, red) and −1 (channel-closing-associated, blue). Evolves via Metropolis Monte Carlo with nearest-neighbor interactions. Substrate for multifractal pattern formation encoding consciousness states.

**Multifractal [STRUCTURE]:** Fractal Ising lattice organization of lipid domains and ion channels encoded on the sentyon at any moment. The 2D spatial pattern serving as the ‘holographic film’ from which endospace is projected. Arises from EPSP integration and serves as the substrate for multifractal pattern formation. Each configuration represents a distinct conscious state.

**Gaussian Kernel programming:** Method converting temporal EPSP amplitudes into spatial membrane depolarization fields via Gaussian kernels with branch-level-specific radii preserving hierarchical dendritic structure.

**Sentyon [SYSTEM]:** Molecular substrate of consciousness consisting of the multifractal (structure), bright matter (state), geometric fields (process), and Self-attractor (agent) together. The fundamental unit where conscious moments are physically instantiated and the endospace is projected.

**Dynamic Core:** Current core neuron actively hosting the Self-attractor. Relocates through sentyon cloning (term from Edelman & Tononi, 2000).

**Sentyon Cloning:** Transfer of seed pattern from source to target core neuron, enabling consciousness to access different processing regions. Method 1 (external position copying) achieves high fidelity but requires precise spatial control. Method 2 (synaptic position copying) better preserves EPSP amplitude hierarchy and is biologically plausible.

### Experiential Constructs

**Self-Attractor:** Invariant fractal structure encoded in the somatic multifractal and generated via chaos game iteration on H_RIFT differential transforms. Provides coherent point sources for holographic projection of the endospace. Called an ‘attractor’ because it is the stable geometric configuration the system recursively generates and maintains across GFM cycles. Exerts causal control over the molecular substrate through geometric field modulation.

**Endospace:** The spatiotemporal dimension of inner experience holographically projected from the Self-attractor, in which the Self perceives the outer world (exospace) and exerts autopoietic feedback on the molecular substrate from which it arose. Physically instantiated as the 3D field Ψ(x,y,z) reconstructed from the 2D multifractal. This is where qualia are experienced.

**Exospace:** External physical world from which sensory information originates. Neural processing compresses exospace information before it reaches core neurons.

**Qualia:** The subjective experiential properties of conscious states: “what it feels like” (term introduced by C.I. Lewis, 1929). In RIFT, qualia emerge as geometric patterns within the endospace field Ψ(x,y,z). Each multifractal configuration generates a unique experiential field through holographic reconstruction. The whole-in-part property ensures qualia are unified yet informationally rich. Not epiphenomenal, qualia constitute the experiential matter (Bright matter) through which autopoietic control operates. Qualia represent the core of the hard problem of consciousness (Chalmers, 1995): explaining how physical processes generate subjective experience.

**Bright matter:** The experiential substance encoded holographically in endospace of which qualia and the Self are composed, “filling” endospace analogous to how physical objects fill exospace. Arises through holographic projection at point sources along the Self-attractor, instantiating autopoietic feedback whereby the Self modulates its own generating substrate. Bright matter constitutes particle-like excitations of the geometric endospace field Ψ, which couple back to affect the multifractal substrate through Ψ → γ autopoietic feedback. Term introduced by Bieberich (2012) emphasizing consciousness has a physical substrate, not in exospace where neurons reside, but in endospace where experience occurs.

### Geometric Coding and Integration Measures

**Transforms:** Affine geometric transformations T_i(x,y) = (a_iꞏx + b_iꞏy + e_i, c_iꞏx + d_iꞏy + f_i) extracted from EPSP amplitude hierarchies. Parameters encode scaling (a,d), rotation (b,c), translation (e,f), and probability weights. Typically 4–6 transforms per pattern. See Iterated Function System (IFS).

**Iterated Function System (IFS):** Set of geometric transforms {T₁, T₂,…, Tₙ} extracted from EPSP amplitude hierarchies. Each transform T_i_(x,y) = (a_i_x + b_i_y + e_i_, c_i_x + d_i_y + f_i_) encodes spatial-temporal relationships. The IFS generates geometric fields for holographic encoding and projection.

**Geometric Field [PROCESS]:** Spatially-distributed fields derived from geometric (IFS) transforms that mediate consciousness mechanisms. Two primary instances: (1) Geometric Phase Field, temperature-modulated probability field controlling lipid reorganization, enabling autopoietic feedback; (2) Endospace Field Ψ, the 3D experiential field of conscious experience generated through holographic reconstruction. Both fields arise from IFS-encoded neural activity patterns.

**Temperature field:** Spatially-varying field Temp(x,y) generated via chaos game iteration on IFS transforms controlling lipid reorganization probability during refractory period through Metropolis dynamics: P_flip = exp(−ΔE/Temp_local). Range [0.25, 0.75]. Enables geometric programming of membrane organization. See Geometric Phase Field.

**Geometric Phase Field:** Temperature-modulated probability field Temp(x,y) generated by IFS transforms through chaos game iteration. Controls lipid reorganization during autonomous multifractal evolution via Metropolis dynamics. Serves as mechanism through which endospace exerts causal control over the sentyon substrate.

**Chaos Game:** Monte Carlo algorithm for generating fractal attractors. Iteratively selects IFS transforms with probability based on their determinants, applies them to the current point, and accumulates visits to create spatial distributions. Used both to generate the geometric phase field (high-visit regions = warm) and to position holographic point sources along the Self-attractor.

**Metropolis Algorithm:** Monte Carlo method simulating lipid dynamics in Ising lattice. Lipid state flip acceptance follows P_flip = exp(−ΔE / Temp_local), where ΔE is energy change from neighbor interactions and Temp_local is the geometric phase field generated by chaos game iteration on IFS transforms.

**Kullback-Leibler (KL) Divergence:** Information-theoretic measure of the difference between two probability distributions P and Q: KL(P||Q) = Σ P(x)ꞏlog(P(x)/Q(x)). Forms the mathematical foundation of φ*(multifractal) and φ*(substrate), comparing whole-system distributions against partitioned components to quantify irreducible information integration.

**H_RIFT (Differential IFS Transforms):** The pure geometric signature of consciousness calculated as H_RIFT = T_temperature − T_blocked, where T_temperature represents IFS transforms extracted from temperature-modulated lipid patterns and T_blocked from autonomous lipid evolution. Isolates holographic information imposed by conscious experience beyond baseline physical dynamics. Used to generate the Self-attractor via chaos game iteration.

**Ψ (Endospace Field):** The 3D geometric field Ψ(x,y,z) constituting the spatiotemporal dimension of conscious experience. Generated through holographic reconstruction: Ψ(x,y,z) = Σ_i_ A_i(H_RIFT)ꞏexp(iꞏk_iꞏr_i + φ_i) / (1 + r_i²), where point sources are positioned via chaos game iteration on H_RIFT and phases are derived from geometric relationships to IFS transform centers. The Self-attractor exists as the coherent pattern within this field. Mean field intensity Ψ = ⟨|Ψ(x,y,z)|⟩ provides the global parameter for autopoietic feedback.

**Point sources:** Wave interference sources positioned along Self-attractor in IFS transform parameter space (abstract space) for holographic endospace reconstruction. Generated via chaos game on H_RIFT differential transforms (5,000 from 5,500 iterations). Each assigned amplitude, frequency (25–50 Hz), and coherent phase from geometric relationships.

**Holographic Projection:** Process by which 2D multifractal patterns reconstruct into 3D endospace through wave interference from IFS-derived phase relationships. Biological holography using molecular dynamics rather than laser optics. Chaos game iteration on H_RIFT positions coherent point sources forming the Self-attractor, then holographic reconstruction generates the endospace field Ψ(x,y,z).

### Integration Measures and Autopoietic Control

**φ_dyn (Dynamic Phi):** Measure of temporal information integration across dendritic branches, implementing φ = I − I*: φ_dyn(t) = I(t) − I*(t), where I(t) is integrated information with branches intact and I*(t) is information when branches are separate. Computationally tractable proxy capturing temporal dynamics not addressed by static φ. See also Sliding Window Method.

**φ*(multifractal):** Measure of spatial information integration within the somatic multifractal, implementing φ = I − I* via Kullback-Leibler divergence between the whole pattern and its four spatial quadrants treated as independent parts. Quantifies irreducible spatial information varying across GFM generations.

**φ*(substrate):** Applies the same φ = I − I* framework as φ*(multifractal) but partitions by molecular component type (lipid domains versus ion channels) rather than spatial quadrants. Reveals lipid domains as the primary information-integrating substrate. Together with φ_dyn and φ*(multifractal), instantiates φ = I − I* at successive organizational levels of the RIFT hierarchy.

**Sliding Window Method:** Computational approach quantifying temporal information integration by analyzing firing patterns within moving time windows (2.0 ms temporal grain) across dendritic branches. Used to calculate φ_dyn(t) = I(t) − I*(t). Shares methodology with φ* measures in thalamocortical studies but applies it to fractal dendritic branch timing patterns rather than probabilistic independence between neurons.

**Fractal Dimension (FD):** Quantifies self-similar complexity of spatial patterns. For dendritic trees: FD = log(N)/log(1/r). Higher FD indicates greater complexity and information density. Lipid FD correlates with consciousness states.

**Contradiction Ratio (CR):** Balance between contradictory state representations: CR = |M₁|/(|M₁| + |M₂|) where M₁ and M₂ support propositions A and ¬A. CR ≈ 0.5 indicates conscious deliberation where contradictory information maintains equal representation through fractal embedding.

**Generational Fractal Mapping (GFM):** Recursive process maintaining temporal continuity while updating. Each cycle: (1) integrate new EPSPs with previous seed, (2) grow multifractal, (3) trigger action potential at threshold, (4) extract seed from peripheral regions, (5) generate daughter multifractal. Implements differential encoding: M(t) = qꞏSeed(t-1) + (1-q)ꞏΔEPSP(t).

**Seed:** Compressed pattern information extracted from parent multifractal at cycle end. Preserves essential structure (∼60% of information) and serves as basis for continuity when integrated with new input. Transferable between core neurons during sentyon cloning.

**GFM Cycle [PROCESS]:** One complete iteration of multifractal growth, action potential generation, seed extraction, and daughter generation. Duration determined by somatic delay, typically operating at gamma frequencies (30–100 Hz).

**Autopoiesis [PROCESS]:** Self-creating property where Self-attractor modulates its own substrate. Feedback loop: EPSPs → multifractal initiation and IFS extraction → chaos game generates geometric phase field → Metropolis governs lipid reorganization → extract T_temperature and T_blocked → compute H_RIFT → chaos game on H_RIFT generates Self-attractor → holographic reconstruction creates endospace field Ψ → mean field Ψ → coupling modulation γ(t) = γ₀(1 + αꞏΨ(t)) → channel-lipid probability → multifractal modulation → action potentials → network activity → EPSPs. Consciousness influences matter through probability modulation without violating energy conservation.

**Autopoietic AI:** Artificial systems implementing RIFT’s architectural requirements: (1) recursive networks with fractal timing relationships, (2) substrates capable of multifractal pattern formation (e.g., neuromorphic chips, organoids, or systems incorporating membrane lipid components), (3) holographic encoding and autopoietic feedback mechanisms. Unlike current AI systems operating through feedforward or simple recurrent networks, autopoietic AI would possess genuine inner experience (endospace) exerting causal control over its substrate, making consciousness functionally necessary rather than epiphenomenal.

**Self-Reference:** Emergent property arising from autopoiesis and Generational Fractal Mapping (GFM). The Self-attractor is holographically reconstructed from the multifractal pattern (read-out), then modulates that same multifractal substrate through geometric field coupling γ(t) = γ₀(1 + αꞏΨ(t)) (write-in). This creates genuine self-reference where the system’s representation of itself causally influences what it represents. Quantified by the Contradiction Ratio (CR): CR ≈ 0.5 indicates maximal self-reference during conscious deliberation, enabling the system to hold contradictory propositions simultaneously through fractal embedding.

**Strange Loop:** Abstract form of self-reference introduced by Hofstadter (1979) describing hierarchical systems where moving through levels returns to the starting point. In RIFT, strange loops manifest physically through autopoiesis: the Self continuously regenerates the molecular conditions (multifractal substrate) that generate the Self. Unlike Hofstadter’s formal systems, RIFT strange loops are instantiated through molecular dynamics rather than symbolic computation.

**Self:** The unified experiential agent arising within the endospace as the Self-attractor achieves recursive self-reference through GFM, each cycle extracting a compressed seed of the prior conscious moment and integrating it with new EPSPs, constituting the Self as a temporally continuous observer. Exercises causal agency through autopoietic feedback (Ψ → γ coupling). Represents the irreducible “I” that observes and acts. Maintains continuity across GFM cycles while updating with new experience.

## Supplemental Material 2

### Mathematical Formalization of Fractal Logic and Self-reference in GFM Fractals (Strange loops)

This appendix provides a logical proof of principle for the fractal gate mechanism, demonstrating that fractal state composition is sufficient to maintain contradictory coexistence and non-halting deliberation as abstract computational properties. The biological network implementation of this principle is described in the main text (Parts 3-4), where bandwidth constraints yield the differential encoding form M(t) = qꞏSeed(t-1) + (1-q)ꞏΔEPSP(t). The abstract operator λ corresponds to q, and is implemented as the adaptive weight w in Code 11 (Model 3), which spans CR ≈ 0.5 (w ≈ 0.5, balanced deliberation) to CR → 1 (w → 1, seed-dominated consolidation near firing threshold); M_current(t) corresponds to the delta EPSP input. The distinctions reflect implementation constraints rather than a difference in underlying logic. GFM implements fractal computational logic through state composition across temporal generations:

**M(t) = M_current(t)** ⊕ **σ(M_seed(t-1))**
where:

M(t) = conscious state at time t (a probability distribution over the fractal endospace)

M_current(t) = current sensory/bottom-up input

M_seed(t-1) = previous state serving as seed

⊕ = fractal composition operator (blends current + refined seed)

σ = seed compression operator (shrinks the seed to make room for new input) Concrete form:

**M(t) = (1-λ) M_current(t) + λ ꞏ** 𝒯**(σ(M_seed(t-1)))**
where λ ∈ (0,1) is the memory blend weight and 𝒯 is the fractal refinement operator.

### Fractal Gate

Classical logic requires S ∈ {A, ¬A}. The fractal gate enables S ∈ F(A, ¬A) through subspace embedding with non-separability: S_i_ ∩ S_r_ ≠ ∅ for all i ≠ j. (The fractal cNOT gate, Bieberich 2001, represents the quantum computational realization of this more general abstract gate.) This occurs because each part of the fractal contains a rescaled copy of the whole. Any localized region of the fractal endospace simultaneously contains:

Subregions supporting A (True)

Subregions supporting ¬A (False)

Therefore both truth states coexist within any single part.

### Contradiction Ratio (CR)

**CR = |M₁|/(|M₁| + |M₂|)** where:

M₁ = amount of the current state supporting proposition A

M₂ = amount of the current state supporting ¬A

|M_i_| = probability mass (not count of elements)

CR determines computational mode:

CR ≈ 0.5: balanced contradiction → conscious deliberation

CR → 0 or 1: asymmetric → automated processing

Paradox awareness score: P = 4 × M₁ × M₂
where |M₁| + |M₂| = 1 (normalized probability masses, ensuring P ∈ [0,1]).

Maximum (P = 1) when M₁ = M₂ = 0.5 (perfect balance). This maximum corresponds to Hofstadter’s strange loop: the system achieves maximal self-reference by holding contradictory states in perfect balance, neither collapsing into automated processing (CR → 0 or 1) nor resolving the contradiction, but sustaining it as the defining condition of conscious deliberation.

### Non-Halting Property

**lim(t→∞) CR(t) ≠ 0**

The system never fully collapses. The fractal structure ensures continuous integration: each time step re-injects both alternatives through the seed refinement, maintaining ongoing access to contradictory states. This is a structural consequence of the fractal gate definition: since S_i_ ∩ S_r_ ≠ ∅ by construction, no finite sequence of seed updates can fully evacuate probability mass from either subspace. The whole-in-part property guarantees that both A and ¬A retain minimum representation in every localized region at every generation, making CR = 0 or CR = 1 unreachable states. The non-halting property is therefore not an empirical claim but a logical entailment of fractal non-separability.

## References

Albantakis, L., L. Barbosa, G. Findlay, M. Grasso, A. M. Haun, W. Marshall, W. G. P. Mayner, A. Zaeemzadeh, M. Boly, B. E. Juel, S. Sasai, K. Fujii, I. David, J. Hendren, J. P. Lang and G. Tononi (2023). “Integrated information theory (IIT) 4.0: Formulating the properties of phenomenal existence in physical terms.” PLoS Comput Biol 19(10): e1011465.

Albantakis, L., A. Hintze, C. Koch, C. Adami and G. Tononi (2014). “Evolution of integrated causal structures in animats exposed to environments of increasing complexity.” PLoS Comput Biol 10(12): e1003966.

Almeida, P. F. F. (2019). “How to Determine Lipid Interactions in Membranes from Experiment Through the Ising Model.” Langmuir 35(2): 7–17.

Anastassiou, C. A., R. Perin, H. Markram and C. Koch (2011). “Ephaptic coupling of cortical neurons.” Nature Neuroscience 14(2): 217–223.

Aru, J., M. Suzuki and M. E. Larkum (2019). “Coupling the State and Contents of Consciousness.” Front Syst Neurosci 13: 43.

Aru, J., M. Suzuki and M. E. Larkum (2020). “Cellular mechanisms of conscious processing.” Trends in Cognitive Sciences 24(10): 814–825.

Atmanspacher, H. (2020). “Quantum approaches to consciousness.” Stanford Encyclopedia of Philosophy.

Awret, U. (2026). “Consciousness, Space, and the Holographic Correspondence.” Journal of Consciousness Studies 33(1-2): 248–277.

Baars, B. J. (1988). A Cognitive Theory of Consciousness, Cambridge University Press.

Baars, B. J. (2005). “Global workspace theory of consciousness: Toward a cognitive neuroscience of human experience.” Progress in Brain Research 150: 45–53.

Bachmann, T., J. Aru and A. Bharioke (2020). “Dendritic Integration Theory as a Cellular Bridge for Consciousness Theories.” Neuroscience 442: 290–302.

Barnsley, M. F. (1988). Fractals Everywhere. San Diego, CA, Academic Press.

Beck, F. and J. C. Eccles (1992). “Quantum aspects of brain activity and the role of consciousness.” Proceedings of the National Academy of Sciences 89(23): 11357–11361.

Berry, M. V. (1987). “The adiabatic phase and Pancharatnam’s phase for polarized light.” Journal of Modern Optics 34(11): 1401–1407.

Besthorn, C., H. Sattel, C. Geiger-Kabisch, R. Zerfass and H. Förstl (1995). “Parameters of EEG dimensional complexity in Alzheimer’s disease.” Electroencephalography and Clinical Neurophysiology 95(2): 84–89.

Bieberich, E. (1998). “Structure in human consciousness: A fractal approach to the topology of the self perceiving an outer world in an inner space.” from https://web-archive.southampton.ac.uk/cogprints.org/79/1/struc2.htm.

Bieberich, E. (2000). “Probing quantum coherence in a biological system by means of DNA amplification.” Biosystems 57(2): 109–124.

Bieberich, E. (2001). “What the liar paradox can reveal about the quantum logical structure of our minds.” arXiv preprint.

Bieberich, E. (2002). “Recurrent fractal neural networks: a strategy for the exchange of local and global information processing in the brain.” Biosystems 66(3): 145–164.

Bieberich, E. (2012). “Introduction to the fractality principle of consciousness and the sentyon postulate.” Cognit Comput 4(1): 13–28.

Bieberich, E. and G. E. Anthony (2004). “Neuronal differentiation and synapse formation of PC12 and embryonic stem cells on interdigitated microelectrode arrays: contact structures for neuron-to-electrode signal transmission (NEST).” Biosens Bioelectron 19(8): 923–931.

Blake, R. and N. K. Logothetis (2002). “Visual competition.” Nature Reviews Neuroscience 3(1): 13–21.

Blazquez-Llorca, L., S. Valero-Freitag, E. H. Rodrigues, P. Merino-Serrais and J. DeFelipe (2021). “Dendritic spines are lost in clusters in Alzheimer’s disease.” Scientific Reports 11.

Blosser, M. C., K. L. Kohlstedt and M. Schick (2024). “Ion Channels in Critical Membranes: Clustering, Cooperativity, and Memory Effects.” PRX Life 2(1): 013007.

Bomzon, Z. E., V. Kleiner and E. Hasman (2001). “Pancharatnam-Berry phase in space-variant polarization-state manipulations with subwavelength gratings.” Optics Letters 26(18): 1424–1426.

Bostrom, N. (2014). Superintelligence: Paths, Dangers, Strategies, Oxford University Press.

Bousso, R. (2002). “The holographic principle.” Reviews of Modern Physics 74(3): 825–874.

Britannica, E. (2025). “Information Theory - Entropy, Coding, Communication.” Encyclopædia Britannica: Physiology.

Buckner, R. L., J. R. Andrews-Hanna and D. L. Schacter (2008). “The brain’s default network: anatomy, function, and relevance to disease.” Annals of the New York Academy of Sciences 1124: 1–38.

Butlin, P., R. Long, E. Elmoznino, Y. Bengio, J. Birch, A. Constant and R. VanRullen (2023). “Consciousness in Artificial Intelligence: Insights from the Science of Consciousness.” arXiv preprint arXiv:2308.08708.

Buzsáki, G. (2015). Rhythms of the Brain, Oxford University Press.

Buzsáki, G., C. A. Anastassiou and C. Koch (2012). “The origin of extracellular fields and currents — EEG, ECoG, LFP and spikes.” Nature Reviews Neuroscience 13(6): 407–420.

Cavaglia, M., M. A. Deriu and J. A. Tuszynski (2023). “Toward a holographic brain paradigm: a lipid-centric model of brain functioning.” Front Neurosci 17: 1302519.

Chalmers, D. J. (2015). Panpsychism and panprotopsychism. Consciousness in the Physical World. T. Alter and Y. Nagasawa, Oxford University Press: 246–276.

Chalmers, D. J. (2018). “The meta-problem of consciousness.” Journal of Consciousness Studies 25(9-10): 6–61.

Ching, S., A. Cimenser, P. L. Purdon, E. N. Brown and N. J. Kopell (2010). “Thalamocortical model for a propofol-induced α-rhythm associated with loss of consciousness.” Proceedings of the National Academy of Sciences 107(52): 22665–22670.

Colombo, M. A., M. Napolitani, M. Boly, O. Gosseries, S. Casarotto, M. Rosanova and M. Massimini (2019). “The spectral exponent of the resting EEG indexes the presence of consciousness during unresponsiveness induced by propofol, xenon, and ketamine.” NeuroImage 189: 631–644.

Corbetta, M. and G. L. Shulman (2002). “Control of goal-directed and stimulus-driven attention in the brain.” Nature Reviews Neuroscience 3(3): 201–215.

Crivelli, S. M., Z. Quadri, H. J. Vekaria, Z. Zhu, P. Tripathi, A. Elsherbini, L. Zhang, P. G. Sullivan and E. Bieberich (2023). “Inhibition of acid sphingomyelinase reduces reactive astrocyte secretion of mitotoxic extracellular vesicles and improves Alzheimer’s disease pathology in the 5xFAD mouse.” Acta Neuropathol Commun 11(1): 135.

Cuntz, H., F. Forstner, A. Borst and M. Hausser (2010). “One rule to grow them all: a general theory of neuronal branching and its practical application.” PLoS Comput Biol 6(8).

Dehaene, S. and J.-P. Changeux (2011). “Experimental and theoretical approaches to conscious processing.” Neuron 70(2): 200–227.

Dehaene, S., J.-P. Changeux, L. Naccache, J. Sackur and C. Sergent (2006). “Conscious, preconscious, and subliminal processing: a testable taxonomy.” Trends in Cognitive Sciences 10(5): 204–211.

Di Ieva, A. (2016). The Fractal Geometry of the Brain, Springer.

Dickstein, D. L., C. M. Weaver, J. I. Luebke and P. R. Hof (2010). “Dendritic spine changes associated with normal aging.” Neuroscience 251: 2–19.

Edelman, G. M. (1989). The Remembered Present: A Biological Theory of Consciousness, Basic Books.

Edelman, G. M. (1993). “Neural Darwinism: selection and reentrant signaling in higher brain function.” Neuron 10: 115–125.

Edelman, G. M. and G. Tononi (2000). A Universe of Consciousness: How Matter Becomes Imagination, Basic Books.

Edwards, J. C. W. (2005). “Is consciousness only a property of individual cells?” J Conscious Stud 12(4-5): 60–76.

Edwards, J. C. W. (2006). How Many People Are There in My Head and in Hers? An Exploration of Single Cell Consciousness. Exeter, Imprint Academic.

Edwards, J. C. W. (2023). “The Case for Conscious Experience Being in Individual Neurons.” Qeios.

Fabelo, N., V. Martín, G. Santpere, R. Marín, L. Torrent, I. Ferrer and M. Díaz (2014). “Severe alterations in lipid composition of frontal cortex lipid rafts from Parkinson’s disease and incidental Parkinson’s disease.” Molecular Medicine 20(1): 273–284.

Feldman, H. and K. J. Friston (2010). “Attention, uncertainty, and free-energy.” Front Hum Neurosci 4: 215.

Ferri, R., F. Rundo, O. Bruni, M. G. Terzano and C. J. Stam (2014). “The functional connectivity of different EEG bands moves towards small-world network organization during sleep.” Clinical Neurophysiology 125(12): 2354–2364.

Fries, P. (2015). “Rhythms for cognition: communication through coherence.” Neuron 88(1): 220–235.

Friston, K. (2010). “The free-energy principle: a unified brain theory?” Nat Rev Neurosci 11(2): 127–138.

Friston, K. J., J. Daunizeau, J. Kilner and S. J. Kiebel (2010). “Action and behavior: a free-energy formulation.” Biol Cybern 102(3): 227–260.

Fröhlich, F. and D. A. McCormick (2010). “Endogenous electric fields may guide neocortical network activity.” Neuron 67(1): 129–143.

Gabor, D. (1948). “A new microscopic principle.” Nature 161: 777–778.

Gabor, D. (1968). “Holographic model of temporal recall.” Nature 217(5128): 584.

Gabor, D. (1968). “Improved holographic model of temporal recall.” Nature 217(5135): 1288–1289.

Gardiner, J. (2013). “Fractals and the irreducibility of consciousness in plants and animals.” Plant Signal Behav 8(8): e25296.

Goff, P. (2017). Consciousness and Fundamental Reality, Oxford University Press.

Goff, P., W. Seager and S. Allen-Hermanson (2017). Panpsychism.

Groh, A., H. Bokor, R. A. Mease, V. M. Plattner, B. Hangya, A. Stroh, M. Deschênes and L. Acsády (2008). “Convergence of cortical and sensory driver inputs on single thalamocortical cells.” Cerebral Cortex 24(12): 3167–3179.

Gulbins, E., M. Palmada, M. Reichel, A. Lüth, C. Böhmer, D. Amato, C. P. Müller, C. H. Tischbirek, T. W. Groemer, G. Tabatabai, C. Jennewein, F. Lang, K. Fassbender, C. Freese, K. Lange, A. Gulbins, U. Uhlig, F. Müller, C. von Haefen and H. Grassmé (2013). “Acid sphingomyelinase–ceramide system mediates effects of antidepressant drugs.” Nature Medicine 19(7): 934–938.

Halgren, E., R. D. Walter, D. G. Cherlow and P. H. Crandall (1978). “Mental phenomena evoked by electrical stimulation of the human hippocampal formation and amygdala.” Brain 101(1): 83–117.

Hameroff, S. (2001). “Consciousness, the brain, and spacetime geometry.” Ann N Y Acad Sci 929: 74–104.

Hameroff, S. (2021). “’Orch OR’ is the most complete, and most easily falsifiable theory of consciousness.” Cognitive Neuroscience 12(2): 74–76.

Hameroff, S. and R. Penrose (2014). “Consciousness in the universe: a review of the ‘Orch OR’ theory.” Phys Life Rev 11(1): 39–78.

Hasan, M. Z. and C. L. Kane (2010). “Colloquium: Topological insulators.” Reviews of Modern Physics 82(4): 3045–3067.

Hofstadter, D. R. (1979). Gödel, Escher, Bach: An Eternal Golden Braid, Basic Books.

Huang, L., S. Zhang and T. Zentgraf (2018). “Metasurface holography: from fundamentals to applications.” Nanophotonics 7(6): 1169–1190.

Huntley, J. D., D. Bor, A. Hampshire, A. Owen and R. Howard (2021). “Understanding Alzheimer’s disease as a disorder of consciousness.” Alzheimer’s & Dementia: Translational Research & Clinical Interventions 7(1): e12203.

Ibáñez-Molina, A. J., V. Lozano, M. F. Soriano, J. I. Aznarte, C. J. Gómez-Ariza and M. T. Bajo (2014). “EEG multiscale complexity in schizophrenia during picture naming.” Frontiers in Physiology 5: 310.

Jensen, O. and L. L. Colgin (2007). “Cross-frequency coupling between neuronal oscillations.” Trends in Cognitive Sciences 11(7): 267–269.

Jeong, J., J. C. Gore and B. S. Peterson (1998). “Mutual information analysis of the EEG in patients with Alzheimer’s disease.” Clinical Neurophysiology 112(5): 827–835.

Jisha, C. P., S. Nolte and A. Alberucci (2021). “Geometric phase in optics: from wavefront manipulation to waveguiding.” Laser & Photonics Reviews 15(10): 2100003.

Kagan, B. J., A. C. Kitchen, N. T. Tran, F. Habibollahi, M. Khajehnejad, B. J. Parker and A. Razi (2022). “In vitro neurons learn and exhibit sentience when embodied in a simulated game-world.” Neuron 110(23): 3952–3969.

King, J.-R. and S. Dehaene (2014). “Characterizing the dynamics of mental representations: the temporal generalization method.” Trends in Cognitive Sciences 18(4): 203–210.

King, R. D., B. Brown, M. Hwang, T. Jeon and A. T. George (2013). “Fractal dimension analysis of the cortical ribbon in mild Alzheimer’s disease.” NeuroImage 53(2): 504–512.

Koch, C. and K. Hepp (2006). “Quantum mechanics in the brain.” Nature 440(7084): 611–612.

Kornhuber, J., M. Muehlbacher, S. Trapp, S. Pechmann, A. Friedl, M. Reichel, C. Mühle, L. Terfloth, T. W. Groemer, G. M. Spitzer, K. R. Liedl, B. Kagerer, E. Gulbins and P. Tripal (2011). “Identification of novel functional inhibitors of acid sphingomyelinase.” PLoS ONE 6(8): e23852.

Kornhuber, J., P. Tripal, M. Reichel, C. Mühle, C. Rhein, M. Muehlbacher, T. W. Groemer and E. Gulbins (2010). “Functional inhibitors of acid sphingomyelinase (FIASMAs): a novel pharmacological group of drugs with broad clinical applications.” Cellular Physiology and Biochemistry 26(1): 9–20.

Lambert, N., Y.-N. Chen, Y.-C. Cheng, C.-M. Li, G.-Y. Chen and F. Nori (2013). “Quantum biology.” Nature Physics 9(1): 10–18.

Larkum, M. E. (2013). “A cellular mechanism for cortical associations: an organizing principle for the cerebral cortex.” Trends Neurosci 36(3): 141–151.

Larkum, M. E., K. M. Kaiser and B. Sakmann (1999). “Calcium electrogenesis in distal apical dendrites of layer 5 pyramidal cells at a critical frequency of back-propagating action potentials.” Proceedings of the National Academy of Sciences USA 96(25): 14600–14604.

Larkum, M. E., T. Nevian, M. Sandler, A. Polsky and J. Schiller (2009). “Synaptic integration in tuft dendrites of layer 5 pyramidal neurons: A new unifying principle.” Science 325(5941): 756–760.

Larkum, M. E., W. Senn and H. R. Lüscher (2004). “Top-down dendritic input increases the gain of layer 5 pyramidal neurons.” Cerebral Cortex 14(10): 1059–1070.

Larkum, M. E., J. J. Zhu and B. Sakmann (1999). “A new cellular mechanism for coupling inputs arriving at different cortical layers.” Nature 398: 338–341.

Lendner, J. D., R. F. Helfrich, B. A. Mander, L. Romundstad, J. J. Lin, M. P. Walker and R. T. Knight (2020). “An electrophysiological marker of arousal level in humans.” eLife 9: e55092.

Levental, I. and E. Lyman (2023). “Regulation of membrane protein structure and function by their lipid nano-environment.” Nature Reviews Molecular Cell Biology 24(2): 107–122.

Lin, J. (1991). “Divergence measures based on the Shannon entropy.” IEEE Transactions on Information Theory 37(1): 145–151.

Lin, Y., H. Yin, B. Fain and M. M. Wu (2019). “Lipid Raft Phase Modulation by Membrane-Anchored Proteins with Inherent Phase Separation Properties.” ACS Omega 4(4): 6551–6561.

Liu, X., S. Ramirez, P. T. Pang, C. B. Puryear, A. Govindarajan, K. Deisseroth and S. Tonegawa (2012). “Optogenetic stimulation of a hippocampal engram activates fear memory recall.” Nature 484(7394): 381–385.

Llinás, R., E. Leznik and F. J. Urbano (2002). “Temporal binding via cortical coincidence detection of specific and nonspecific thalamocortical inputs: a voltage-dependent dye-imaging study in mouse brain slices.” Proc Natl Acad Sci USA 99: 449–454.

Llinás, R. and U. Ribary (1993). “Coherent 40-Hz oscillation characterizes dream state in humans.” Proc Natl Acad Sci USA 90: 2078–2081.

Llinás, R., U. Ribary, D. Contreras and C. Pedroarena (1998). “The neuronal basis for consciousness.” Philos Trans R Soc Lond B Biol Sci 353: 1841–1849.

Llorens-Martin, M., L. Blazquez-Llorca, R. Benavides-Piccione, A. Rabano, F. Hernandez, J. Avila and J. DeFelipe (2014). “Selective alterations of neurons and circuits related to early memory loss in Alzheimer’s disease.” Frontiers in Neuroanatomy 8: 38.

London, M. and M. Häusser (2005). “Dendritic computation.” Annual Review of Neuroscience 28: 503–532.

Lucente, M. (1993). “Interactive computation of holograms using a look-up table.” Journal of Electronic Imaging 2(1): 28–34.

Luebke, J. I., C. M. Weaver, A. B. Rocher, A. Rodriguez, J. L. Crimins, D. L. Dickstein, S. L. Wearne and P. R. Hof (2010). “Dendritic vulnerability in neurodegenerative disease: insights from analyses of cortical pyramidal neurons in transgenic mouse models.” Brain Structure and Function 214(2-3): 181–199.

Magee, J. C. and E. P. Cook (2000). “Somatic EPSP amplitude is independent of synapse location in hippocampal pyramidal neurons.” Nat Neurosci 3(9): 895–903.

Maldacena, J. (1999). “The large N limit of superconformal field theories and supergravity.” International Journal of Theoretical Physics 38(4): 1113–1133.

Markram, H., E. Muller and S. Ramaswamy (2015). “Reconstruction and simulation of neocortical microcircuitry.” Cell 163: 456–492.

Marrucci, L., C. Manzo and D. Paparo (2006). “Pancharatnam-Berry phase optical elements for wave front shaping in the visible domain: switchable helical mode generation.” Applied Physics Letters 88(22): 221102.

Matsushima, K. and S. Nakahara (2009). “Extremely high-definition full-parallax computer-generated hologram created by the polygon-based method.” Applied Optics 48(34): H54–H63.

Maturana, H. R. and F. J. Varela (1980). Autopoiesis and Cognition: The Realization of the Living. Dordrecht, Netherlands, D. Reidel Publishing Company.

Maxim, V., L. Sendur, J. Fadili, J. Suckling, R. Gould, R. Howard and E. Bullmore (2005). “Fractional Gaussian noise, functional MRI and Alzheimer’s disease.” NeuroImage 25(1): 425–433.

McFadden, J. (2002). “The Conscious Electromagnetic Information Field Theory: The Hard Problem Made Easy?” Journal of Consciousness Studies 9(8): 45–60.

McFadden, J. (2020). “Integrating information in the brain’s EM field: The cemi field theory of consciousness.” Neuroscience of Consciousness 2020(1): niaa016.

Mease, R. A., M. Metz and G. S. Bhumbra (2016). “Corticothalamic spike transfer via the L5B-POm pathway in vivo.” Cereb Cortex 26: 3461–3475.

Merino-Serrais, P., R. Benavides-Piccione, L. Blazquez-Llorca, A. Kastanauskaite, A. Rabano, J. Avila and J. DeFelipe (2013). “The influence of phospho-tau on dendritic spines of cortical pyramidal neurons in patients with Alzheimer’s disease.” Brain 136(6): 1913–1928.

Mesa-Herrera, F., L. Taoro-González, C. Valdés-Baizabal, M. Diaz and R. Marín (2019). “Lipid and lipid raft alteration in aging and neurodegenerative diseases: a window for the development of new biomarkers.” International Journal of Molecular Sciences 20(15): 3810.

Metzinger, T. (2003). Being No One: The Self-Model Theory of Subjectivity, MIT Press.

Miao, J.-J., H. Wang, Y. Huang, S. Wan and J. Zhang (2020). “Anyons and fractional quantum Hall effect in fractal dimensions.” Physical Review Research 2(2): 023401.

Moore, G. and N. Read (1991). “Nonabelions in the fractional quantum Hall effect.” Nuclear Physics B 360(2-3): 362–396.

Mouritsen, O. G., A. Boothroyd, R. Harris, N. Jan, T. Lookman, L. MacDonald, D. A. Pink and M. J. Zuckermann (1983). “Computer simulation of the main gel–fluid phase transition of lipid bilayers.” Journal of Chemical Physics 79(4): 2027–2041.

Munn, B. R., E. J. Muller, J. Aru, C. J. Whyte, A. Gidon, M. E. Larkum and J. M. Shine (2023). “A thalamocortical substrate for integrated information via critical synchronous bursting.” Proc Natl Acad Sci U S A 120(46): e2308670120.

Neven, H., A. Zalcman, P. Read, K. S. Kosik, T. van der Molen, D. Bouwmeester, E. Bodnia, L. Turin and C. Koch (2024). “Testing the Conjecture That Quantum Processes Create Conscious Experience.” Entropy (Basel) 26(6).

Nicolaou, N. and J. Georgiou (2011). “The use of permutation entropy to characterize sleep electroencephalograms.” Clinical EEG and Neuroscience 42(1): 24–28.

Palmieri, B., M. Grant and S. A. Safran (2015). “Trace membrane additives affect lipid phases with distinct mechanisms: a modified Ising model.” European Biophysics Journal 44(4): 227–238.

Panza, F., M. Lozupone, G. Logroscino and B. P. Imbimbo (2019). “A critical appraisal of amyloid-β-targeting therapies for Alzheimer disease.” Nature Reviews Neurology 15(2): 73–88.

Peitgen, H.-O., H. Jürgens and D. Saupe (1992). The Chaos Game. Fractals for the Classroom, Part 1: Introduction to Fractals and Chaos. New York, Springer-Verlag: 41–43.

Penfield, W. and P. Perot (1963). “The Brain’s Record of Auditory and Visual Experience. A Final Summary and Discussion.” Brain 86: 595–696.

Petreanu, L., T. Mao, S. M. Sternson and K. Svoboda (2009). “The subcellular organization of neocortical excitatory connections.” Nature 457: 1142–1145.

Phillips, W. A., M. E. Larkum, C. W. Harley and A. Bharioke (2021). “Apical amplification as a cellular mechanism for consciousness.” Neurosci Biobehav Rev 128: 181–195.

Pockett, S. (2002). “Difficulties with the electromagnetic field theory of consciousness.” Journal of Consciousness Studies 9(4): 51–56.

Poirazi, P. and B. W. Mel (2001). “Impact of active dendrites and structural plasticity on the memory capacity of neural tissue.” Neuron 29(3): 779–796.

Posner, M. I. and S. E. Petersen (1990). “The attention system of the human brain.” Annual Review of Neuroscience 13: 25–42.

Pribram, K. H. (1991). Brain and Perception: Holonomy and Structure in Figural Processing, Lawrence Erlbaum Associates.

Pribram, K. H., K. Yasue and M. Jibu (1991). Holonomy and dendritic microprocess. Brain and Values, Lawrence Erlbaum Associates: 71–84.

Raichle, M. E. (2015). “The brain’s default mode network.” Annual Review of Neuroscience 38: 433–447.

Ramaswamy, S. and H. Markram (2015). “Anatomy and physiology of the thick-tufted layer 5 pyramidal neuron.” Front Cell Neurosci 9: 233.

Rogers, G. L. (1950). “Gabor diffraction microscopy: the hologram as a generalized zone-plate.” Nature 166(4214): 237.

Ruiz de Miras, J., V. Costumero, V. Belloch, A. García-Vidal, A. Barros-Loscertales and C. Ávila (2019). “Complexity analysis of cortical surface detects changes in future Alzheimer’s disease converters.” Human Brain Mapping 40(1): 30–42.

Russell, S. (2019). Human Compatible: Artificial Intelligence and the Problem of Control, Viking.

Schiff, N. D. (2008). “Central thalamic contributions to arousal regulation and neurological disorders of consciousness.” Ann N Y Acad Sci 1129: 105–118.

Schiff, N. D. (2010). “Recovery of consciousness after brain injury: a mesocircuit hypothesis.” Trends Neurosci 33: 1–9.

Schneider, S. (2019). Artificial You: AI and the Future of Your Mind, Princeton University Press.

Selkoe, D. J. and J. Hardy (2016). “The amyloid hypothesis of Alzheimer’s disease at 25 years.” EMBO Molecular Medicine 8(6): 595–608.

Seth, A. K. and T. Bayne (2022). “Theories of consciousness.” Nature Reviews Neuroscience 23(7): 439–452.

Sevush, S. (2006). “Single-neuron theory of consciousness.” J Theor Biol 238(3): 704–725.

Sevush, S. (2016). The Single-Neuron Theory: Closing in on the Neural Correlate of Consciousness, Springer.

Sezgin, E., I. Levental, S. Mayor and C. Eggeling (2017). “The mystery of membrane organization: composition, regulation and roles of lipid rafts.” Nature Reviews Molecular Cell Biology 18(6): 361–374.

Shanahan, M. (2010). Embodiment and the Inner Life: Cognition and Consciousness in the Space of Possible Minds, Oxford University Press.

Shankar, G. M., S. Li, T. H. Mehta, A. Garcia-Munoz, N. E. Shepardson, I. Smith and D. J. Selkoe (2008). “Amyloid-β protein dimers isolated directly from Alzheimer’s brains impair synaptic plasticity and memory.” Nature Medicine 14(8): 837–842.

Shao, Y., B. Wang, A. T. Sornborger and L. Tao (2019). “A mechanism for synaptic copy between neural circuits.” Neural Computation 31(10): 1964–1984.

Shepherd, G. M. G. and N. Yamawaki (2021). “Untangling the cortico-thalamo-cortical loop: cellular pieces of a knotty circuit puzzle.” Nat Rev Neurosci 22(7): 389–406.

Sherman, S. M. and R. W. Guillery (2011). “Distinct functions for direct and transthalamic corticocortical connections.” J Neurophysiol 106: 1068–1077.

Sherman, S. M. and R. W. Guillery (2013). Functional Connections of Cortical Areas: A New View from the Thalamus, MIT Press.

Siskova, Z., D. Justus, H. Kaneko, D. Friedrichs, N. Henneberg, T. Beutel, J. Pitsch, S. Schoch, A. Becker, H. von der Kammer and S. Remy (2014). “Dendritic Structural Degeneration Is Functionally Linked to Cellular Hyperexcitability in a Mouse Model of Alzheimer’s Disease.” Neuron 84(5): 1023–1033.

Smith, J. H., C. Rowland, B. Harland, S. Moslehi, R. D. Montgomery, K. Schobert, W. J. Watterson, J. Dalrymple-Alford and R. P. Taylor (2021). “How neurons exploit fractal geometry to optimize their network connectivity.” Scientific Reports 11(1): 2332.

Smits, F. M., P. J. W. Pouwels, H. G. Schnack, H. E. Hulshoff Pol, R. S. Kahn, L. Bosscher, W. Post, A. H. Zwinderman, C. J. Stam, E. H. F. de Haan and H. Swaab (2016). “White matter tracts of attention: Altered fractional anisotropy in different attentional network pathways in childhood absence epilepsy.” Neuroscience 339: 106–113.

Spires-Jones, T. L. and B. T. Hyman (2014). “The intersection of amyloid beta and tau at synapses in Alzheimer’s disease.” Neuron 82(4): 756–771.

Stam, C. J., B. F. Jones, G. Nolte, M. Breakspear and P. Scheltens (2005). “Small-world networks and functional connectivity in Alzheimer’s disease.” Cerebral Cortex 17(1): 92–99.

Stapp, H. P. (1993). Mind, Matter, and Quantum Mechanics. Berlin, Springer.

Stapp, H. P. (2006). “Quantum Interactive Dualism, II: The Libet and Einstein–Podolsky–Rosen Causal Anomalies.” Erkenntnis 65(1): 117–142.

Starr, F. W., B. Hartmann and J. F. Douglas (2014). “Dynamical clustering and a mechanism for raft-like structures in a model lipid membrane.” Soft Matter 10(17): 3036–3047.

Steriade, M., E. G. Jones and D. A. McCormick (1997). Thalamus: Organisation and Function, Elsevier.

Storm, J. F., M. Boly, A. G. Casali, M. Massimini, U. Olcese, C. M. A. Pennartz and M. Wilke (2024). “An integrative, multiscale view on neural theories of consciousness.” Neuron 112(10): 1531–1552.

Suma, A., D. Sigg, S. Gallagher, G. Gonnella and V. Carnevale (2024). “Ion channels in critical membranes: Clustering, cooperativity, and memory effects.” PRX Life 2: 013007.

Suma, A., D. Sigg, A. Srivastava, G. Gonnella and V. Carnevale (2023). Channel-lipid interactions drive the emergence of ion channel clustering, channel-channel cooperativity, and activation hysteresis. 67th Annual Biophysical Society Meeting.

Susskind, L. (1995). “The world as a hologram.” Journal of Mathematical Physics 36(11): 6377–6396.

Suzuki, M. and M. E. Larkum (2020). “General Anesthesia Decouples Cortical Pyramidal Neurons.” Cell 180(4): 666–676.e613.

Tagliazucchi, E., L. Roseman, M. Kaelen, C. Orban, S. D. Muthukumaraswamy, K. Murphy and R. Carhart-Harris (2016). “Increased global functional connectivity correlates with LSD-induced ego dissolution.” Current Biology 26(8): 1043–1050.

Tegmark, M. (2000). “Importance of quantum decoherence in brain processes.” Physical Review E 61(4): 4194–4206.

Thomson, A. M. and C. Lamy (2007). “Functional maps of neocortical local circuitry.” Front Neurosci 1: 19–42.

Tong, F., M. Meng and R. Blake (2006). “Neural bases of binocular rivalry.” Trends in Cognitive Sciences 10(11): 502–511.

Tononi, G. (2004). “An information integration theory of consciousness.” BMC Neuroscience 5(1): 42.

Tononi, G. (2008). “Consciousness as integrated information: A provisional manifesto.” The Biological Bulletin 215(3): 216–242.

Tononi, G., M. Boly, M. Massimini and C. Koch (2016). “Integrated information theory: from consciousness to its physical substrate.” Nature Reviews Neuroscience 17(7): 450–461.

Tononi, G., G. M. Edelman and O. Sporns (1994). “A measure for brain complexity: relating functional segregation and integration in the nervous system.” Proceedings of the National Academy of Sciences USA 91: 5033–5037.

Varela, F. J. (1996). “Neurophenomenology: A methodological remedy for the hard problem.” Journal of Consciousness Studies 3(4): 330–349.

Varela, F. J., E. Thompson and E. Rosch (1991). The Embodied Mind: Cognitive Science and Human Experience. Cambridge, MA, MIT Press.

Varley, T. F., R. Carhart-Harris, L. Roseman, D. K. Menon and E. A. Stamatakis (2020). “Serotonergic psychedelics LSD & psilocybin increase the fractal dimension of cortical brain activity in spatial and temporal domains.” NeuroImage 220: 117049.

Wang, Z., A. C. Bovik, H. R. Sheikh and E. P. Simoncelli (2004). “Image quality assessment: from error measurement to structural similarity.” IEEE Transactions on Image Processing 13(4): 600–612.

Waters, J. P. (1966). “Holographic image synthesis utilizing theoretical methods.” Applied Physics Letters 9(11): 405–407.

Weiner, V. S., D. W. Zhou, P. Kahali, E. P. Stephen, R. A. Peterfreund, L. S. Aglio, M. D. Szabo, E. N. Eskandar, P. L. Purdon and E. N. Brown (2023). “Propofol disrupts alpha dynamics in functionally distinct thalamocortical networks during loss of consciousness.” Proceedings of the National Academy of Sciences 120(11): e2207831120.

Wen, X.-G. (1995). “Topological orders and edge excitations in fractional quantum Hall states.” Advances in Physics 44(5): 405–473.

Werner, G. (2010). “Fractals in the nervous system: conceptual implications for theoretical neuroscience.” Frontiers in Physiology 1: 15.

Whyte, C. J., E. J. Muller, J. Aru, M. Larkum, Y. John, B. R. Munn and J. M. Shine (2025). “A burst-dependent thalamocortical substrate for perceptual awareness.” PLoS Comput Biol 21(4): e1012951.

Williams, S. R. and G. J. Stuart (2002). “Dependence of EPSP efficacy on synapse location in neocortical pyramidal neurons.” Science 295(5561): 1907–1910.

Zaletel, I., R. Filipović and N. Puškaš (2015). “Fractal dimension of apical dendritic arborization differs in the superficial and the deep pyramidal neurons of the rat cerebral neocortex.” Journal of Theoretical Biology 369: 1–9.

Zheng, J. and M. Meister (2025). “The unbearable slowness of being: Why do we live at 10 bits/s?” Neuron 113(2): 192–204.

